# Transient cytoskeletal anisotropy encodes short-term mechanical memory

**DOI:** 10.64898/2026.03.09.710573

**Authors:** Clara Gomez-Cruz, Matthieu Gelin, Lucas Pradeau-Phélut, Arrate Muñoz-Barrutia, Sandrine Etienne-Manneville, Daniel Garcia-Gonzalez

**Affiliations:** Department of Continuum Mechanics and Structural Analysis, Universidad Carlos III de Madrid, Avda. de la Universidad 30, 28911 Leganés, Madrid, Spain; 60Nd S.L., Avda. Gregorio Peces Barba 1, 28919 Leganés, Madrid, Spain; Cell Polarity, Migration and Cancer Unit, Institut Pasteur, UMR3691 CNRS, Universite de Paris, Equipe Labellisee Ligue Contre le Cancer 2023, Paris F-75015, France; Departamento de Neurociencia y Ciencias Biomédicas, Universidad Carlos III de Madrid, Avda. de la Universidad 30, 28911 Leganés, Madrid, Spain; Área de Ingeniería Biomédica, Instituto de Investigación Sanitaria Gregorio Marañón, Calle del Doctor Esquerdo 46, Madrid ES28007, Spain

**Keywords:** Glioblastoma, Mechanical memory, Cytoskeletal remodeling, Continuum modeling, Mechanobiology

## Abstract

Cells can experience time-varying mechanical cues, particularly when navigating through changing and complex microenvironments. Yet whether and how cells retain and use a short-term mechanical memory of recent deformations remains unclear. Here we show that, in glioblastoma cells, this memory is encoded by transient cytoskeletal anisotropy. Using uniaxial magneto-mechanical actuation aligned or perpendicular to the cell long axis, nanoindentation, and selective cytoskeletal perturbations, we find that distinct architectures of the actin cytoskeleton drive opposite mechanical responses: actin stress fibers mediate stiffening under stretch, whereas the actin cortex underlies softening under perpendicular loading. Vimentin intermediate filaments are essential to stabilize actin organization under load, preserving deformation-specific mechanics. Quantitative imaging reveals that mechanical actuation induces network-specific alignment and anisotropy, stronger for actin than vimentin, that persists transiently after unloading and bias subsequent responses, revealing a short-lived, deformation-dependent mechanical memory. To integrate these observations, we develop a multi-network constitutive model that links cytoskeletal architecture and loading history to cell-scale mechanics, reproducing both the asymmetric mechanical responses and the measured reorganization dynamics. These findings provide a structural basis for short-term mechanical memory and suggest how cancer cells could exploit residual anisotropy to adapt to fluctuating solid stresses and confinement, transiently biasing polarization, force transmission, and directional persistence during invasion. They also identify vimentin-actin coupling and the kinetics of cytoskeletal remodeling as potential levers to limit the mechanical adaptability of invasive cancer cells.

## 1. Introduction

Cells operate in mechanical environments that evolve continuously during development, tissue repair, and disease. Embryonic morphogenesis, wound healing, and fibrotic remodeling all involve dynamic changes in tissue tension, compression, and stiffness, exposing cells to complex sequences of mechanical deformation rather than static cues [1, 2, 3]. Across these contexts, mechanical cues are not static inputs but dynamic signals that unfold over time, requiring cells to sense, integrate, and adapt to sequences of mechanical deformations rather than isolated stimuli [4, 5]. Cancer progression represents an extreme manifestation of this mechanical variability. As tumors grow and invade surrounding tissue, cancer cells encounter spatially and temporally heterogeneous mechanical environments shaped by solid stress, confinement, altered extracellular matrix architecture, and interactions with neighboring cells [6, 7]. In highly invasive cancers such as glioblastoma, tumor cells migrate through mechanically distinct regions of the brain, including perivascular niches, white matter tracts, and densely packed tumor cores, experiencing alternating phases of compression, relaxation, and tensile deformation [8, 9, 10].

A growing body of work indicates that mechanosensing and mechanoresponse unfold across broad timescales, from rapid cytoskeletal adjustments to slower nuclear signaling events, and depend not only on the instantaneous magnitude or mode of loading, but also on the temporal characteristics of mechanical stimulation, including duration, rate, and sequence. [4, 11, 12]. In this context, the concept of mechanical memory has emerged, describing situations in which past mechanical environments influence future cellular behavior even after the original stimulus has been removed [13]. Most studies of mechanical memory have focused on long-term effects mediated by transcriptional reprogramming, epigenetic modifications, or stable changes in cell state, particularly in cancer and fibrosis [14, 7, 3]. However, migrating cells often operate on much shorter timescales (minutes to hours) raising the possibility that memory could also be encoded through faster, purely structural mechanisms [15].

The cytoskeleton is a prime candidate for such short-term mechanical memory. As the primary load-bearing architecture of the cell, cytoskeletal networks continuously assemble, disassemble, and reorganize in response to mechanical cues, thereby introducing intrinsic timescales into cellular mechanics [16, 17, 18]. Actin filaments, intermediate filaments, and microtubules have well-established but distinct roles in force generation, mechanical stability, and migration [19, 20, 21, 22]. In particular, actin stress fibers support contractility and directional force transmission, while the actin cortex regulates cell shape and resistance to deformation, and vimentin intermediate filaments contribute to mechanical resilience and invasive potential in cancer cells [8, 14]. Mechanical loading can induce cytoskeletal alignment and anisotropy, which are closely linked to cell polarity, persistent migration, and directional force generation [23, 24, 25, 18]. Whether such mechanically-induced organization can persist after unloading and bias subsequent mechanical responses remains largely unexplored.

Understanding how short-term mechanical memory arises requires dissecting how individual cytoskeletal components contribute to cellular mechanics under different deformation modes and how their interactions evolve over time [26]. While actin and vimentin networks are known to be structurally interdependent [27], with vimentin stabilizing actin organization and enhancing resistance to mechanical stress, their interaction has mainly been studied under static or simplified loading conditions [28, 26]. How cytoskeletal networks respond to temporally complex, sequential deformations and whether transient cytoskeletal anisotropy can act as a memory of prior mechanical states remain open questions [29].

Recent computational and theoretical models have begun to capture key features of cytoskeletal mechanics, including nonlinear stiffening, anisotropy, and active remodeling [29, 30, 31]. However, most existing frameworks focus on steady-state behavior or isolated loading protocols and do not explicitly address how deformation history shapes transient cytoskeletal organization and short-term mechanical memory [32]. Bridging this gap requires integrative approaches that combine controlled mechanical perturbations, targeted cytoskeletal manipulations, and mechanistically grounded models capable of linking cytoskeletal architecture to time-dependent mechanical responses.

Here, we address these questions by combining controlled magneto-mechanical actuation, selective cytoskeletal perturbations, and a mechanistically grounded constitutive model to investigate how glioblastoma cells encode and retain short-term mechanical information. We show that transient cytoskeletal anisotropy, arising from deformation-dependent reorganization of actin and its stabilization by vimentin, underlies short-term mechanical memory in living cells. By explicitly linking cytoskeletal structure, deformation mode, and mechanical history, our work establishes a structural basis for short-term mechanical memory and provides a framework for understanding how invasive cancer cells adapt to dynamically evolving mechanical environments.

## 2. Results

### 2.1. Actin-driven deformation-dependent stiffening requires vimentin-mediated structural stabilization

To dissect how cytoskeletal composition and mechanical state jointly regulate cellular mechanics, we combined selective cytoskeletal perturbations with mechanical actuation controlling tensile and compressive deformation modes. We focused on four major cytoskeletal components: branched actin forming the actin cortex, actin stress fibers, vimentin intermediate filaments and microtubules, and assessed their respective contributions to basal and deformation-induced cell stiffness (Fig. 1a).

**Figure 1.**
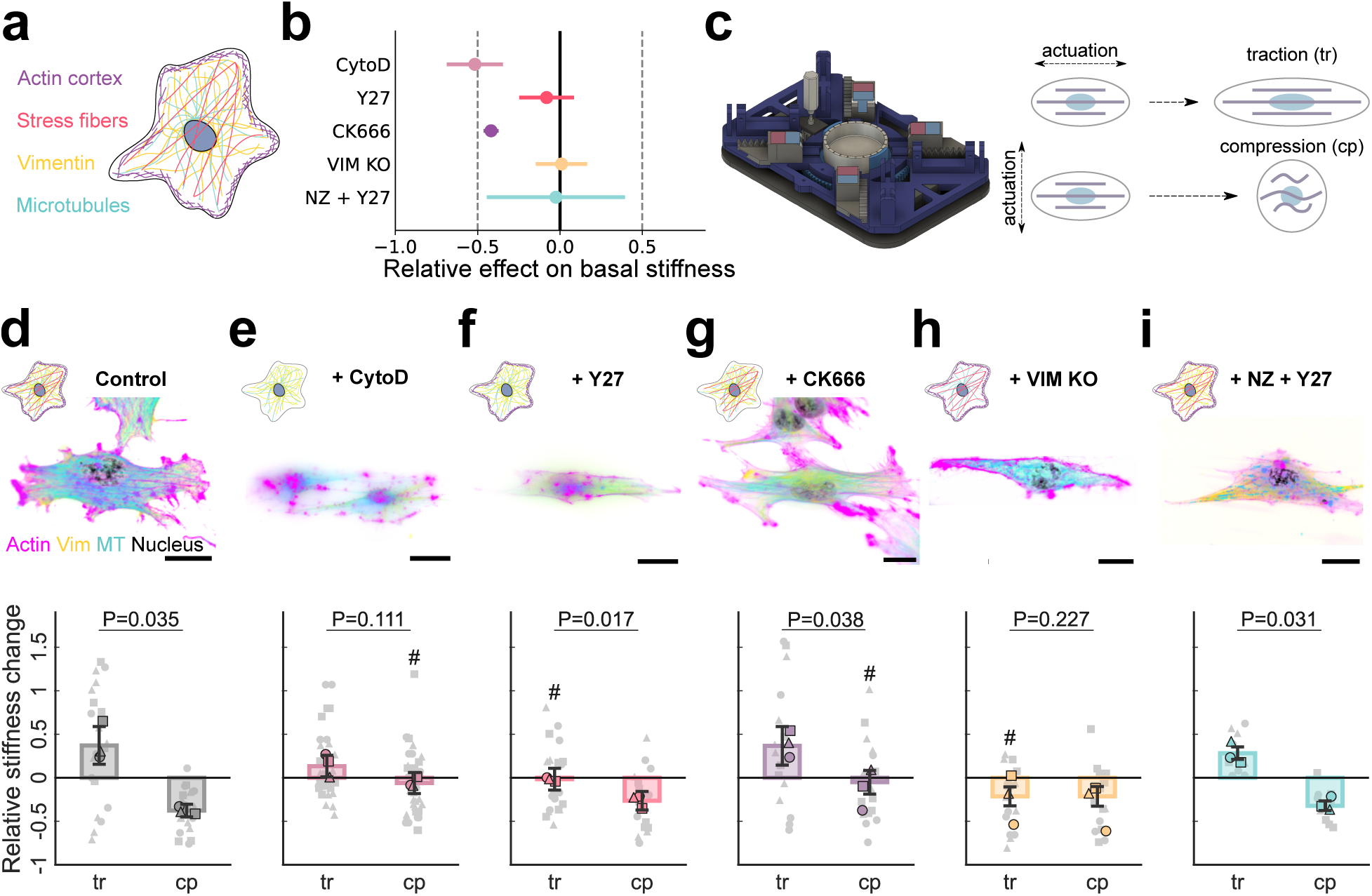
Cytoskeletal interplay governs cell mechanics under deformation. **a** Schematic representation of the main elements of the glioblastoma cytoskeleton: actin cortex, actin stress fibers, vimentin intermediate filaments, and microtubules. **b** Relative fold changes in relaxed (non-actuated) cell stiffness of selective cytoskeletal perturbations measured relative to control cells. **c** Schematic of the magneto-mechanical actuation platform and loading configurations. Actuation parallel to the cell long axis results in tensile stretch (traction), whereas actuation perpendicular to this axis results in compression. **d-i** Representative actin (magenta), vimentin (yellow) and microtubule (cyan) organization in actuated glioblastoma cells and relative stiffness change between relaxed and actuation states for cells under traction and compression in **d** control conditions, cells subjected to **e** 1 *µ*M Cytochalasin D, **f** 10 *µ*M Y-27632 or **g** 50 *µ*M Ck-666, **h** vimentin KO cells, and cells subjected to **i**10 *µ*M Nocodazole + 1 *µ*M Y-27632. Scale bars: 20 *µm*. n=40-70 cells analyzed per condition from N=3 independent nanoindentation assays. Data are presented as mean *±* SEM. A two-tailed unpaired t-test with Welch’s correction was used for statistical analysis. P-values shown in the graphs compare relative stiffness changes between cells under traction (tr) and compression (cp) for each condition. Comparisons with the corresponding control case for each orientation are marked by hash symbols (#), with ns p > 0.05 (not shown), # p*≤*0.05.

We first quantified the effect of selectively disrupting individual cytoskeletal networks on basal stiffness, measured in the relaxed, non-actuated state by nanoindentation (Fig. 1b and Supplementary Fig. S1). Disruption of the actin cytoskeleton using Cytochalasin D, which inhibits actin polymerization, resulted in a pronounced reduction of basal stiffness by 50%, establishing actin as the primary determinant of baseline mechanical resistance. Inhibition of ROCK-dependent actomyosin contractility using Y-27632 led to a much smaller decrease in basal stiffness (5%), indicating that stress fiber–associated contractility contributes only modestly to the relaxed mechanical state. In contrast, inhibition of Arp2/3-dependent branched actin polymerization using CK666 caused a substantial softening comparable to Cytochalasin D (50%), consistent with a dominant contribution of branched cortical actin networks to basal stiffness. To limit the effect of increased actomyosin contractility associated with nocodazole treatment, a low dose of Y27632 was added to the microtubule disruption condition. In addition, complementary assays were performed using nocodazole alone. Both assays showed no measurable change in basal stiffness following microtubule depolymerization (Fig. 1b and Supplementary Fig. S2). Strikingly, vimentin knockout did not produce a measurable change in basal stiffness either, indicating that this network does not significantly contribute to the cell’s relaxed cortical mechanical response, although vimentin depletion has been reported to reduce global whole-cell stiffness [33].

We next probed how these cytoskeletal components regulate stiffness under active mechanical loading. Cells were subjected to controlled uniaxial magneto-mechanical actuation using an upgraded version of the Neo-Mag device (60Nd S.L.) [34], compatible with inverted microscopy and nanoindentation (Fig. 1c). Two loading configurations were applied: actuation parallel to the cell’s long axis, resulting in uniaxial stretching, and actuation perpendicular to this axis, resulting in the compression of the long axis. As previously demonstrated using the same platform and 30% actuation amplitude, perpendicular substrate stretch generates compressive strain along the cell major axis at the cellular scale [34]. These two deformation modes therefore decouple directionally coherent tensile loading from compressive deformation.

In control cells, uniaxial stretch induced a robust stiffening response, with stiffness increasing by 37% relative to the relaxed state, whereas uniaxial compression led to a significant softening (38% decrease) (Fig. 1d). This asymmetric mechanical response demonstrates that cell stiffness is not only deformation-dependent but also mode-specific, with cells actively reinforcing their mechanics under tensile loading while yielding under compression. These results are consistent with our prior findings in primary astrocytes [34]. Disrupting the actin cytoskeleton abolished this deformation-dependent behavior. Following Cytochalasin D treatment, cells failed to stiffen under stretch and did not soften under compression, remaining mechanically indistinguishable from the relaxed state in both loading modes (Fig. 1e). This complete loss of mechanical adaptability establishes actin as a major driver of deformation-dependent stiffening and softening. Pharmacological perturbation of actin sub-networks revealed distinct functional roles under different deformation modes. Inhibition of ROCK-dependent actomyosin contractility using Y-27632 suppressed stretch-induced stiffening, while compression still produced a marked softening response (Fig. 1f). In contrast, inhibition of Arp2/3-dependent branched actin polymerization using CK666 preserved stretch-induced stiffening but abolished compression-induced softening (Fig. 1g). These differential responses show that distinct actin architectures mediate mechanically asymmetric behaviors through fundamentally different mechanisms. In particular, stretch-induced stiffening appears to rely on stress fiber–associated contractile actomyosin structures that bear and progressively reinforce tensile loads, consistent with their aligned, tension-dominated organization and known strain-stiffening behavior [35, 34]. By contrast, the pronounced softening observed under compression points to a distinct contribution of branched cortical actin networks, which forms a thin, shell-like network at the cell periphery and may lose effective load-bearing capacity under compressive or in-plane contractile states, for instance through buckling or wrinkling instabilities [36, 37, 38]. Together, these results suggest that actin stress fibers actively drive stiffening under uniaxial stretch, whereas the actin cortex governs the cell’s ability to soften under compression.

In contrast to actin-specific perturbations, vimentin-deficient cells exhibited pronounced mechanical softening under both uniaxial stretch and compression, with stiffness decreasing by 22% in both conditions (Fig. 1h). In these cells, fluorescence imaging revealed a marked disruption of the actin cytoskeleton under deformation, indicating that the observed softening arises from a failure to preserve actin network organization rather than due to the lack of vimentin directly. Thus, although actin-driven mechanisms are responsible for deformation-dependent stiffening, it appears that vimentin is required to mechanically stabilize the actin cytoskeleton under load. By contrast, microtubule-depleted cells retained the full deformation-dependent mechanical response observed in controls, stiffening under stretch and softening under compression (Fig. 1i), further confirming that microtubules do not play a dominant role in regulating cell stiffness under either basal or the actuated conditions in this study. Additional experiments on cells at different actuation angles were performed, see Supplementary Fig. S3.

Together, these results establish a hierarchical mechanical organization of the cytoskeleton in which actin networks actively drive deformation-dependent mechanical responses, while vimentin is required to stabilize actin organization and preserve these responses under load. Stress fiber-associated contractile structures and branched cortical actin networks contribute preferentially to tensile stiffening and compressive softening, respectively, whereas vimentin ensures the structural integrity of actin architectures during deformation. In the absence of vimentin, actin-driven mechanics collapse, leading to global mechanical softening irrespective of deformation mode. These findings indicate that deformation-dependent cell mechanics do not arise solely from the presence of specific cytoskeletal components, but from their mechanically coupled organization, motivating a structural analysis of cytoskeletal reorganization and anisotropy as the origin of deformation mode–specific mechanics in the following section.

### 2.2. Cytoskeletal reorganization and anisotropy provide the structural origin of deformation mode–specific mechanics

To identify the structural mechanisms underlying deformation mode-dependent mechanical responses, we quantified the organization, orientation, and anisotropy of actin and vimentin networks under relaxed and traction-actuated conditions. Cytoskeletal organization was extracted from fluorescence microscopy images by resolving individual filament orientations and color-coding fibers according to their angular direction (Fig. 2a). This approach enabled a quantitative comparison of fiber alignment relative to the imposed uniaxial deformation axis, as well as the computation of network anisotropy and principal orientation. Because microtubule disruption had no measurable effect on mechanical response under the loading conditions considered here (Fig 1i; Supplementary Fig. S2), microtubules were not included in this structural analysis.

**Figure 2.**
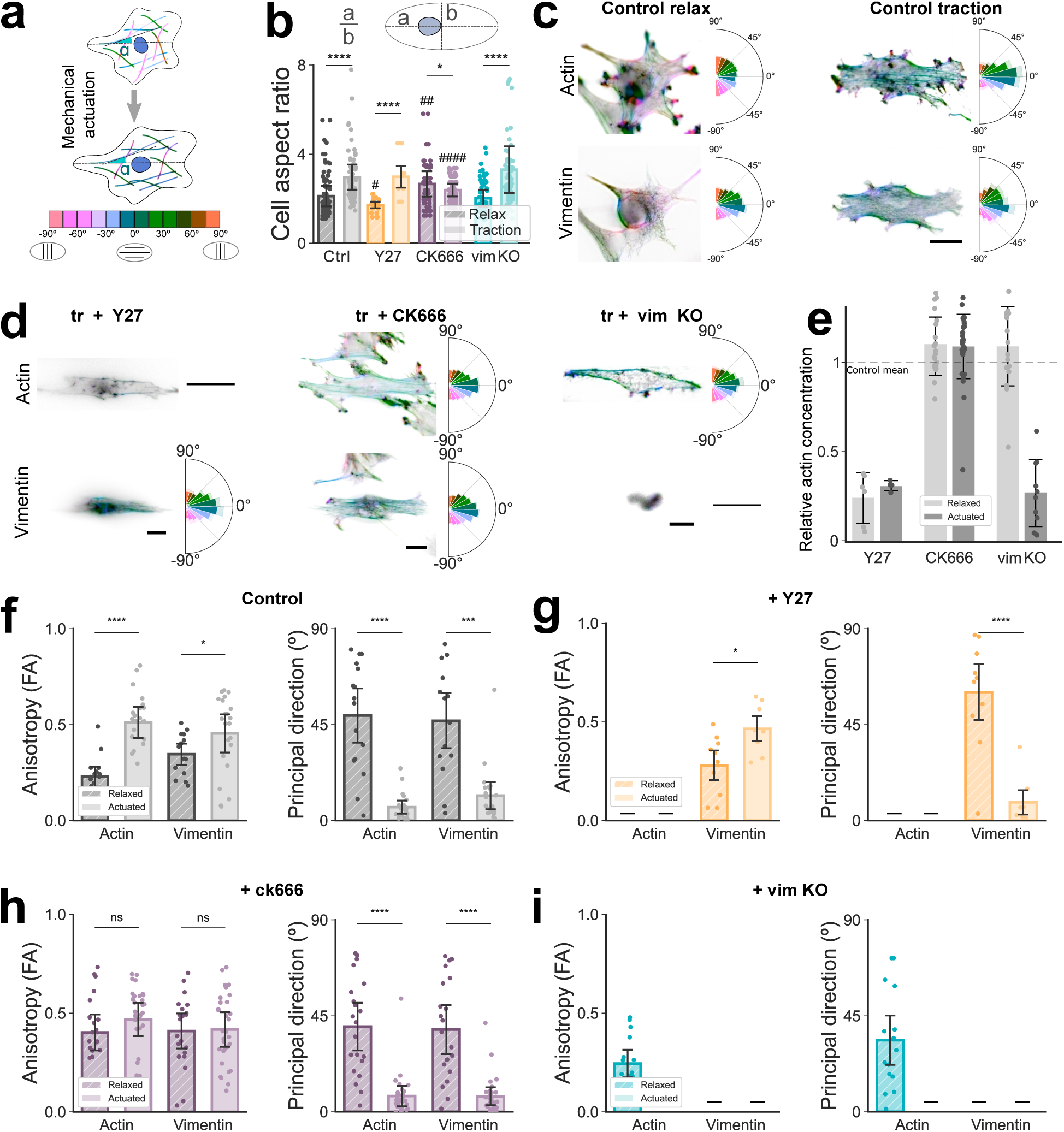
Cytoskeletal reorganization and anisotropy underlie deformation-dependent cell mechanics. **a** Schematic illustration of the quantification of cytoskeletal organization, showing fiber orientation maps color-coded by direction and classification of fibers aligned parallel or perpendicular to the imposed uniaxial deformation axis. **b** Cell aspect ratio, reflecting global cell elongation, measured in relaxed and actuated conditions for control cells and cells subjected to 10 *µ*M Y-27632, 50 *µ*M CK-666, or vimentin knockout. **c** Representative actin and vimentin organization in control cells under relaxed and traction conditions, colorcoded by fiber orientation, showing isotropic organization in the relaxed state and deformation-induced anisotropy upon actuation. **d** Representative actin and vimentin organization under traction for cells subjected to 10 *µ*M Y-27632, 50 *µ*M CK-666, or vimentin KO. **e** Relative actin and vimentin content (total fluorescence intensity) normalized to control cells across perturbations. **f-i** Quantification of anisotropy (fractional anisotropy, FA) and principal direction of actin and vimentin networks in relaxed and actuated conditions for **f** control cells, **g** Y-27632–treated cells, **h** CK-666–treated cells, and **i** vimentin KO cells, showing deformation-induced alignment and network-specific anisotropy evolution across cytoskeletal perturbations. Histograms represent the average fiber distribution for each condition. Scale bars: 20 *µ*m. n=10-20 cells analyzed per condition from N=3 independent assays. Data are presented as mean *±* SEM. A two-tailed unpaired t-test with Welch’s correction was used for statistical analysis. Comparisons with the corresponding relaxed condition for each network are marked by hash symbols (#), with ns p > 0.05, # p*≤*0.05.

We first verified that the applied actuation effectively deforms cells across conditions by quantifying global cell shape changes. Cell aspect ratios were measured under relaxed and actuated states for all perturbations (Fig. 2b). In control cells, as well as in Y27632-treated and vimentin knockout cells, tensile actuation induced a significant increase in cell aspect ratio (40% increase), confirming elongation along the loading direction. CK666-treated cells exhibited a high aspect ratio already in the relaxed state, which remained largely unchanged upon actuation, indicating that these cells were pre-elongated prior to loading. Importantly, these measurements confirm that actuation induces comparable global deformation across conditions, and that the differences in cytoskeletal organization described below cannot be attributed to the absence of cell elongation, but instead reflect network-specific remodeling responses.

We next analyzed the spatial organization of actin and vimentin networks in cells under non-actuated conditions (Fig. 2c and Supplementary Fig. S4). In control cells, both actin and vimentin fibers displayed on average largely isotropic orientation distributions, with no dominant preferential direction. Upon uniaxial stretch, actin fibers reorganized into a highly anisotropic architecture aligned with the loading direction, while vimentin also aligned but to a lesser extent. Quantitatively, actin fractional anisotropy increased by 0.24, compared to a smaller increase of 0.11 for vimentin (Fig. 2f), and the principal direction of both networks rotated toward the actuation axis, with a stronger directional bias for actin. These results establish that deformation induces coordinated but non-identical reorganization of actin and vimentin, consistent with their distinct mechanical roles identified in Fig. 1. Selective cytoskeletal perturbations revealed how this reorganization depends on network integrity (Fig. 2d and Supplementary Fig. S5). In Y27632-treated cells, actin stress fibers were absent, resulting in a weak and poorly organized actin signal. Despite this, the vimentin network developed a pronounced anisotropy and aligned along the direction of actuation, closely resembling the behavior observed in control cells. In CK666-treated cells, both actin and vimentin networks exhibited partial anisotropy, but to a lesser degree than in controls. Notably, in vimentin knockout cells, vimentin was absent and the actin network appeared severely disrupted under actuation, with few discernible fibers and no coherent alignment.

To establish in which conditions cytoskeletal alignment can be meaningfully quantified, we first assessed the relative abundance of actin and vimentin networks across perturbations in relaxed non-actuated conditions (Fig. 2e). CK666 treatment did not significantly alter the overall abundance of either network. Y27632-treated cells exhibited a strong reduction in actin content while maintaining vimentin levels comparable to control, whereas vimentin knockout cells retained actin levels similar to control in relaxed conditions and lacked vimentin, as expected. These measurements indicate that, depending on the perturbation, one or both filamentous networks may be partially depleted or structurally compromised, constraining the interpretation of alignment and anisotropy metrics. Accordingly, quantitative analyses of fiber orientation and anisotropy were interpreted only in conditions where a sufficiently organized filament network was present. Within this framework, quantitative analysis of anisotropy and principal direction revealed distinct, network-specific reorganization responses to actuation (Fig. 2f–i). In control cells, both actin and vimentin exhibited a significant increase in anisotropy upon actuation, accompanied by alignment along the loading axis, with a more pronounced response for actin (Fig. 2f). In Y27632-treated cells, actin anisotropy remained negligible due to the absence of organized actin fibers, whereas the vimentin network developed anisotropy and aligned with the actuation direction similarly to control cells (Fig. 2g). In CK666-treated cells, both actin and vimentin networks displayed elevated anisotropy already in the relaxed state, which did not increase further upon actuation. Nevertheless, both networks reoriented their principal direction toward the loading axis (Fig. 2h), indicating reorientation without an additional increase in anisotropy. Finally, in vimentin knockout cells, actin organization under relaxed conditions was comparable to control, but actuation failed to induce increased anisotropy or stable alignment due to disruption of the actin network under loading (Fig. 2i).

Together, these results demonstrate that deformation mode–dependent mechanical behavior arises from active, network-specific cytoskeletal reorganization. Actin undergoes pronounced alignment and anisotropy development under uniaxial stretch, directly correlating with deformation-induced stiffening. In contrast, vimentin aligns more weakly but plays a critical role in regulating actomyosin stability and stress fiber organization under tensile load. This role is complementary to previous studies showing that vimentin also supports cortical actin organization and mechanics [39]. Perturbations that preserve actin-driven force generation but compromise its structural stabilization disrupt anisotropic remodeling and, consequently, mechanical response. These findings establish cytoskeletal anisotropy as the structural origin of deformation-dependent mechanics and provide the mechanistic foundation for the constitutive modeling framework introduced next.

### 2.3. A multi-network constitutive model reproduces actin-driven stiffening and vimentin-stabilized anisotropy

The experimental results presented above reveal that deformation-dependent cell mechanics arise from a tightly coupled interplay between cytoskeletal structure, network-specific mechanics and active remodeling. While these experiments identify clear functional roles for actin stress fibers, actin cortex and vimentin intermediate filaments, they also raise fundamental mechanistic questions that cannot be resolved from experiments alone. In particular, they motivate the need for a unified framework capable of integrating cytoskeletal architecture, anisotropy and deformation history into a predictive description of cellular mechanics. This framework is necessary to test the mechanisms suggested by the data, to separate the specific contributions of each cytoskeletal network, and to understand how their interactions produce the observed mechanical responses. To address this need, we developed a continuum constitutive model that explicitly links cytoskeletal organization to deformation-dependent cell mechanics. The model is grounded on the experimentally observed hierarchy of cytoskeletal contributions and is designed to serve as a quantitative testbed for the hypotheses summarized in Table 1. Within this framework, the cell is described as a continuum whose mechanical response emerges from the interaction between an isotropic background contribution, representing the cytosolic matrix and unresolved components, and anisotropic contributions associated with actin stress fibers, actin cortex, and vimentin intermediate filaments (Fig. 3a). All cytoskeletal networks are assumed to experience the same macroscopic deformation, while contributing through distinct constitutive responses and evolving internal microstructural states.

**Figure 3.**
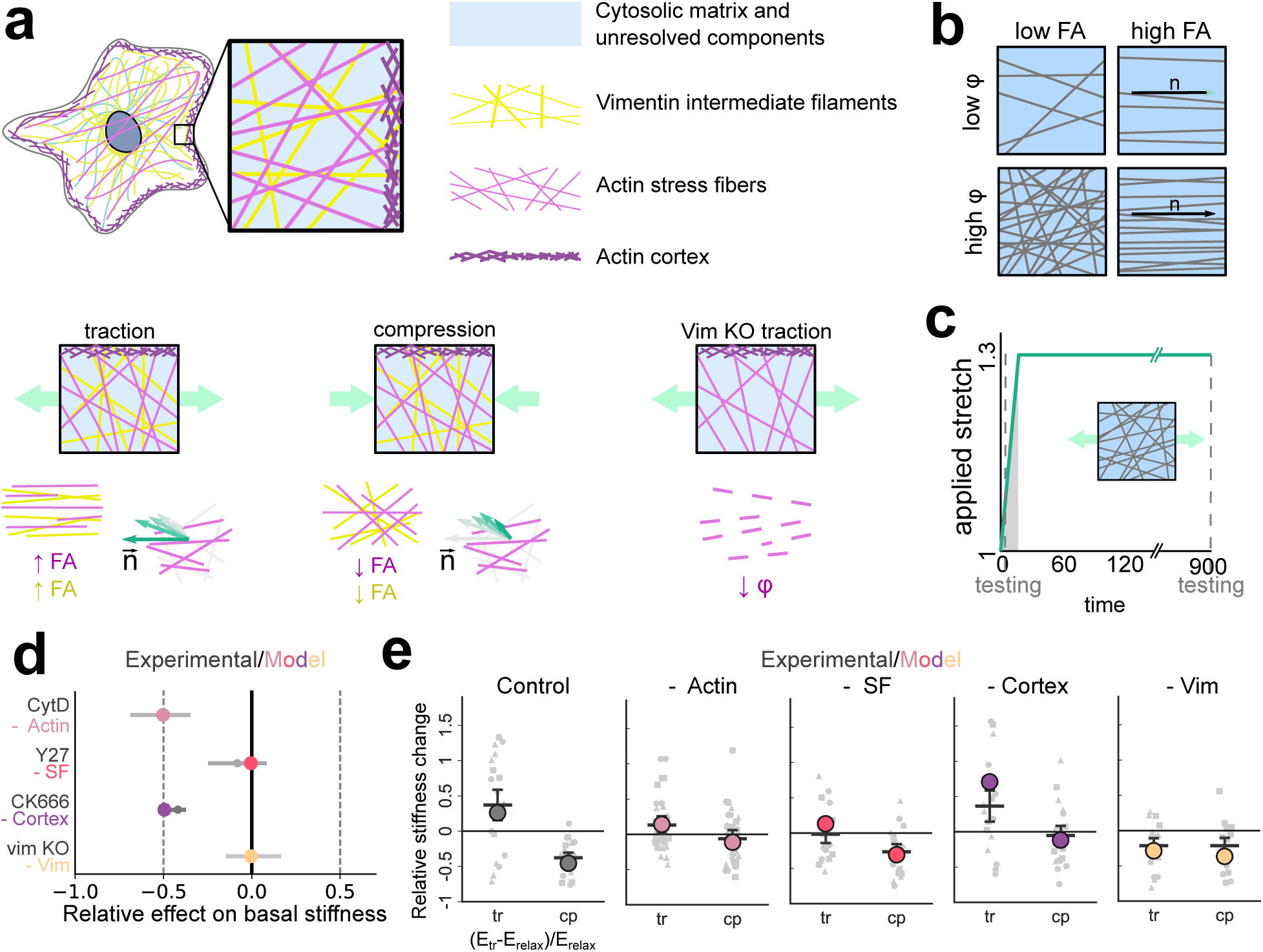
A continuum constitutive model links cytoskeletal anisotropy to deformation-dependent cell mechanics. **a** Schematic overview of the multi-network constitutive framework, in which the cell is modeled as a composite continuum whose mechanical response emerges from the additive contributions of an isotropic background, actin stress fibers, actin cortex, and vimentin intermediate filaments. The model incorporates deformation-driven evolution of fiber anisotropy, preferred orientation, and actin-vimentin coupling. **b** Illustration of low and high fractional anisotropy (FA) states related to fiber alignment, and illustration of low and high fibrosity. **c** Applied stretch protocol used for simulations, consisting of a rapid loading ramp followed by a sustained deformation period, matching the experimental actuation conditions. **d** Comparison between experimentally measured (gray) and model-predicted (colored) relative effects of selective cytoskeletal perturbations on basal cell stiffness, showing quantitative agreement across conditions. **e** Experimental (gray) and model-predicted (colored) relative stiffness changes between traction (tr) and compression (cp) for control cells and for selective removal or inhibition of individual cytoskeletal networks (actin complete network, actin stress fibers, actin cortex and vimentin), demonstrating that the model reproduces actin-driven stiffening under traction and vimentin-dependent stabilization of deformation-induced mechanical responses.

**Table 1:** Main constitutive hypotheses of the cytoskeletal model. The model describes glioblastoma cells as a continuum whose mechanical response and adaptation emerge from the interaction between isotropic cellular components, actin networks, and vimentin intermediate filaments.

| **Component / concept** | **Hypothesis** | **Mechanical implication** |
| --- | --- | --- |
| Composite cytoskeletal architecture | The cell behaves as a composite continuum with additive energetic contributions from an isotropic background, actin networks, and vimentin intermediate filaments. | Total cellular stress and stiffness emerge from the superposition of multiple cytoskeletal phases. |
| Baseline isotropic response | Cytosol and unresolved cellular components behave as an incompressible, non-linear isotropic material. | Provides a deformation-dependent baseline stiffness independent of cytoskeletal alignment. |
| Deformation-driven remodeling | Cytoskeletal reorganization is driven by principal stretch components and deformation mode. | Structural alignment follows principal stretches and loading mode (e.g.: uniaxial, biaxial, shear, etc). |
| Tension-dominated filament mechanics | Actin stress fibers and vimentin intermediate filaments transmit mechanical load primarily in tension and buckle under compression. | Cellular stiffness is direction- and sign-dependent. |
| Turnover-enabled stress relaxation | Stress fibers undergo continuous turnover, with newly polymerized filaments incorporated in a stress-free state. | Enables stress relaxation and adaptation under sustained deformation. |
| Stretch-activated actin assembly | Stress-fiber polymerization is activated only beyond a critical tensile stretch threshold. | Couples mechanical loading history to actin network growth. |
| Vimentin-guided actin organization | Vimentin intermediate filaments stabilize actin networks and guide stress-fiber alignment. | Loss of vimentin weakens actin integrity and reduces cellular stiffness under tension. |
| Structural memory | Cytoskeletal reorganization is encoded in evolving internal microstructural variables that persist beyond unloading. | Gives rise to history-dependent mechanics and mechanical memory. |

A central feature of the model is the explicit incorporation of cytoskeletal anisotropy and reorganization as state variables. In particular, the model distinguishes between the degree of network alignment, quantified by a fractional anisotropy (FA) parameter, and the effective load-bearing fraction of polymerized filaments (fibrosity), which together control the magnitude and directionality of cytoskeletal reinforcement (Fig. 3b). Network-specific descriptors capture the degree of alignment, preferred orientation, and effective load-bearing fraction of each filamentous network, enabling a direct structural link between the cytoskeletal organization quantified in Fig. 2 and the mechanical response measured by nanoindentation in Fig. 1. In this formulation and motivated by the experimental observations, actin stress fibers and vimentin intermediate filaments are treated as tension-dominated networks that progressively reinforce under stretch, whereas the actin cortex is modeled as an effective isotropic shell-like contribution whose mechanical role is strongly reduced under compressive deformation. Coupling between actin and vimentin is incorporated to reflect the experimentally observed stabilizing role of vimentin on actin organization under load. A detailed description of the constitutive equations, internal variables, and evolution laws is provided in Methods and Supplementary Information.

Model simulations were performed under loading protocols that mirror the experimental actuation conditions, including a rapid stretch ramp followed by sustained deformation (Fig. 3c). Model parameters were identified using a combination of experimental measurements reported in this study and values drawn from the literature, ensuring that each contribution operates within biologically relevant ranges (Supplementary Table S1). Without further fitting to the mechanical actuation data, the model accurately reproduces the relative effects of selective cytoskeletal perturbations on basal cell stiffness (Fig. 3d). In addition, it captures the asymmetric mechanical response observed under active loading, predicting actin-driven stiffening under uniaxial traction and softening under compression, as well as the collapse of deformation-dependent mechanics upon vimentin removal (Fig. 3e). Further parametric analyses are collected in Supplementary Fig. S6. These results demonstrate that the proposed framework quantitatively links cytoskeletal composition and organization to deformation-dependent cell mechanics across distinct loading modes and perturbation conditions.

Beyond reproducing the experimental observations reported here, the constitutive model provides a predictive framework for exploring how targeted alterations of specific cytoskeletal components, such as pharmacological inhibition or genetic depletion, affect cellular mechanical behavior under different mechanical environments. This predictive capability is particularly relevant in the context of glioblastoma progression, where cytoskeletal remodeling and mechanical adaptability are tightly linked to invasive and migratory phenotypes [40, 41]. Importantly, the formulation naturally extends to time-dependent remodeling processes, allowing the evolution of cytoskeletal organization to be explicitly coupled to mechanical loading history. Next, we leverage this capability to investigate how the model predicts the dynamics of cytoskeletal reorganization under sustained deformation and perturbation conditions.

### 2.4. Model-guided analysis predicts cytoskeletal alignment dynamics and responses to biological perturbations

Having established that the constitutive framework quantitatively reproduces deformation-dependent changes in cell stiffness across cytoskeletal perturbations, we next asked whether the same model, without any additional fitting or parameter recalibration, can also predict the structural dynamics of cytoskeletal reorganization observed experimentally. This mirrors the experimental progression presented above, where deformation-induced mechanical responses were first identified at the level of cell stiffness and subsequently traced back to network-specific cytoskeletal anisotropy. Here, we leverage the model as a predictive tool to assess whether capturing mechanical responses is sufficient to also recover the temporal evolution of cytoskeletal organization under sustained deformation.

Model simulations were performed under stretch protocols matching the experimental conditions, consisting of a rapid loading ramp followed by sustained deformation (Fig. 4a). Starting from different initial network configurations, the model predicts a progressive, time-dependent evolution of both actin and vimentin organization, characterized by changes in network orientation and anisotropy (Fig. 4b). In agreement with experimental observations, actin networks exhibit a rapid increase in anisotropy and align efficiently with the principal stretch direction, whereas vimentin reorganizes more gradually, displaying slower alignment kinetics and more moderate anisotropy development. These distinct temporal responses emerge naturally from the network-specific remodeling mechanisms encoded in the model and reflect the different structural roles of actin and vimentin identified experimentally. Quantitative comparison with experimental measurements confirms that the model accurately predicts the evolution of cytoskeletal orientation and anisotropy without further adjustment of parameters. Both vimentin (Fig. 4c) and actin (Fig. 4d) networks progressively rotate toward the direction of applied stretch, with alignment dynamics and anisotropy growth that depend on the initial network orientation and degree of organization. Importantly, the model captures not only the steadystate aligned configurations but also the transient pathways by which these states are reached, demonstrating that the observed cytoskeletal reorganization arises from continuous remodeling rather than instantaneous structural realignment. The predictive capability of the framework further extends to biologically perturbed conditions. In simulations mimicking vimentin depletion, the model predicts a destabilization of actin organization under sustained deformation, manifested as a progressive loss of actin fibrosity and reduced capacity to maintain aligned, load-bearing structures (Fig. 4e). This behavior is consistent with experimental observations showing that vimentin is required to preserve actin integrity under mechanical load, and highlights the role of vimentin as a stabilizing scaffold rather than a primary driver of force generation. Further parametric analyses on vimentin and actin reorganization are shown in Supplementary Fig. S7.

**Figure 4.**
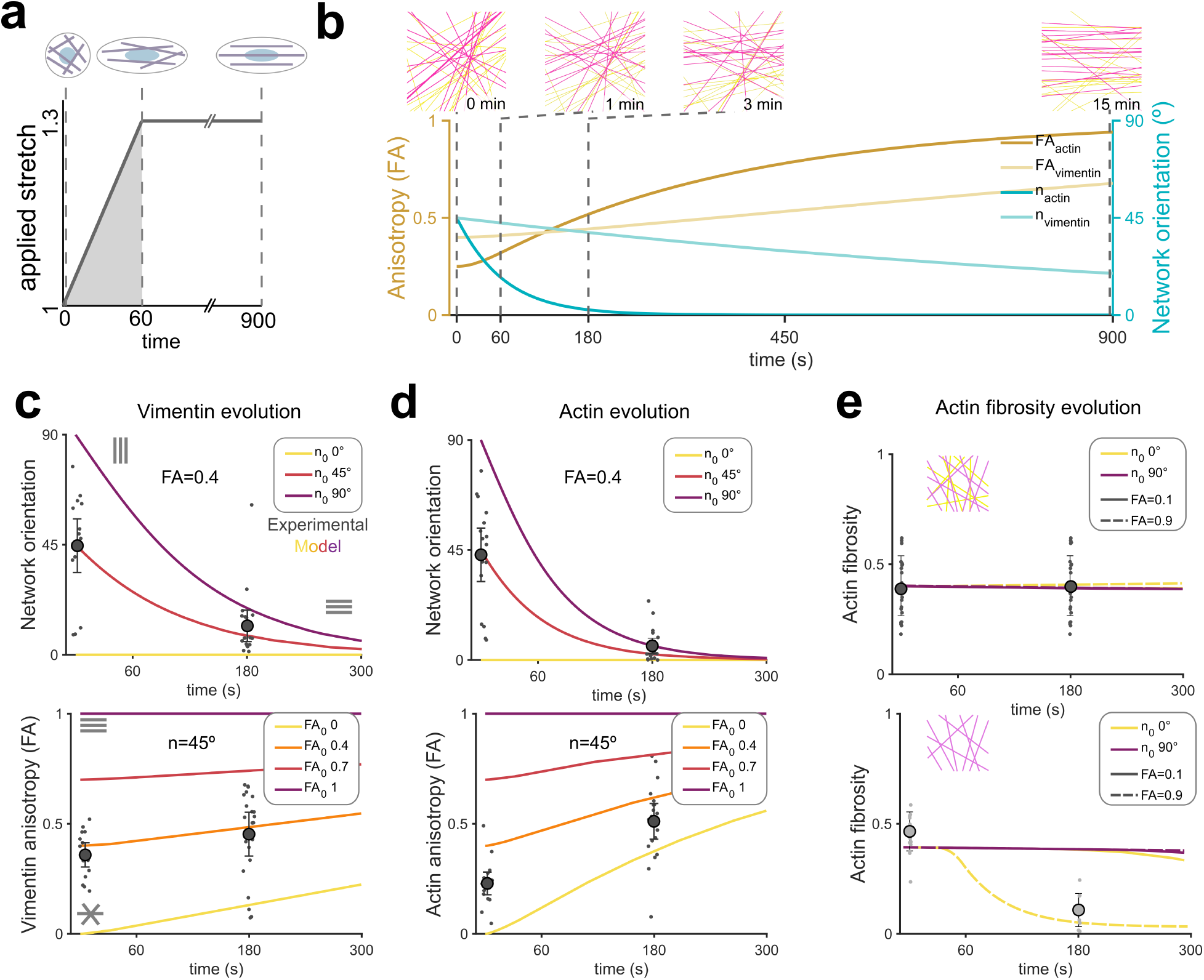
Time-dependent cytoskeletal reorganization emerges from model-guided dynamics under sustained deformation. **a** Applied stretch protocol used to probe cytoskeletal reorganization dynamics, consisting of a rapid stretch ramp followed by sustained deformation. **b** Model-predicted temporal evolution of actin and vimentin network anisotropy (FA) and principal orientation under sustained actuation, with representative fiber orientation maps shown at selected time points. **c,d** Experimental measurements (symbols) and model predictions (lines) of network orientation and anisotropy evolution for **c** vimentin and **d** actin networks under different initial alignment conditions, demonstrating progressive reorientation toward the principal stretch direction and network-specific remodeling kinetics. **e** Model-predicted evolution of actin fibrosity under sustained deformation for low and high initial anisotropy states in control and vimentin-deficient cells, illustrating the stabilizing role of vimentin on actin network maintenance during remodeling.

Together, these results demonstrate that the proposed constitutive model not only reproduces deformation-dependent cell mechanics, but also predicts the time-dependent structural reorganization of the cytoskeleton across networks and perturbations. The model reveals that cytoskeletal remodeling is inherently transient, with characteristic time scales governing the evolution and relaxation of anisotropy and alignment under sustained loading. This time-dependent behavior suggests that cells retain a partial memory of prior mechanical states encoded in their cytoskeletal organization. In the following section, we build on this insight to investigate how residual cytoskeletal order gives rise to short-term mechanical memory and to identify the characteristic relaxation processes that govern the transition between adaptive and memory-driven mechanical responses.

### 2.5. Residual cytoskeletal order enables short-term mechanical memory

In soft active materials, history-dependent mechanical responses are often difficult to interpret because inelastic (plastic) microstructural deformations and long characteristic-time viscous relaxation can produce similar macroscopic signatures [42, 43]. In ultra-soft magneto-responsive elastomers, for instance, responses that resemble yielding or permanent reconfiguration may instead arise from field-driven viscous mechanisms coupled to slow microstructural rearrangements, leading to apparent memory effects that persist over long times and can be partially reset after sufficient relaxation [44]. More broadly, these systems illustrate how mechanical history can be encoded transiently in an evolving internal architecture, with the observed response emerging from an interplay between instantaneous deformation, rate-dependent relaxation, and microstructural adaptation. Motivated by this conceptual framework, we posed whether glioblastoma cells exhibit a comparable microstructure-encoded short-term memory, and whether our constitutive model can predict these history effects without additional variations or calibration. To probe mechanical memory, we designed a two-step actuation protocol (Fig. 5a) consisting of a first stretching actuation (1 h), followed by a relaxation interval (3 min, 15 min, or 1 h), and a second actuation applying either traction or compression during 3 min or 1 h. The central hypothesis is that cytoskeletal remodeling is inherently transient, and that a fraction of the deformation-induced anisotropy may persist after unloading, thereby biasing subsequent reorganization and enabling memory-like behavior.

**Figure 5.**
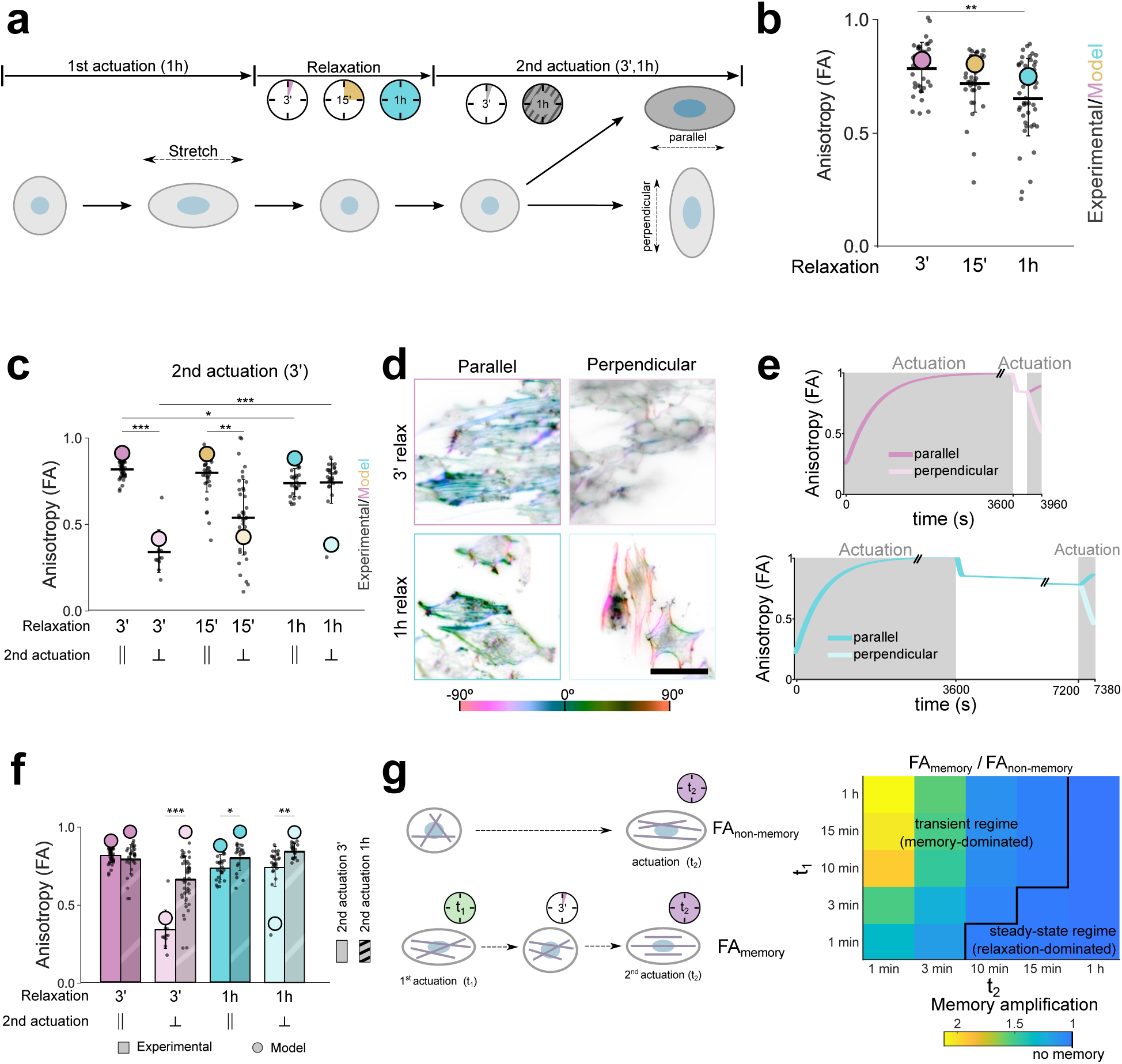
Residual cytoskeletal order encodes short-term mechanical memory in glioblastoma cells. **a** Mechanical memory actuation protocol consisting of a 1 h first uniaxial actuation, a relaxation period (3 min, 15 min, or 1 h), and a second actuation applied either parallel or perpendicular to the first direction for 3 min or 1 h. **b** Residual cytoskeletal anisotropy retained after unloading as a function of relaxation time. **c** Final anisotropy after the full memory protocol for the 3 min second actuation, comparing experimental measurements (dispersion plot) with model predictions (colored dots) for parallel and perpendicular reactuation. **d** Representative fiber orientation images at the end of the protocol for cells re-actuated for 3 min after either 3 min or 1 h relaxation, under parallel or perpendicular second actuation. **e** Model-predicted time evolution of anisotropy during the memory protocol for short (3 min) and long (1 h) relaxation. **f** Summary comparison of final anisotropy across relaxation times (3 min vs 1 h) and second actuation durations (3 min vs 1 h) for parallel and perpendicular re-actuation comparing experimental results and model predictions. **g** Heatmaps of the memory ratio, defined as the final anisotropic fraction after the full protocol normalized by the anisotropic fraction obtained under the corresponding single-actuation control (no-memory) for parallel and perpendicular reactuation. Scale bars: 20 *µ*m. n=20-30 cells analyzed per condition from N=3 independent assays. Experimental data are presented as mean *±* SEM. A two-tailed unpaired t-test with Welch’s correction was used for statistical analysis, with ns p > 0.05 (not shown), * p*≤*0.05, ** p*≤*0.01, *** p*≤*0.001.

We first quantified the mechanically induced anisotropy remaining after unloading. After the relaxation period, cells retained a substantial residual anisotropy, which decayed progressively as relaxation time increased (Fig. 5b). This result supports the idea that unloading does not instantly erase the structural state: instead, cytoskeletal organization relaxes on characteristic timescales, leaving a measurable “state variable” that can carry information forward into a subsequent loading event. We next tested whether this residual anisotropy translates into directional memory during reloading. When the second actuation was short (3 min), the final anisotropy strongly depended on whether reloading was applied parallel or perpendicular to the first deformation (Fig. 5c–e). A second actuation parallel to the first rapidly re-established a highly anisotropic state, consistent with a pre-aligned microstructure that can be reinforced with minimal additional remodeling. In contrast, a perpendicular reloading drove cells toward a markedly lower anisotropy at the protocol endpoint, consistent with competition between the residual alignment encoded by the first actuation and the new principal direction imposed by the second. Representative orientation maps (Fig. 5d and Supplementary Fig. S8) illustrate these distinct outcomes, with visibly reinforced alignment for parallel reloading and a more isotropic/disordered configuration after perpendicular reloading, particularly with short relaxation times.

Importantly, these experiments were used as a predictive test of the model: simulations were run using the previously identified parameters (i.e., no variations or recalibration at this stage), and only then compared against the experimental protocol outcomes. In this regime, the model reproduced the measured anisotropy trends across loading direction and short relaxation times (Fig. 5c,e and Supplementary Fig. S9), indicating that encoding cytoskeletal history via internal microstructural variables is sufficient to capture short-term memory. However, agreement deteriorated for the longest relaxation time (1 h), despite correct trends at shorter waits. We interpret this not as a contradiction of the cytoskeletal framework, but as evidence that the system transitions toward a broader cell-level adaptation regime when sufficient time is allowed between loading events. In particular, whereas short relaxation primarily probes the persistence (and partial decay) of cytoskeletal alignment, an hour-scale relaxation is expected to involve additional processes such as focal adhesion maturation/disassembly, changes in adhesion turnover and force transmission, and other slow reconfigurations of the cellular mechanical apparatus that effectively reshape boundary conditions for the second actuation. These elements are not explicitly represented in the present constitutive description, which was designed to encode history through cytoskeletal internal variables. The loss of predictivity at hour-scale relaxation thus likely reflects the onset of additional, adhesion- and coupling-mediated adaptation processes beyond cytoskeletal remodeling, which provides a clear direction for future multi-timescale extensions.

Finally, we asked whether memory persists when the second actuation is prolonged (1 h). In this case, the memory signature was largely diminished (Fig. 5f and Supplementary Figs. S8 and S10), outcomes approached those obtained under a single actuation of equivalent duration and direction, indicating that long reloading allows the system to “overwrite” prior history and converge toward a state dominated by the current loading conditions. Consistent with this interpretation, the ratio map comparing the anisotropic fraction achieved with and without prior mechanical history (Fig. 5g and Supplementary Fig. S11) showed the largest memory amplification for short timescales (short second actuation), while the ratio approached unity when the second actuation was long, for both parallel and perpendicular configurations i.e., the response became effectively indistinguishable from a simple one-step loading. Together, these results delineate a mechanically meaningful short-term memory window governed by transient cytoskeletal order: mechanical memory is strongest when the protocol interrogates the system before slow relaxation and broader cell-level remodeling processes erase or reconfigure the state that the first actuation encoded.

## 3. Discussion

Our results establish a direct structural link between cytoskeletal microstructural organization and cell-scale mechanical behavior. We show that deformation-induced cytoskeletal anisotropy is a key microstructural variable that controls both the magnitude and the sign of mechanical adaptation under load. Rather than behaving as a homogeneous material, the cytoskeleton reorganizes into architecture-specific states whose mechanical roles depend on the imposed deformation: acto-myosin fibers selectively drive stiffening under tensile loading, whereas branched cortical actin governs mechanical softening under compression, revealing a functional division of actin networks emerging from microstructural anisotropy. This cortical softening under in-plane compression appears to contrast with some reports in the literature. While numerous previous studies have investigated the response of the actin cortex under compression, this compression is typically applied via AFM indentation, which effectively loads the cortex primarily through out-of-plane bending and in-plane stretching of the membrane–cortex [45, 46]. In contrast, our approach induces in-plane compressive stresses that have previously been shown to trigger buckling of actin cortical networks [36]. We further demonstrate that vimentin intermediate filaments do not directly contribute to basal stiffness, but play a critical stabilizing role by preserving actin organization under mechanical load, complementing previous reports that vimentin stabilizes cells under compressive stress and promotes compression stiffening [47]. In contrast, microtubule depolymerization did not measurably affect cell mechanics under basal or compressive conditions in our experiments. However, given their intimate crosstalk with actin and intermediate filaments, more subtle or indirect contributions to mechanically induced short-term memory cannot be excluded [48, 27]. Notably, microtubules can resist compressive stresses and mechanical breakage, and posttranslational modifications such as acetylation enhance their stability under load, likely operating on longer timescales [49, 50]. Importantly, deformation induces cytoskeletal reorganization and anisotropy that persist transiently after unloading, leaving a residual structural state whose relaxation occurs over characteristic time scales and gives rise to a short-term mechanical memory linking deformation history to subsequent mechanical and structural responses. These experimental findings are unified and rationalized through a multi-network constitutive framework that explicitly couples deformation, cytoskeletal anisotropy and remodeling dynamics to cell-scale mechanics. By quantitatively reproducing both mechanical responses and cytoskeletal dynamics across deformation modes and perturbations, the model was instrumental in testing mechanistic hypotheses and in establishing transient cytoskeletal anisotropy as a structural mechanism encoding short-term mechanical memory in living cells.

The present study focuses on short time scales, capturing transient cytoskeletal remodeling and short-term mechanical memory. Importantly, deformation-induced cytoskeletal alignment does not translate into stable cell polarization, as supplementary analyses reveal minimal reorganization of the nucleus–centrosome axis despite clear anisotropic deformation of the cell body (Supplementary Fig. S12). Thus, transient cytoskeletal memory can bias filament organization and global morphology but is insufficient, on these time scales, to establish a fully polarized migratory phenotype. In contrast, nuclear morphology appears more sensitive to mechanical deformation. Sustained actuation induces alignment and deformation of the nucleus along the principal stretch direction, with enhanced nuclear strain observed in vimentin-deficient cells (Supplementary Fig. S13). These findings support the idea that cytoskeletal anisotropy can be mechanically transmitted to the nucleus, potentially modulating nuclear mechanics in a manner dependent on intermediate filament integrity [33, 51, 52]. In light of recent work linking increased nuclear deformation in vimentin knockout cells to altered migratory behavior and transcriptional responses [52], these observations suggest that cytoskeletal reorganization may propagate toward longer-term regulatory processes through nucleus–cytoskeleton coupling. Moreover, the pronounced increase in actin stress fiber alignment upon stretching suggests that focal adhesion size and spatial distribution are likely remodeled on similar timescales. Slower processes, including focal adhesion maturation [53], nucleus-cytoskeleton coupling [54], transcriptional regulation [55] and long-term phenotypic adaptation [56, 57, 58], are not explicitly addressed here and may contribute to longer-lived forms of mechanical memory. Experimentally, extending the current approach to include migration and motility assays, as well as three-dimensional matrices and confined environments, would help determine how the transient cytoskeletal memory mechanisms identified here impact persistent migration, invasion efficiency and long-time mechanical adaptation in physiologically relevant settings. Together, such extensions would enable a more complete assessment of how deformation history, cytoskeletal remodeling and force transmission jointly regulate cell behavior in vivo.

From a modeling perspective, the constitutive framework adopts an isostrain assumption at the macroscopic scale, whereby all cytoskeletal networks experience the same deformation gradient. At the same time, each network is endowed with its own internal variables and branch-specific kinematics, including distinct internal deformation measures and remodeling laws. Thus, although the formulation operates within a continuum mixture description, the individual cytoskeletal contributions are not mechanically independent, as they are physically interconnected through cytolinker proteins such as plectin that bridge intermediate filaments, microtubules and F-actin [59]. These interactions are incorporated in the model through explicit dependencies in the evolution equations governing anisotropy, phase fractions and internal reference configurations. In this sense, the framework captures inter-network coupling at the level of remodeling kinetics and load-bearing capacity, while remaining computationally tractable at the cell scale. While this description is appropriate at the cell scale and for the time scales considered, the underlying formulation is fully consistent with spatially resolved implementations. In particular, the present framework can be naturally extended to finite element settings that account for spatially heterogeneous distributions of cytoskeletal networks, anisotropy and internal state variables within the cell [29, 31]. Such extensions would enable the explicit representation of local decoupling between networks, as well as the incorporation of focal adhesions and nucleus-cytoskeleton interactions, providing a route to address spatial heterogeneity, long-time relaxation and mechanically regulated migration in confined environments. Another interesting approach to explore is coupling the present framework with molecular clutch models [60, 61]. This would provide a natural route to account for the effective transmission of substrate-derived mechanical cues to the cytoskeleton and to explore how cytoskeletal memory interacts with adhesion dynamics.

Placed in a physiological context, the mechanisms identified here are particularly relevant given the highly dynamic mechanical environments experienced by glioblastoma cells in vivo. Tumor growth, solid stress, edema, inflammation and tissue heterogeneity generate evolving sequences of compression, relaxation and tensile loading, rather than static mechanical conditions [62, 63, 64]. Our results suggest that such deformation histories can transiently bias cytoskeletal organization through residual anisotropy, thereby modulating subsequent mechanical responses and endowing cells with a short-term, history-dependent mechanical state [65, 58]. This transient mechanical memory provides a structural mechanism by which glioblastoma cells may enhance migratory efficiency and persistence under fluctuating mechanical constraints. When the characteristic time between mechanical events in the tumour microenvironment is comparable to cytoskeletal relaxation times, residual anisotropy can bias force generation, polarization and reorganization during invasion. In this regime, vimentin-mediated stabilization of actin architectures is expected to promote mechanical robustness and adaptability, not by increasing stiffness per se, but by preserving the integrity of load-bearing cytoskeletal structures across successive deformation events. Beyond migration, deformation-history dependent cytoskeletal states may also influence proliferation and mechanotransduction pathways implicated in tumor progression [57, 66]. Mechanically regulated signaling nodes associated with actomyosin contractility, cortical organization [67] and mechanosensitive channels [68] are likely to be transiently modulated by prior deformation, suggesting that short-term mechanical memory can shape cell behavior on time scales relevant to tumor dynamics [69]. In this sense, cytoskeletal anisotropy emerges not only as a mechanical descriptor, but as a dynamic internal variable linking mechanical history to downstream cellular responses. While these findings provide a mechanistic framework for deformation-history dependent cytoskeletal remodeling in glioblastoma cells, several limitations should be noted. Experiments were performed primarily in U251-MG cells and a vimentin knockout derivative cultured on 2D substrates; generalization to other GBM lines or in 3D and confined microenvironments remains to be established. Moreover, measurements were obtained at a defined substrate stiffness, and given the well-known dependence of cytoskeletal organization and force transmission on matrix rigidity and adhesion, the magnitude of the observed responses may vary across different mechanical environments.

From a translational and conceptual perspective, the combined experimental-theoretical framework introduced here provides a powerful platform to interrogate how cytoskeletal architecture, deformation mode and mechanical history jointly regulate cell behavior. The constitutive model enables in silico testing of mechanisms and helps interpret complex, history-dependent responses that are otherwise difficult to disentangle experimentally. More broadly, by positioning glioblastoma cells as active materials with microstructure-dependent memory, this work establishes a general framework to explore mechanically informed therapeutic strategies, including the identification of strategies to disrupt mechanical adaptability by targeting the structural and dynamical rules governing cytoskeletal anisotropy.

## 4. Methods

### Cell culture

U251-MG cells were cultured growth medium containing Minimum Essential Medium (MEM) with Glutamax, supplemented with 10% FBS, 1% penicillin-streptomycin, and 1% Non-Essential AminoAcids (10 mM of each amino acid). Cells were cultured under standard conditions of 37°C and 5% CO_2_. Cell cultures were passaged every 3-4 days at 1:10 cell concentration.

U251 vimentin knockout cells were obtained as described in [33]. Briefly, U251 GBM cells were transfected with guide RNAs-containing plasmids targeting vimentin exon 2. Transfection was done by electroporation in Mirus (Lonza) transfection solution with a Nucleofector device (Lonza). Vimentin knockout was determined using western blot analysis, and complete disruption of vimentin networks was confirmed by immunofluorescence.

### Generation of a U251-MG Vimentin–mCherry Knock-In Cell Line (Vim-mCherry KI)

Endogenous tagging of vimentin with mCherry was performed using CRISPR–Cas9–mediated homology-directed repair. A single guide RNA targeting exon 10 (sgRNA: UCAGGAGCGCAAGAUAGAUU) was cloned into the pSpCas9(BB)-2A-GFP (PX458) vector (#48138, Addgene) using BbsI restriction sites, according to Ran et al. method [70]. A donor repair template was designed to introduce mCherry in frame with the endogenous coding sequence. This template contained a 600 bp 5’ homology arm, a flexible Gly-Ser linker, the mCherry coding sequence, and a 600 bp 3’ homology arm. Homology arms and linker–mCherry were respectively amplified from genomic and plasmid DNA (Supplementary Table S2). Fragments were assembled using Gibson assembly, cloned into the PCR™Blunt II-TOPO™ vector and amplified. The donor template was linearized by EcoRI digestion prior to delivery. U251-MG cells were co-transfected with the PX458-sgRNA plasmid and linearized donor template using Nucleofector 2b Basic Kit for Glial Cells (Lonza) and the Nucleofector™ II (Amaxa), according to manufacturer’s instructions (Prog: T20[71]). Cas9-GFP-positive cells were enriched by FACS 48h after nucleofection. A subsequent round of FACS was performed to isolate mCherry-positive cells. Correct insertion at the endogenous locus was confirmed by genomic sequencing, western blotting, and immunofluorescence analysis (Supplementary Figure S14).

### Western blot of Vim-mCherry KI

Cells were lysed in LDS sample buffer (Merck) supplemented with 100 mM DTT, heated at 95°C for 5 min, and sonicated. Proteins were separated on 4–12% Bis–Tris gels (Merck) and transferred onto PVDF membranes (Merck). Membranes were blocked in 5% milk in TBSTween or in EveryBlot Blocking Buffer (Bio-Rad). They were incubated either with primary antibodies for 1h at room temperature, followed by HRP-conjugated secondary antibodies, or directly with fluorescently labelled primary antibodies. Antibodies were diluted in 5% milk in TBS-Tween or EveryBlot Blocking Buffer (Bio-Rad). Signal detection was performed using enhanced chemiluminescence (Bio-Rad Clarity) and imaged using a ChemiDoc MP system (Bio-Rad). Quantification was performed using Fiji, with normalization to GAPDH. Western blots were quantified using Fiji/ImageJ. The fraction of tagged protein was calculated as the ratio of tagged Vimentin to total Vimentin (tagged and untagged). Statistical analyses were performed in Prism (Wilcoxon test for relative expression; Mann–Whitney test for fraction of tagged protein).Antibodies used are detailed in Supplementary Table S3.

### Drug treatments

To study the effect of cytoskeletal disruption on cell mechanics in relaxed and actuated conditions, cultured cells were subjected to different drug treatments. Cells were incubated with 1 *µ*M cytochalasin D, 10 *µ*M Nocodazole + 1 *µ*M Y-27632, 10 *µ*M Y-27632, or 50 *µ*M CK666 for 30 minutes at 37°C and 5% CO_2_ prior to analysis.

### Magneto-mechanical actuation system

Mechanical actuation experiments were performed using the magneto-mechanical stimulation platform NeoMag (60Nd S.L., Madrid), which is directly derived from the magnetically actuated system originally developed and validated in our previous work [34]. The device enables controlled, contactless application of mechanical deformation to adherent cells through externally generated magnetic fields, while remaining fully compatible with standard cell culture and microscopy workflows. Mechanical actuation is achieved by generating a controlled magnetic field that induces deformation of magneto-active substrates coupled to the cellular microenvironment. The applied magnetic field is generated by permanent magnets whose relative position is precisely controlled, allowing for reproducible deformation modes. To enable high-resolution imaging and seamless integration with inverted fluorescence microscopy, the magneto-active substrates were adapted for this study. Specifically, cells were cultured on custom-fabricated magneto-active elastomeric substrates mounted in P35 plastic culture dishes with enlarged inner apertures, allowing optical access with high–numerical aperture objectives. These modifications preserve the mechanical actuation capabilities of the system while enabling simultaneous mechanical stimulation and fluorescence imaging. The effective transmission of substrate deformation to adherent cells was verified by direct observation of immediate cell elongation upon actuation.

### Dynamic indentation setup and measurement protocol

Indentation measurements were performed 48-72 h after seeding the U251 cells. All measurements were done with cells in complete medium supplemented with 10mM HEPES and without phenol-red to avoid changes in medium color that may affect the measurements. Indentation probes of 0.015-0.033 N/m stiffness and 3-3.5 *µ*m spherical tip radius were used with a Chiaro nanoindentation system (Optics11 Life, Netherlands). Displacement-controlled monotonous indentation was performed with a final indentation of 4 *µ*m, corresponding to approximately ∼2/3 of the bead diameter, with an indentation rate of 1 *µ*m/s. The raw data was analyzed using Piuma Viewer software. Young’s modulus was obtained from the slope at the start of the indentation following the Hertzian model, assuming a Poisson ratio of 0.5.

### Immunofluorescence staining

Cells were fixed 10 min at room temperature in cytoskeleton buffer (MES 10mM, KCl 138mM, MgCl 3mM, EGTA 2mM) supplemented with 10% sucrose, 0.1% Triton-X100, 0.05% glutaraldehyde and 4% paraformaldehyde (PFA), as described in [72]. Fixation was followed by incubation in 1 mg/ml NaBH_4_ in PBS solution for 10 min to reduce the aldehyde functions. Samples were then incubated with 0.1% Triton-X100 for 10 minutes and blocked for 1 h at room temperature. Samples were washed 3 times with PBS between each step. Primary antibody incubation was performed in 2% BSA in PBS for 1 h at room temperature. Secondary antibody, Hoetsch and labelled phalloidin incubation was performed in 2% BSA in PBS for 1 h at room temperature. Samples were washed three times with 2% BSA in PBS for 5 min each after each antibody incubation. Samples were mounted in Prolong Diamond (Invitrogen, P36961). Images were acquired with a Leica DM6B epifluorescent microscope equipped with a x63 1.4 NA oil objective and recorded on a CCD Leica DFC3000G camera with Leica Application Suite X Software or with a Nikon Ti Eclipse 2 with Yokogawa Spinning unit CSU W1, equipped with a Plan Apo x60 1.4 NA oil Nikon objective and recorded on a Prime BSI Express sCMOS camera (Teledyne Photometrics, Tucson, AZ, USA) with Micromanager Software (purchased through Gataca systems).

### Image processing and data analysis

Most image processing and analysis were performed using the Fiji software [73]. Visualization of cytoskeletal orientation was performed using the OrientationJ plugin [74]. For cytoskeletal analysis, a mask of the cell contour and fluorescence images of cytoskeletal markers were used for single-cell quantification of cytoskeleton structures using the CSKMorphometrics algorithm in MATLAB [75]. Additional processing, graph design, and statistical analysis were performed using custom Python codes. Statistical analyses are listed in the figure legends with p-values (ns p ≥ 0.05, * p≤0.05, ** p≤0.01, *** p≤0.001 **** p≤0.0001). Statistical comparisons are unpaired t-tests or two-way ANOVA with multiple conditions.

### Constitutive model

The mechanical behavior of glioblastoma cells was described using a continuum constitutive model explicitly grounded in cytoskeletal structure and remodeling. Motivated by the experimental evidence presented above, the cell was modeled as an incompressible continuum whose macroscopic mechanical response emerges from the combined contributions of multiple cytoskeletal networks and an isotropic background. This framework enables a direct link between cytoskeletal organization, deformation history, and cell-scale mechanical response. The total Helmholtz free-energy density Ψ was assumed to decompose additively into contributions associated with an isotropic background, the vimentin intermediate filament network, and the actin cytoskeleton,

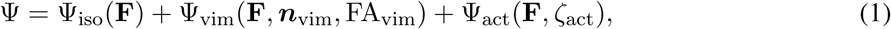

where **F** denotes the deformation gradient and *ζ*_act_ are state variables associated with the actin network. The isotropic contribution Ψ_iso_ represents the effective mechanical response of the cytosolic matrix and other cellular components that are not explicitly modeled as filamentous networks and depends solely on the deformation gradient. The vimentin contribution Ψ_vim_ accounts for the anisotropic mechanical response of the vimentin intermediate filament network and depends on the deformation gradient, the preferred vimentin orientation ***n***_vim_, and the fractional anisotropy FA_vim_ describing the degree of network alignment.

The actin contribution is written as a weighted sum of three actin phases,

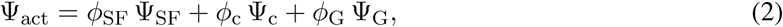

where *ϕ*_SF_, *ϕ*_c_, and *ϕ*_G_ denote the fractions of actin incorporated into stress fibers, actin cortex, and monomeric (G-)actin, respectively, and satisfy *ϕ*_SF_ + *ϕ*_c_ + *ϕ*_G_ = 1. The stress-fiber contribution is defined as

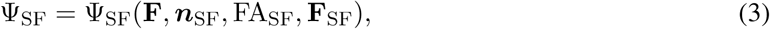

where ***n***_SF_ is the preferred stress-fiber orientation, FA_SF_ is the stress-fiber fractional anisotropy, and **F**_SF_ is an internal variable representing the evolving elastic reference configuration of the stress-fiber network. The actin cortex contribution is defined as

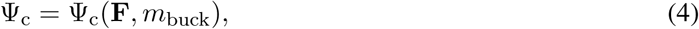

where *m*_buck_ is a scalar buckling reduction factor that modulates the effective mechanical contribution of the cortex under compressive in-plane deformation. The G-actin contribution Ψ_G_ does not introduce an effective mechanical response.

The first Piola–Kirchhoff stress tensor is obtained from the free-energy density through

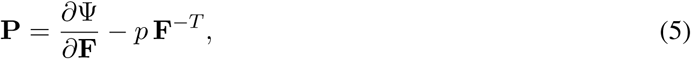

where *p* is a Lagrange multiplier enforcing incompressibility.

All constituents are assumed to experience the same macroscopic deformation gradient at the continuum scale, consistent with an isostrain assumption. Nevertheless, the networks are not mechanically independent, as inter-network coupling is incorporated through their respective internal variables and evolution equations. Time-dependent cytoskeletal remodeling is captured through evolution equations for selected internal variables. In the present model, the preferred orientations ***n***_vim_ and ***n***_SF_, the fractional anisotropies FA_vim_ and FA_SF_, the stress-fiber internal reference configuration **F**_SF_, the actin phase fractions *ϕ*_SF_, *ϕ*_c_, and *ϕ*_G_, and the cortical buckling variable *m*_buck_ evolve in time according to deformation-dependent flow rules. These evolution laws account for deformation-driven reorientation, anisotropy development, actin turnover between phases, stress-fiber remodeling, and cortical buckling and recovery. The specific functional forms of the free-energy contributions, the evolution equations, and all model parameters are provided in the Supplementary Information.

## Acknowledgements

The authors acknowledge support from the European Research Council (ERC) under the European Union’s Horizon 2020 Research and Innovation Programme (Grant agreement No. 947723, project: 4D-BIOMAP, and Grant agreement No. 101247449, project: MAGMATED) and from the European Innovation Council (EIC) under the Horizon Europe Programme (Grant agreement No. 101284471, project: NEOMAG). The authors acknowledge support from from MICIU/AEI/10.13039/501100011033 under Grant number PID2023-149255NB-I00 and PID2023-152631OB-I00 and from FEDER, UE. CGC acknowledges support from the Ministerio de Universidades, Spain (FPU20/01459). This work was also supported by La Ligue contre le cancer (EL2023 - DN/IP/IQ – 17691), Worldwide Cancer Research (WCR 23-0156), INSERM PCSI (N° 22CP073-00) and Institut Pasteur (PTR-548-22).

## Contributions

CGC and DGG conceived and conceptualized the study with inputs from MG, SEM and AMB. Imaging experiments were performed by CGC, with support from MG and supervision from SEM. Mechanical experiments were conducted by CGC under the supervision of DGG. LP developed and characterized the m-cherry U251 cell line. DGG and CGC developed the theoretical model and its numerical implementation. CGC and DGG wrote the original manuscript. All authors discussed the results and reviewed the manuscript.

## Ethics Declarations

Daniel Garcia-Gonzalez is co-founder of the spin-off 60Nd S.L. that commercializes magneto-mechanical actuation technology for mechanobiology. Clara Gomez-Cruz is an employee of 60Nd S.L.

## References

[1] L. Przybyla, J. M. Muncie, V. M. Weaver, Mechanical control of epithelial-to-mesenchymal transitions in development and cancer, Annual review of cell and developmental biology 32 (1) (2016) 527–554.

[2] B. Kuehlmann, C. A. Bonham, I. Zucal, L. Prantl, G. C. Gurtner, Mechanotransduction in wound healing and fibrosis, Journal of clinical medicine 9 (5) (2020) 1423.

[3] Y. Lu, C. Chen, H. Li, P. Zhao, Y. Zhao, B. Li, W. Zhou, G. Fan, D. Guan, Y. Zheng, Visible light-responsive hydrogels for cellular dynamics and spatiotemporal viscoelastic regulation, Nature communications 16 (1) (2025) 1365.

[4] X. Di, X. Gao, L. Peng, J. Ai, X. Jin, S. Qi, H. Li, K. Wang, D. Luo, Cellular mechanotransduction in health and diseases: from molecular mechanism to therapeutic targets, Signal transduction and targeted therapy 8 (1) (2023) 282.

[5] D. Garcia-Gonzalez, Magneto-mechanics in mechanobiology: enabling remote force transmission to cells and extracellular matrix, Biophysical Reviews (Nov 2025).

[6] D. J. Tschumperlin, D. Lagares, Mechano-therapeutics: targeting mechanical signaling in fibrosis and tumor stroma, Pharmacology & therapeutics 212 (2020) 107575.

[7] J. W. N. Lee, A. W. Holle, Engineering approaches for understanding mechanical memory in cancer metastasis, APL bioengineering 8 (2) (2024).

[8] T. A. Ulrich, E. M. de Juan Pardo, S. Kumar, The mechanical rigidity of the extracellular matrix regulates the structure, motility, and proliferation of glioma cells, Cancer research 69 (10) (2009) 4167–4174.

[9] A. Isomursu, K.-Y. Park, J. Hou, B. Cheng, G. Shamsan, B. Fuller, J. Kasim, M. M. Mahmoodi, T. J. Lu, G. M. Genin, F. Xu, M. Lin, M. Distefano, J. Ivaska, D. J. Odde, Negative durotaxis: cell movement toward softer environments, preprint, Biophysics (Oct. 2020).

[10] K.-J. Streitberger, L. Lilaj, F. Schrank, J. Braun, K.-T. Hoffmann, M. Reiss-Zimmermann, J. A. Käs, I. Sack, How tissue fluidity influences brain tumor progression, Proceedings of the National Academy of Sciences 117 (1) (2020) 128–134.

[11] M. Walker, P. Rizzuto, M. Godin, A. E. Pelling, Structural and mechanical remodeling of the cytoskeleton maintains tensional homeostasis in 3d microtissues under acute dynamic stretch, Scientific reports 10 (1) (2020) 7696.

[12] B. Cheng, M. Li, M. Lin, H. Guo, F. Xu, Mechanobiology across timescales, Nature Reviews Physics 7 (11) (2025) 621–644.

[13] S. K. Kureel, R. Maroto, K. Davis, M. Sheetz, Cellular mechanical memory: a potential tool for mesenchymal stem cell-based therapy, Stem Cell Research & Therapy 16 (1) (2025) 159.

[14] E. Cambria, M. F. Coughlin, M. A. Floryan, G. S. Offeddu, S. E. Shelton, R. D. Kamm, Linking cell mechanical memory and cancer metastasis, Nature Reviews Cancer 24 (3) (2024) 216–228.

[15] A. E. Beedle, A. Jaganathan, A. Albajar-Sigalés, F. M. Yavitt, K. Bera, I. Andreu, I. Granero-Moya, D. Zalvidea, Z. Kechagia, G. Wiche, et al., Fibrillar adhesion dynamics govern the timescales of nuclear mechano-response via the vimentin cytoskeleton, BioRxiv (2023).

[16] C. Anton, F. Lautenschläger, R. J. Hawkins, Modeling cytoskeletal and cell dynamics, Current Opinion in Cell Biology 97 (2025) 102584.

[17] S. Massou, F. Nunes Vicente, F. Wetzel, A. Mehidi, D. Strehle, C. Leduc, R. Voituriez, O. Rossier, P. Nassoy, G. Giannone, Cell stretching is amplified by active actin remodelling to deform and recruit proteins in mechanosensitive structures, Nature cell biology 22 (8) (2020) 1011–1023.

[18] C. A. Dessalles, N. Cuny, A. Boutillon, P. F. Salipante, A. Babataheri, A. I. Barakat, G. Salbreux, Interplay of actin nematodynamics and anisotropic tension controls endothelial mechanics, Nature Physics 21 (6) (2025) 999–1008.

[19] S. Seetharaman, S. Etienne-Manneville, Cytoskeletal crosstalk in cell migration, Trends in cell biology 30 (9) (2020) 720–735.

[20] M. C. Keeling, L. R. Flores, A. H. Dodhy, E. R. Murray, N. Gavara, Actomyosin and vimentin cytoskeletal networks regulate nuclear shape, mechanics and chromatin organization, Scientific reports 7 (1) (2017) 5219.

[21] S. Kasas, X. Wang, H. Hirling, R. Marsault, B. Huni, A. Yersin, R. Regazzi, G. Grenningloh, B. Riederer, L. Forro, et al., Superficial and deep changes of cellular mechanical properties following cytoskeleton disassembly, Cell motility and the cytoskeleton 62 (2) (2005) 124–132.

[22] J. P. Conboy, M. G. Lettinga, P. E. Boukany, F. C. MacKintosh, G. H. Koenderink, Actin and vimentin jointly control cell viscoelasticity and compression stiffening, Molecular Biology of the Cell (2026) mbc–E24.

[23] S. Amiri, C. Muresan, X. Shang, C. Huet-Calderwood, M. A. Schwartz, D. A. Calderwood, M. Murrell, Intracellular tension sensor reveals mechanical anisotropy of the actin cytoskeleton, Nature communications 14 (1) (2023) 8011.

[24] N. Gavara, P. Roca-Cusachs, R. Sunyer, R. Farré, D. Navajas, Mapping cell-matrix stresses during stretch reveals inelastic reorganization of the cytoskeleton, Biophysical journal 95 (1) (2008) 464–471.

[25] S. Nageswaran, J. Haipeter, J. F. Bodenschatz, R. Meyer, S. Koster, C. Steinem, Membrane-bound vimentin filaments reorganize and elongate under strain, Biomacromolecules 24 (6) (2023) 2512–2521.

[26] F. Alisafaei, K. Mandal, R. Saldanha, M. Swoger, H. Yang, X. Shi, M. Guo, H. Hehnly, C. A. Castañeda, P. A. Janmey, et al., Vimentin is a key regulator of cell mechanosensing through opposite actions on actomyosin and microtubule networks, Communications Biology 7 (1) (2024) 658.

[27] L. Pradeau-Phélut, S. Etienne-Manneville, Cytoskeletal crosstalk: A focus on intermediate filaments, Current Opinion in Cell Biology 87 (2024) 102325.

[28] H. Wu, Y. Shen, S. Sivagurunathan, M. S. Weber, S. A. Adam, J. H. Shin, J. J. Fredberg, O. Medalia, R. Goldman, D. A. Weitz, Vimentin intermediate filaments and filamentous actin form unexpected interpenetrating networks that redefine the cell cortex, Proceedings of the National Academy of Sciences 119 (10) (2022) e2115217119.

[29] H. Yang, T. Henzel, E. M. Stewart, M. Guo, An interpenetrating-network theory of the cytoskeletal networks in living cells, Journal of the Mechanics and Physics of Solids 189 (2024) 105688.

[30] J.-T. Hang, Y. Kang, G.-K. Xu, H. Gao, A hierarchical cellular structural model to unravel the universal power-law rheological behavior of living cells, Nature communications 12 (1) (2021) 6067.

[31] H. Borja da Rocha, J. Bleyer, H. Turlier, A viscous active shell theory of the cell cortex, Journal of the Mechanics and Physics of Solids 164 (2022) 104876.

[32] D. R. Katti, K. S. Katti, Cancer cell mechanics with altered cytoskeletal behavior and substrate effects: A 3d finite element modeling study, Journal of the mechanical behavior of biomedical materials 76 (2017) 125–134.

[33] E. J. van Bodegraven, E. Infante, F. Peglion, D. Pereira, Y. Kesenci, V. Roca, I. Perfettini, E. Terriac, J. Geay, L. Soto, et al., Intermediate filaments promote glioblastoma cell invasion by controlling nuclear deformations and mechanosensitive expression of mmp14, Cell Reports 44 (11) (2025).

[34] C. Gomez-Cruz, M. Fernandez-de la Torre, D. Lachowski, M. Prados-de Haro, A. E. del Río Hernández, G. Perea, A. Muñoz-Barrutia, D. Garcia-Gonzalez, Mechanical and functional responses in astrocytes under alternating deformation modes using magneto-active substrates, Advanced Materials 36 (26) (2024) 2312497.

[35] H. Ni, Q. Ni, G. A. Papoian, A. Trache, Y. Jiang, Myosin and alpha-actinin regulation of stress fiber contractility under tensile stress, Scientific Reports 13 (1) (2023) 8662.

[36] Y. Ideses, V. Erukhimovitch, R. Brand, D. Jourdain, J. S. Hernandez, U. Gabinet, S. A. Safran, K. Kruse, A. Bernheim-Groswasser, Spontaneous buckling of contractile poroelastic actomyosin sheets, Nature communications 9 (1) (2018) 2461.

[37] M. P. Murrell, M. L. Gardel, F-actin buckling coordinates contractility and severing in a biomimetic actomyosin cortex, Proceedings of the National Academy of Sciences 109 (51) (2012) 20820–20825.

[38] R. Kusters, C. Simon, R. Lopes Dos Santos, V. Caorsi, S. Wu, J.-F. Joanny, P. Sens, C. Sykes, Actin shells control buckling and wrinkling of biomembranes, Soft Matter 15 (2019) 9647–9653.

[39] M. P. Serres, M. Samwer, B. A. T. Quang, G. Lavoie, U. Perera, D. Görlich, G. Charras, M. Petronczki, P. P. Roux, E. K. Paluch, F-actin interactome reveals vimentin as a key regulator of actin organization and cell mechanics in mitosis, Developmental cell 52 (2) (2020) 210–222.

[40] K. Pogoda, L. Chin, P. C. Georges, F. J. Byfield, R. Bucki, R. Kim, M. Weaver, R. G. Wells, C. Marcinkiewicz, P. A. Janmey, Compression stiffening of brain and its effect on mechanosensing by glioma cells, New Journal of Physics 16 (7) (2014) 075002, publisher: IOP Publishing.

[41] P. Monzo, M. Crestani, Y. K. Chong, A. Ghisleni, K. Hennig, Q. Li, N. Kakogiannos, M. Giannotta, C. Richichi, T. Dini, E. Dejana, P. Maiuri, M. Balland, M. P. Sheetz, G. Pelicci, B. T. Ang, C. Tang, N. C. Gauthier, Adaptive mechanoproperties mediated by the formin fmn1 characterize glioblastoma fitness for invasion, Developmental Cell 56 (20) (2021) 2841–2855.e8.

[42] M. Moreno, J. Gonzalez-Rico, M. Lopez-Donaire, A. Arias, D. Garcia-Gonzalez, New experimental insights into magneto-mechanical rate dependences of magnetorheological elastomers, Composites Part B: Engineering 224 (2021) 109148.

[43] C. Perez-Garcia, R. Ortigosa, J. Martínez-Frutos, D. Garcia-Gonzalez, Topology and material optimization in ultra-soft magneto-active structures: Making advantage of residual anisotropies, Advanced Materials n/a (n/a) e18489.

[44] E. Gonzalez-Saiz, M. L. Lopez-Donaire, L. Gutiérrez, K. Danas, D. Garcia-Gonzalez, Magnetic-driven viscous mechanisms in ultra-soft magnetorheological elastomers offer history-dependent actuation with reprogrammability options, Advanced Science 12 (35) (2025) e06790.

[45] J. Hu, S. Chen, W. Hu, S. Lü, M. Long, Mechanical point loading induces cortex stiffening and actin reorganization, Biophysical Journal 117 (8) (2019) 1405–1418.

[46] P. Chugh, A. G. Clark, M. B. Smith, D. A. D. Cassani, K. Dierkes, A. Ragab, P. P. Roux, G. Charras, G. Salbreux, E. K. Paluch, Actin cortex architecture regulates cell surface tension, Nature Cell Biology 19 (6) (2017) 689–697.

[47] K. Pogoda, F. Byfield, P. Deptuła, M. Ciesluk, Ł. Suprewicz, K. Skłodowski, J. L. Shivers, A. Van Oosten, K. Cruz, E. Tarasovetc, et al., Unique role of vimentin networks in compression stiffening of cells and protection of nuclei from compressive stress, Nano Letters 22 (12) (2022) 4725–4732.

[48] M. Dogterom, G. H. Koenderink, Actin–microtubule crosstalk in cell biology, Nature reviews Molecular cell biology 20 (1) (2019) 38–54.

[49] Z. Xu, L. Schaedel, D. Portran, A. Aguilar, J. Gaillard, M. P. Marinkovich, M. Théry, M. V. Nachury, Microtubules acquire resistance from mechanical breakage through intralumenal acetylation, Science 356 (6335) (2017) 328–332.

[50] Y. Li, O. Kučera, D. Cuvelier, D. M. Rutkowski, M. Deygas, D. Rai, T. Pavlovič, F. N. Vicente, M. Piel, G. Giannone, et al., Compressive forces stabilize microtubules in living cells, Nature materials 22 (7) (2023) 913–924.

[51] A. E. Patteson, A. Vahabikashi, K. Pogoda, S. A. Adam, K. Mandal, M. Kittisopikul, S. Sivagurunathan, A. Goldman, R. D. Goldman, P. A. Janmey, Vimentin protects cells against nuclear rupture and dna damage during migration, Journal of Cell Biology 218 (12) (2019) 4079–4092.

[52] I. Elvira, T. Emmanuel, G. Matthieu, S. Hugo, P. David, R. Vanessa, V. Hugo, K. Sara, R. Reinier, A. Atef, et al., Intratumoral heterogeneity of vimentin modulates nuclear mechanotransduction, dna damage response and cancer cell survival, bioRxiv (2025) 2025–06.

[53] A. Elosegui-Artola, R. Oria, Y. Chen, A. Kosmalska, C. Pérez-González, N. Castro, C. Zhu, X. Trepat, P. Roca-Cusachs, Mechanical regulation of a molecular clutch defines force transmission and transduction in response to matrix rigidity, Nature Cell Biology 18 (5) (2016) 540–548.

[54] C. Guilluy, L. D. Osborne, L. Van Landeghem, L. Sharek, R. Superfine, R. Garcia-Mata, K. Burridge, Isolated nuclei adapt to force and reveal a mechanotransduction pathway in the nucleus, Nature Cell Biology 16 (4) (2014) 376–381.

[55] A. Elosegui-Artola, I. Andreu, A. E. Beedle, A. Lezamiz, M. Uroz, A. J. Kosmalska, R. Oria, J. Z. Kechagia, P. Rico-Lastres, A.-L. Le Roux, C. M. Shanahan, X. Trepat, D. Navajas, S. Garcia-Manyes, P. Roca-Cusachs, Force triggers yap nuclear entry by regulating transport across nuclear pores, Cell 171 (6) (2017) 1397–1410.e14.

[56] T. Wang, S. Hamilla, M. Cam, H. Aranda-Espinoza, S. Mili, Extracellular matrix stiffness and cell contractility control rna localization to promote cell migration, Nature Communications 8 (1) (2017) 896.

[57] M. V. Hunter, E. Joshi, S. Bowker, E. Montal, Y. Ma, Y. H. Kim, Z. Yang, L. Tuffery, Z. Li, E. Rosiek, A. Browning, R. Moncada, I. Yanai, H. Byrne, M. Monetti, E. de Stanchina, P.-J. Hamard, R. P. Koche, R. M. White, Mechanical confinement governs phenotypic plasticity in melanoma, Nature 647 (8089) (2025) 517–527.

[58] W.-H. Jung, E. Humann, J. M. Price, Y. Binenbaum, A. Haseki, S. Iyer, D. J. Mooney, Matrix viscoelasticity regulates dendritic cell migration and immune priming, bioRxiv (2025) 2025–09.

[59] Z. Outla, M. Prechova, K. Korelova, J. Gemperle, M. Gregor, Mechanics of cell sheets: plectin as an integrator of cytoskeletal networks, Open Biology 15 (1) (2025).

[60] A. Isomursu, K.-Y. Park, J. Hou, B. Cheng, M. Mathieu, G. A. Shamsan, B. Fuller, J. Kasim, M. M. Mahmoodi, T. J. Lu, et al., Directed cell migration towards softer environments, Nature materials 21 (9) (2022) 1081–1090.

[61] C. Huerta-López, A. Clemente-Manteca, D. Velázquez-Carreras, F. M. Espinosa, J. G. Sanchez, Álvaro Martínez-del Pozo, M. García-García, S. Martín-Colomo, A. Rodríguez-Blanco, R. Esteban-González, F. M. Martín-Zamora, L. I. Gutierrez-Rus, R. Garcia, P. Roca-Cusachs, A. Elosegui-Artola, M. A. del Pozo, E. Herrero-Galán, P. Sáez, G. R. Plaza, J. Alegre-Cebollada, Cell response to extracellular matrix viscous energy dissipation outweighs high-rigidity sensing, Science Advances 10 (46) (2024) eadf9758.

[62] H. T. Nia, H. Liu, G. Seano, M. Datta, D. Jones, N. Rahbari, J. Incio, V. P. Chauhan, K. Jung, J. D. Martin, V. Askoxylakis, T. P. Padera, D. Fukumura, Y. Boucher, F. J. Hornicek, A. J. Grodzinsky, J. W. Baish, L. L. Munn, R. K. Jain, Solid stress and elastic energy as measures of tumour mechanopathology, Nature Biomedical Engineering 1 (1) (2016) 0004.

[63] R. K. Jain, J. D. Martin, T. Stylianopoulos, The role of mechanical forces in tumor growth and therapy, Annual Review of Biomedical Engineering 16 (Volume 16, 2014) (2014).

[64] R. V. Kondapaneni, S. K. Gurung, P. S. Nakod, K. Goodarzi, V. Yakati, N. A. Lenart, S. S. Rao, Glioblastoma mechanobiology at multiple length scales, Biomaterials Advances 160 (2024) 213860.

[65] Y. Shou, X. Y. Teo, K. Z. Wu, B. Bai, A. R. K. Kumar, J. Low, Z. Le, A. Tay, Dynamic stimulations with bioengineered extracellular matrix-mimicking hydrogels for mechano cell reprogramming and therapy, Advanced Science 10 (21) (2023) 2300670.

[66] W. Fan, K. Adebowale, L. Váncza, Y. Li, M. F. Rabbi, K. Kunimoto, D. Chen, G. Mozes, D. K.-C. Chiu, Y. Li, J. Tao, Y. Wei, N. Adeniji, R. L. Brunsing, R. Dhanasekaran, A. Singhi, D. Geller, S. H. Lo, L. Hodgson, E. G. Engleman, G. W. Charville, V. Charu, S. P. Monga, T. Kim, R. G. Wells, O. Chaudhuri, N. J. Török, Matrix viscoelasticity promotes liver cancer progression in the pre-cirrhotic liver, Nature 626 (7999) (2024) 635–642.

[67] M. L. Cagigas, N. S. Bryce, N. Ariotti, S. Brayford, P. W. Gunning, E. C. Hardeman, Correlative cryo-et identifies actin/tropomyosin filaments that mediate cell–substrate adhesion in cancer cells and mechanosensitivity of cell proliferation, Nature Materials 21 (1) (2022) 120–128.

[68] J. A. Linke, L. L. Munn, R. K. Jain, Compressive stresses in cancer: characterization and implications for tumour progression and treatment, Nature Reviews Cancer 24 (11) (2024) 768–791.

[69] Y. Wang, K. F. Goliwas, P. E. Severino, K. P. Hough, D. Van Vessem, H. Wang, S. Tousif, R. P. Koomullil, A. R. Frost, S. Ponnazhagan, J. L. Berry, J. S. Deshane, Mechanical strain induces phenotypic changes in breast cancer cells and promotes immunosuppression in the tumor microenvironment, Laboratory Investigation 100 (12) (2020) 1503–1516.

[70] F. A. Ran, P. D. Hsu, J. Wright, V. Agarwala, D. A. Scott, F. Zhang, Genome engineering using the crispr-cas9 system, Nature protocols 8 (11) (2013) 2281–2308.

[71] C. Hagemann, D. Amend, A. F. Kessler, T. Linsenmann, R.-I. Ernestus, M. Löhr, High-efficiency transfection of glioblastoma cells and a simple spheroid migration assay, in: RNAi and Small Regulatory RNAs in Stem Cells: Methods and Protocols, Springer, 2017, pp. 63–79.

[72] A. Schaeffer, S. Buracco, M. Gazzola, M. Gelin, B. Vianay, C. de Pascalis, L. Blanchoin, M. Théry, Microtubule-driven cell shape changes and actomyosin flow synergize to position the centrosome, Journal of Cell Biology 224 (7) (2025) e202405126.

[73] M. D. Abràmoff, P. J. Magalhães, S. J. Ram, Image processing with imagej, Biophotonics international 11 (7) (2004) 36–42.

[74] D. Sage, Orientationj, Biomedical Image Group, EPFL, Switzerland (2012).

[75] L. R. Flores, M. C. Keeling, X. Zhang, K. Sliogeryte, N. Gavara, Lifeact-taggfp2 alters f-actin organization, cellular morphology and biophysical behaviour, Scientific reports 9 (1) (2019) 3241.

