## Supplementary Information for "Transient cytoskeletal anisotropy encodes short-term mechanical memory"

### 1. Supplementary Figures

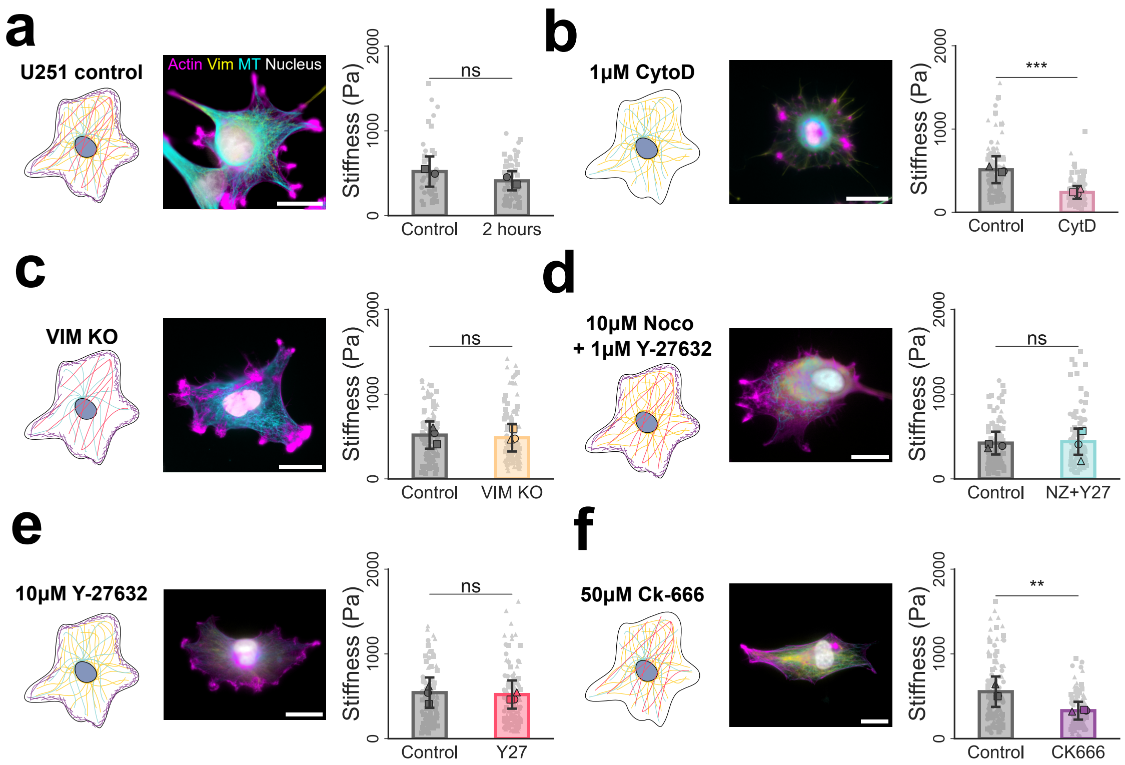


Figure S1. **Cytoskeletal structure determines glioblastoma cell mechanics.** Evaluation of changes in cortical cell stiffness due to alterations in the cytoskeleton structure. Representative actin, vimentin and microtubule organization and orientation for glioblastoma cells in **(a)** control conditions,**(c)** vimentin KO cells, and cells subjected to either **(b)** 1 $\mu$M Cytochalasin D (actin polymerization inhibitor), **(d)** 10 $\mu$M Nocodazole + 1 $\mu$M Y-27632 (tubulin depolymerization promoter), **(e)** 10 $\mu$M Y-27632 (ROCK-inhibitor, disrupts actin stress fibers) or **(f)** 50 $\mu$M Ck-666 (Arp2/3 complex inhibitor, disrupts actin cortex) are shown. The nanoindentation measurements were conducted before and after drug treatment. For the analysis of vimentin contribution, a comparison between control and vimentin KO cells is provided. A control case is included to show no significant changes in the baseline stiffness during measuring times. Scale bars: 20 $\mu m$. n= 100-150 cells analysis per condition from N=3 independent nanoindentation assays. Data are presented as mean ± SEM. Comparisons were made by unpaired t-test, ns $p > 0.05$ * $p \leq 0.05$, ** $p \leq 0.01$, **** $p \leq 0.0001$.

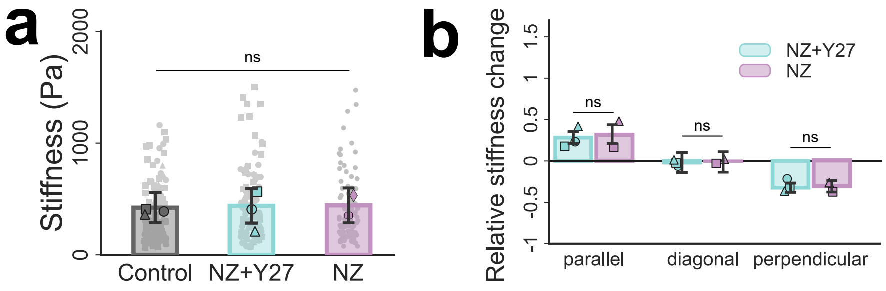


Figure S2. **Effect of microtubule depolymerization on glioblastoma cell mechanics with and without compensation of increased acto-myosin contractility.** Comparison of the mechanical response of glioblastoma cells treated with 10 $\mu$M Nocodazole (NZ) alone or combined with 1 $\mu$M Y-27632 (NZ+Y27). **(a)** Relaxed cortical stiffness measured by nanoindentation under non-actuated conditions **(b)** Relative stiffness change upon magneto-mechanical actuation applied parallel, diagonal, or perpendicular to the cell long axis. Relative stiffness change is calculated with respect to the relaxed state for each condition. Data are presented as mean $\pm$ SEM with individual data points overlaid. n = 40-60 cells per condition from N $\geq$ 2 independent experiments. Statistical significance was assessed using unpaired t-tests; ns p > 0.05.

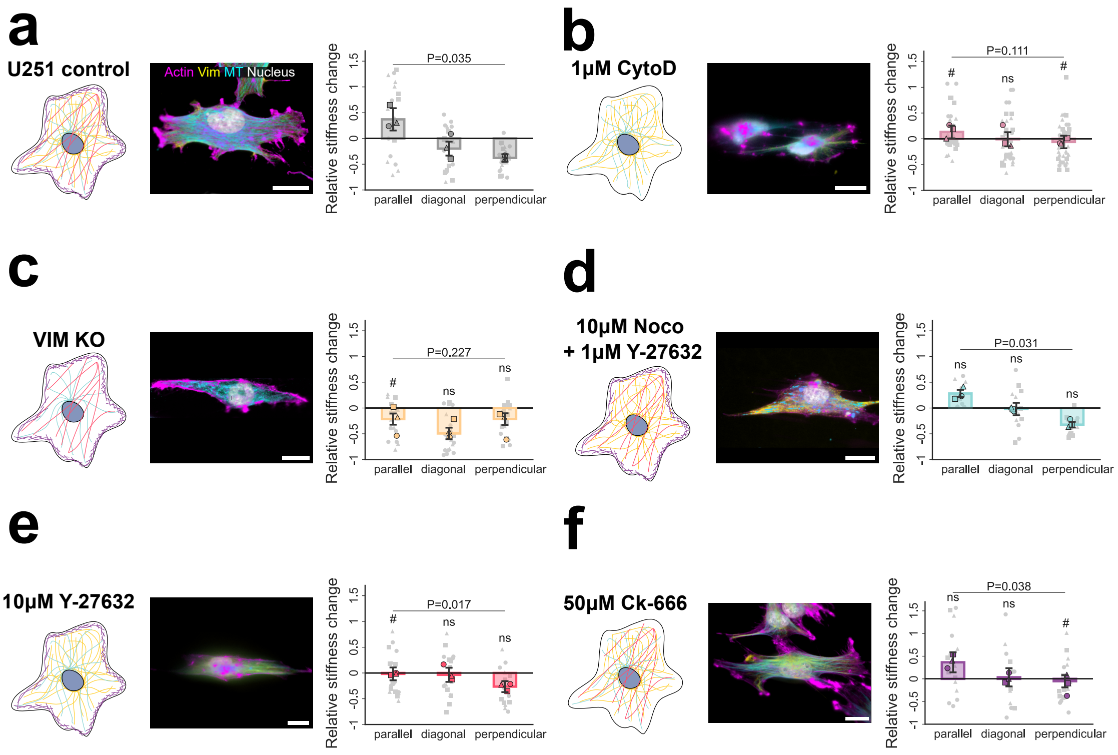


Figure S3. **Cytoskeletal reorganization due to mechanical actuation determines changes in apparent stiffness.** Representative actin, vimentin and microtubule organization for actuated glioblastoma cells and relative stiffness change between relaxed and stretching actuation states for cells oriented parallel, diagonal, and perpendicular to the actuation direction in **(a)** control conditions, **(c)** vimentin KO cells, and cells subjected to either **(b)** 1 $\mu$M Cytochalasin D, **(d)**10 $\mu$M Nocodazole + 1 $\mu$M Y-27632, **(e)** 10 $\mu$M Y-27632 or **(f)** 50 $\mu$M Ck-666. Scale bars: 20 $\mu m$. n=40-70 cells analyzed per condition for N=3 independent nanoindentation assays. Data are presented as mean $\pm$ SEM. Group comparisons were performed using two-way ANOVA with multiple comparisons. Comparisons with the corresponding control case for each condition and orientation are marked by hash symbols (#), with ns p > 0.05, # p$\leq$0.05.

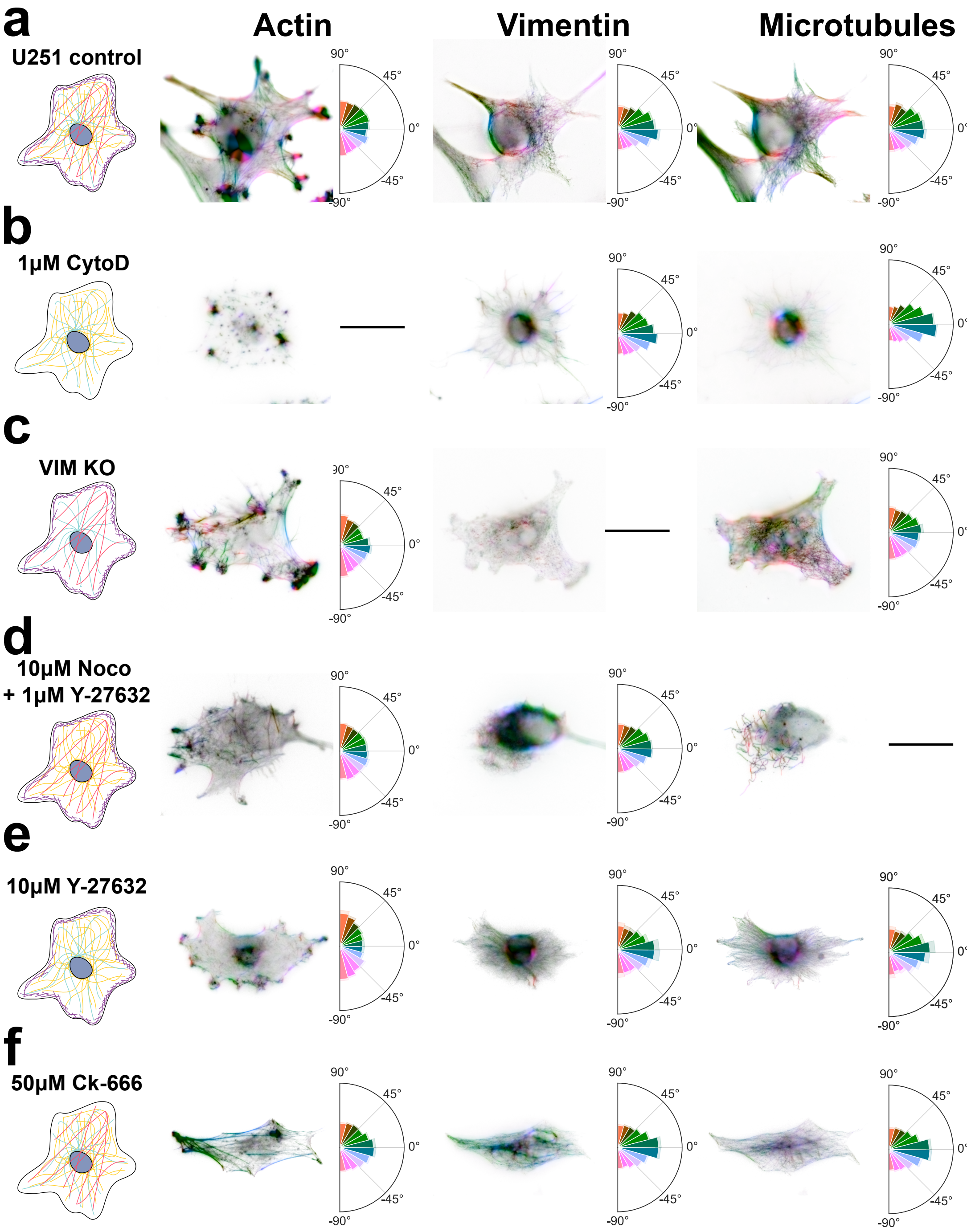


Figure S4. **Cytoskeletal organization under relaxed condition and perturbations.** Representative actin, vimentin and microtubule organization and corresponding fiber orientation distributions in relaxed glioblastoma cells under **(a)** control conditions, **(b)** 1 $\mu$M Cytochalasin D, **(c)** vimentin knockout (VIM KO), **(d)** 10 $\mu$M Nocodazole + 1 $\mu$M Y-27632, **(e)** 10 $\mu$M Y-27632, and **(f)** 50 $\mu$M Ck-666. Fiber orientations are color-coded according to their angular direction, and polar histograms represent the average fiber distribution for each cytoskeletal component and condition. Scale bars: 20 $\mu$m. n=10–20 cells analyzed per condition for cytoskeletal analysis for N=3 independent experiments.

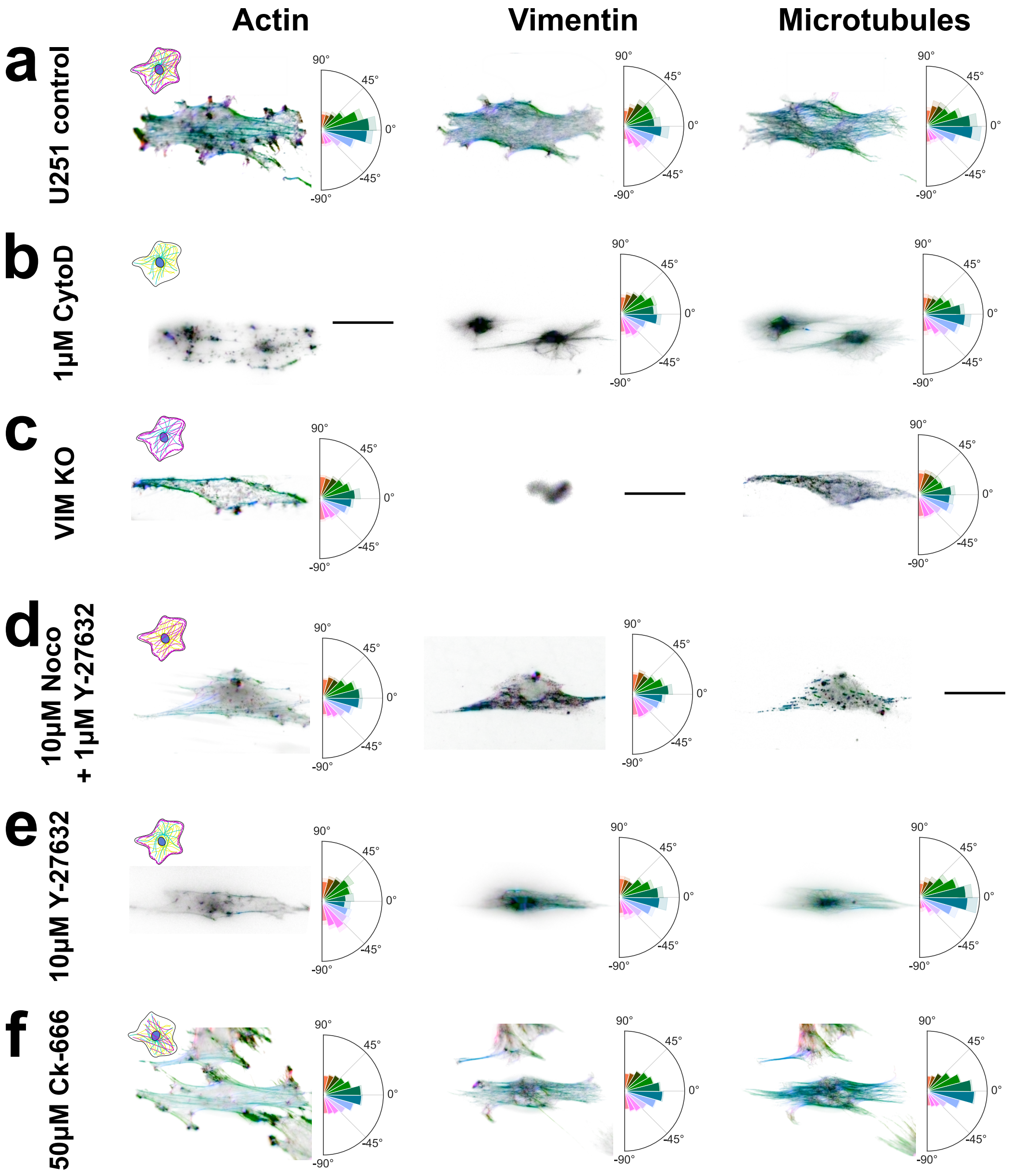


Figure S5. **Cytoskeletal organization under mechanical actuation and perturbations.** Representative actin, vimentin and microtubule organization and corresponding fiber orientation distributions in actuated glioblastoma cells under **(a)** control conditions, **(b)** 1 $\mu$M Cytochalasin D, **(c)** vimentin knockout (VIM KO), **(d)** 10 $\mu$M Nocodazole + 1 $\mu$M Y-27632, **(e)** 10 $\mu$M Y-27632, and **(f)** 50 $\mu$M Ck-666. Fiber orientations are color-coded according to their angular direction, and polar histograms represent the average fiber distribution for each cytoskeletal component and condition. Scale bars: 20 $\mu$m. n=10–20 cells analyzed per condition for cytoskeletal analysis for N=3 independent experiments.

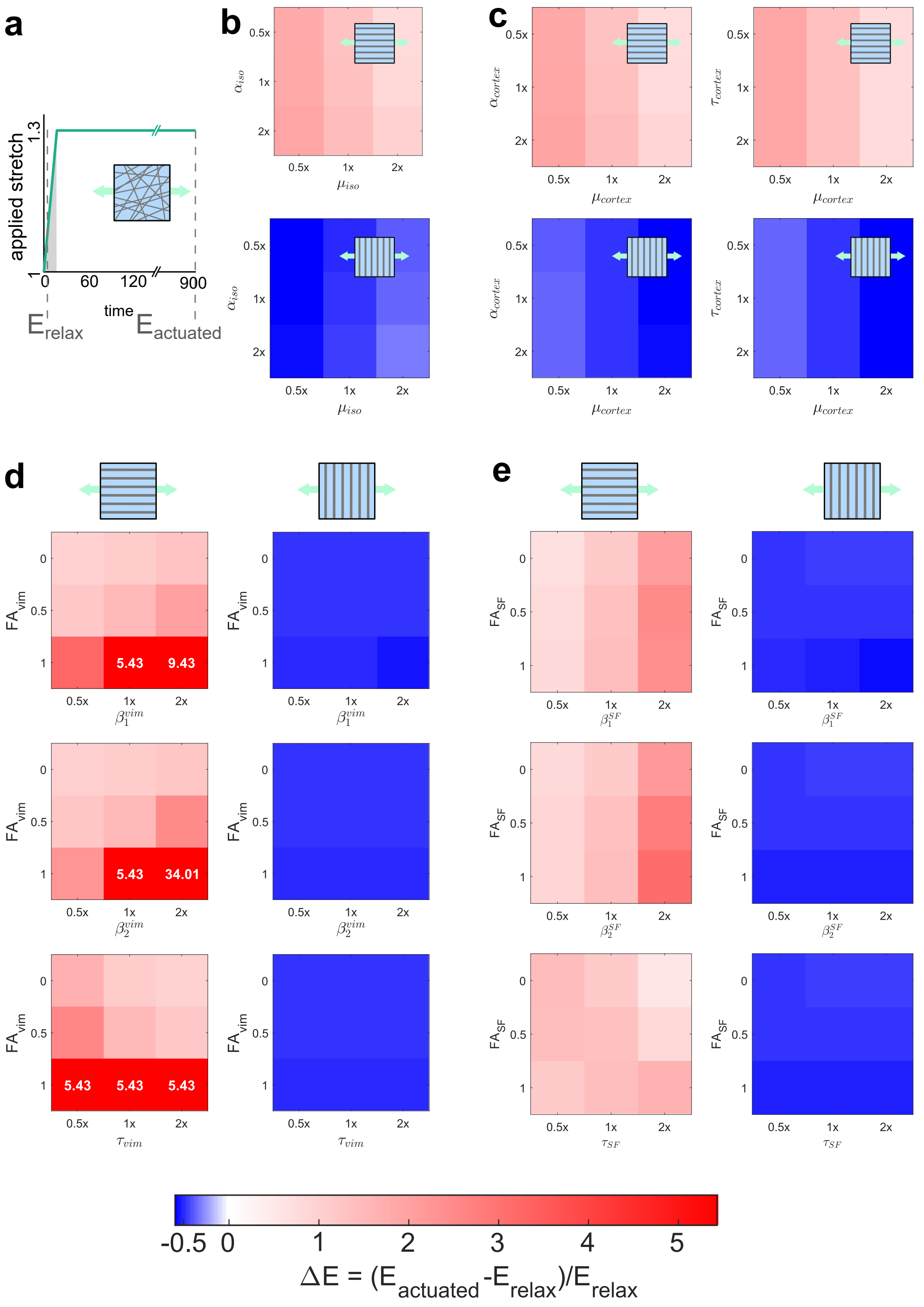


Figure S6. **Parametric sensitivity analysis of deformation-mode-specific stiffness changes.** **(a)** Schematic of the actuation protocol used in the simulations. Cells were subjected to a uniaxial actuation, and the relative stiffness change was computed between actuated and relaxed configurations. **(b–e)** Relative stiffness change, for actuation applied parallel or perpendicular to the fiber orientation. Heatmaps show the effect of varying model parameters to 0.5×, 1×, or 2× their baseline value for different initial fractional anisotropy values (FA = 0, 0.5, 1) for parameters associated with **(b)** the isotropic contribution, **(c)** the actin cortex network, **(d)** the vimentin network, **(e)** the stress fiber network. Color scale indicates the magnitude of the relative stiffness change, with blue representing a relative softening, white a maintenance of stiffness and red a relative stiffening. All simulations have been conducted taking the model parameters collected in Table S1 as reference. Then, specific parameters have been modulated according to the parametric analysis performed in each case.

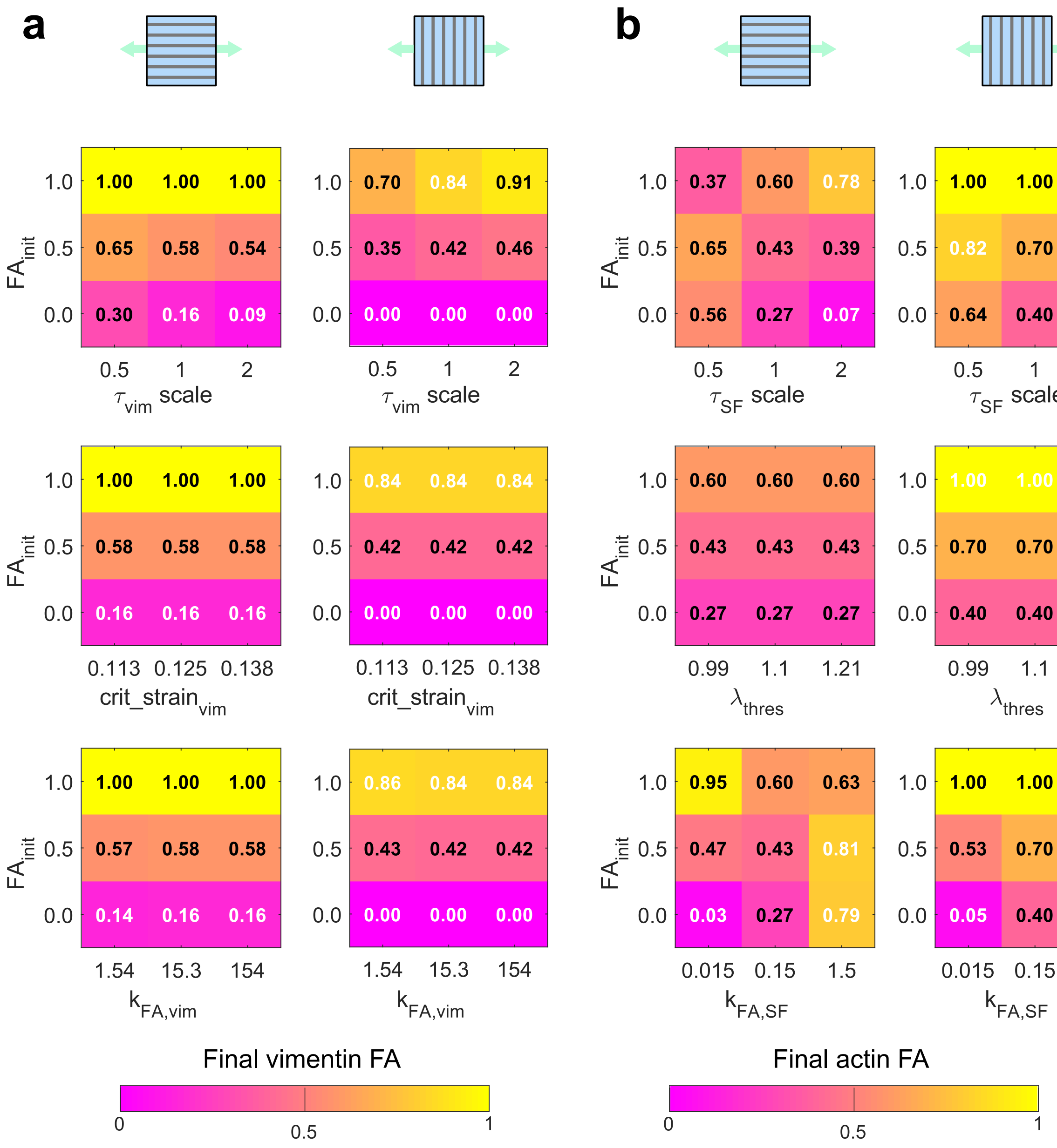


Figure S7. **Parametric sensitivity analysis of final network anisotropy.** Parametric analysis of the final fractional anisotropy (FA) obtained after actuation, as a function of the initial FA and parameters governing fiber dynamics. Heatmaps show FA for parameters scaled around their baseline values. **(a)** Final vimentin FA as a function of initial FA and vimentin fiber dynamics parameters, including characteristic reorganization time, critical strain, and anisotropic reinforcement coefficient. **(b)** Final actin FA as a function of initial FA and actin fiber dynamics parameters, including stress fiber characteristic time, activation threshold and anisotropic reinforcement coefficient. Color scale indicates the magnitude of the final fractional anisotropy. All simulations have been conducted taking the model parameters collected in Table S1 as reference. Then, specific parameters have been modulated according to the parametric analysis performed in each case.

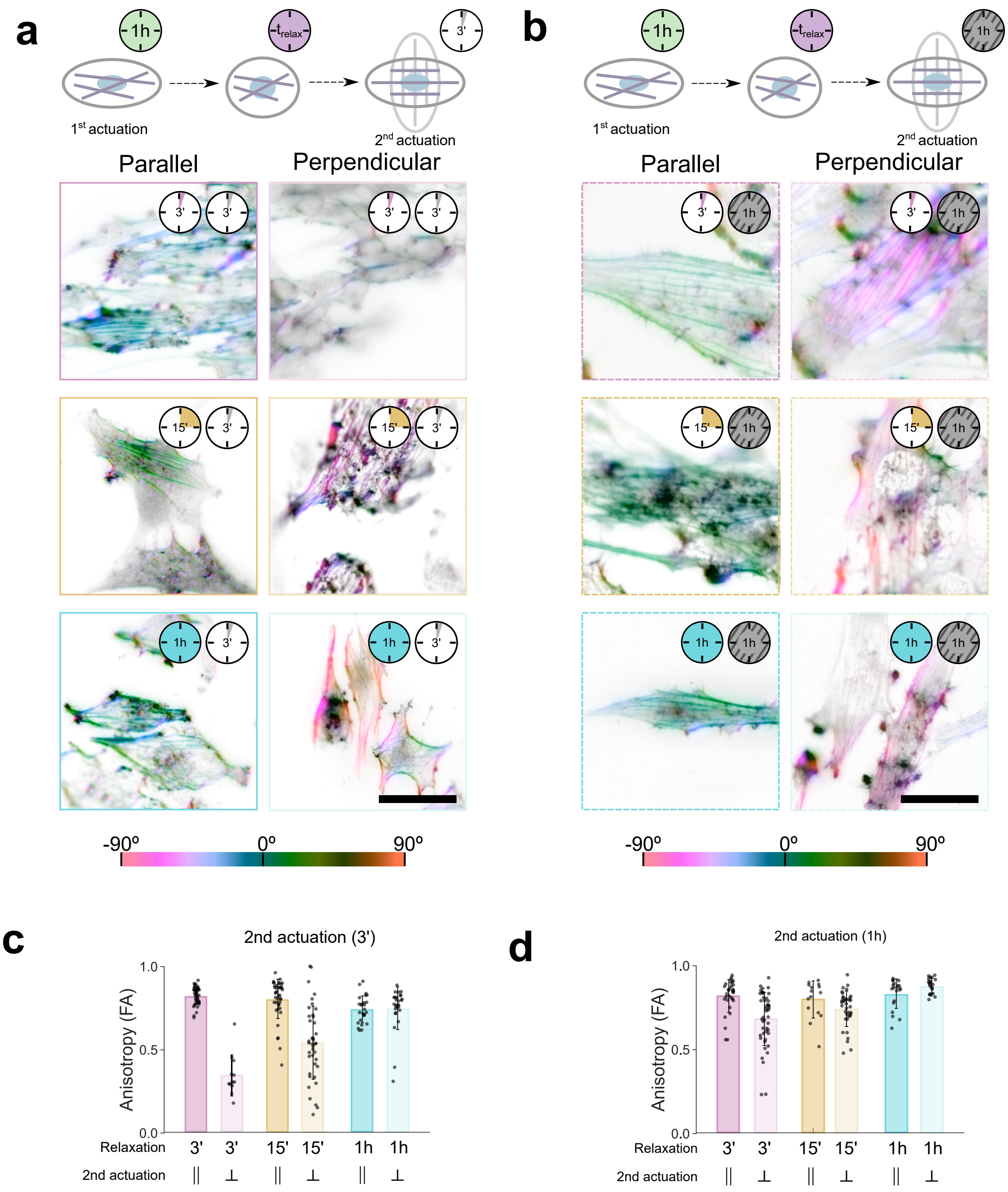


Figure S8. **Cytoskeletal anisotropy following mechanical memory protocols with short and long second actuation.** **(a)** Representative images of orientation of actin fibers after memory protocols in which the second actuation was applied for 3 min. Cells were subjected to an initial actuation (1 h), followed by a relaxation period, and then a second actuation applied either parallel or perpendicular to the first loading direction. Color scale represents fiber orientation angle. **(b)** Representative orientation maps following the same memory protocols with a second actuation duration of 1 h. **(c)** Final fractional anisotropy (FA) quantified for the conditions shown in (a). **(d)** Final FA for the conditions shown in (b). Data are presented as mean ± SEM with individual data points overlaid.

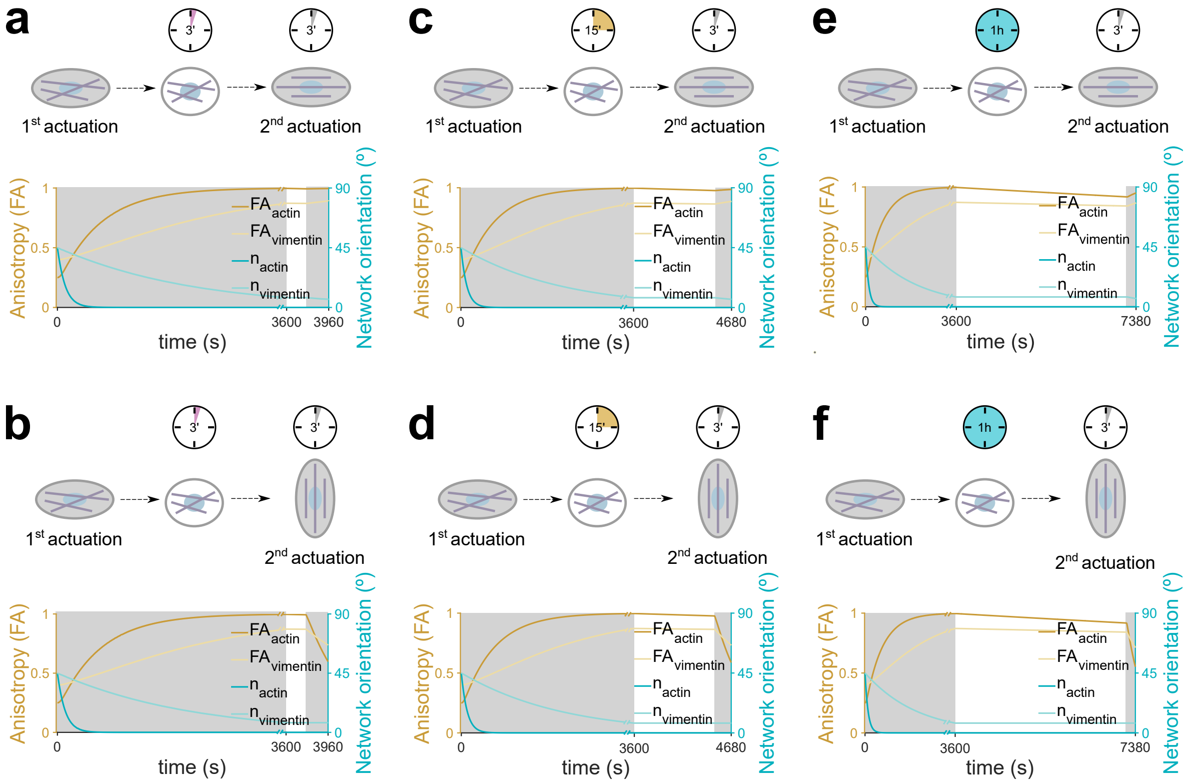


Figure S9. **Model-predicted evolution of cytoskeletal anisotropy and principal orientation during mechanical memory protocols with second actuation of 3 min. (a–f)** Temporal evolution of fractional anisotropy (FA) and principal network orientation predicted by the computational model during memory protocols in which the second actuation was applied for 3 min. Schematics above each panel indicate the loading sequence and relative orientation of first and second actuation. The shaded region denotes the actuation periods. Panels correspond to different relaxation durations between the first and second actuation and to parallel or perpendicular reloading conditions, as indicated schematically.

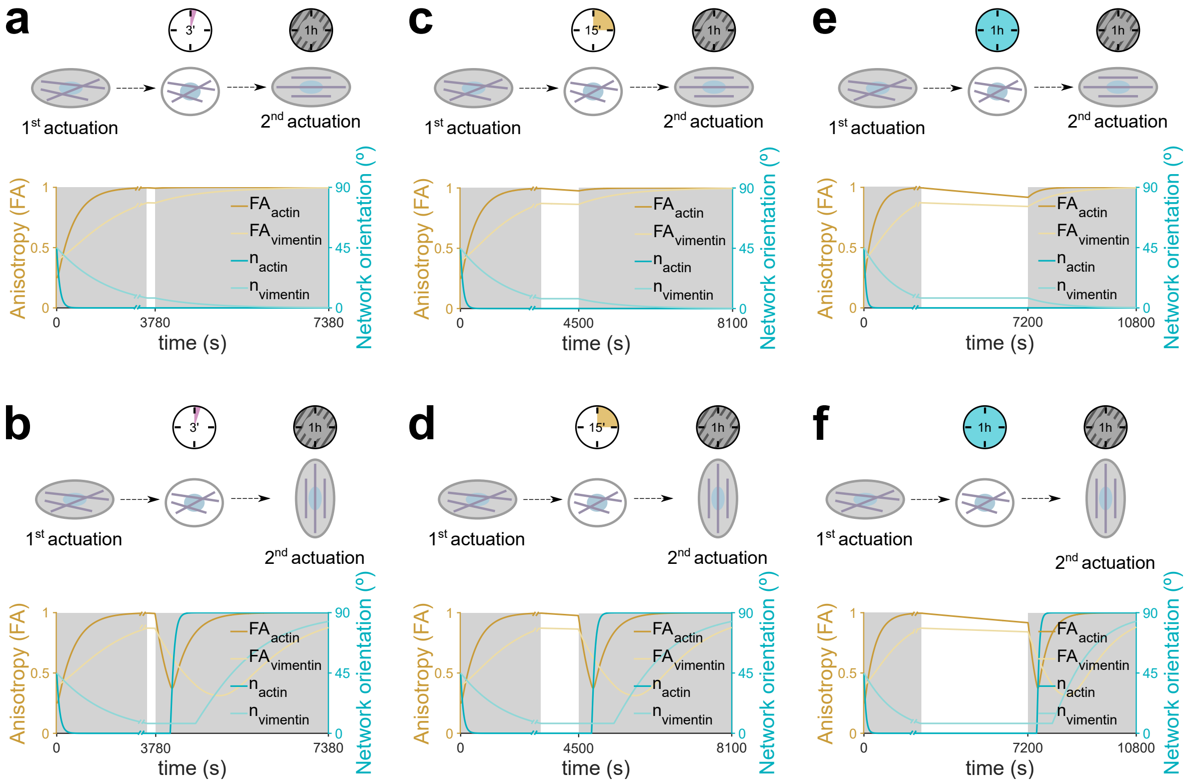


Figure S10. **Model-predicted evolution of cytoskeletal anisotropy and principal orientation during mechanical memory protocols with second actuation of 1 h. (a–f)** Temporal evolution of fractional anisotropy (FA) and principal network orientation predicted by the computational model during memory protocols in which the second actuation was applied for 1 h. Schematics above each panel indicate the loading sequence and relative orientation of first and second actuation. The shaded region denotes the actuation periods. Panels correspond to different relaxation durations between the first and second actuation and to parallel or perpendicular reloading conditions, as indicated schematically.

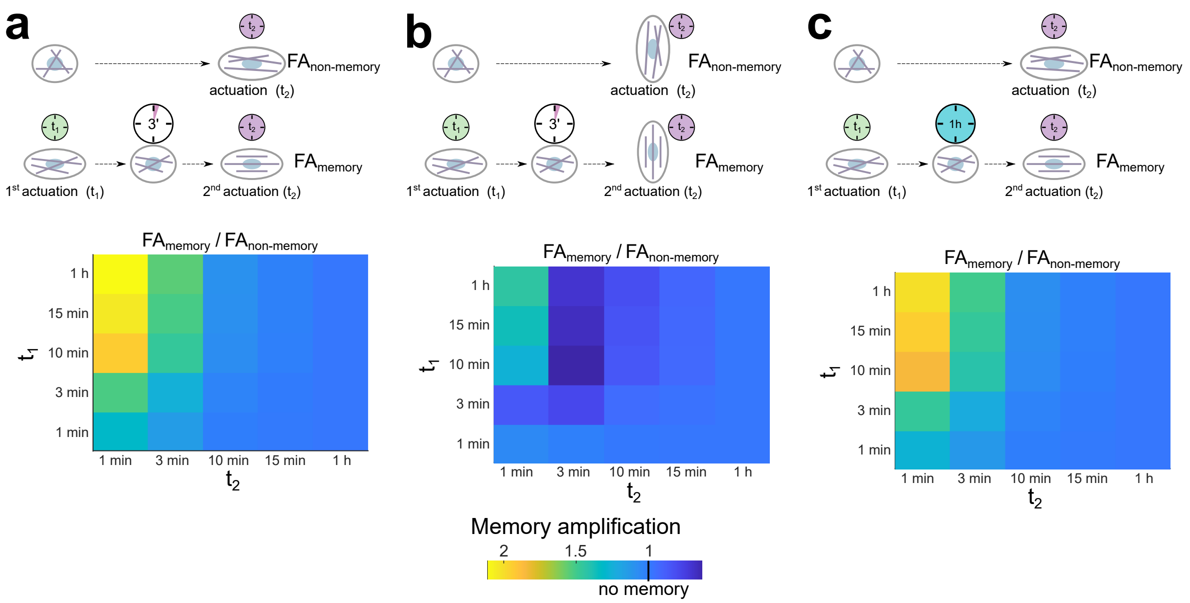

Figure S11. **Model-predicted amplification of anisotropic memory across loading durations.** Heatmaps of the memory ratio, defined as the final fractional anisotropy after the full two-actuation protocol ($FA_{memory}$) normalized by the anisotropy obtained under the corresponding single-actuation control ($FA_{non-memory}$). The first actuation duration is denoted $t_1$ and the second actuation duration $t_2$. **(a)** Parallel re-actuation with 3 min relaxation between the first and second actuation. **(b)** Perpendicular re-actuation with 3 min relaxation. **(c)** Parallel re-actuation with 1 h relaxation. Color scale indicates memory amplification (>1) or absence of memory (≈1).

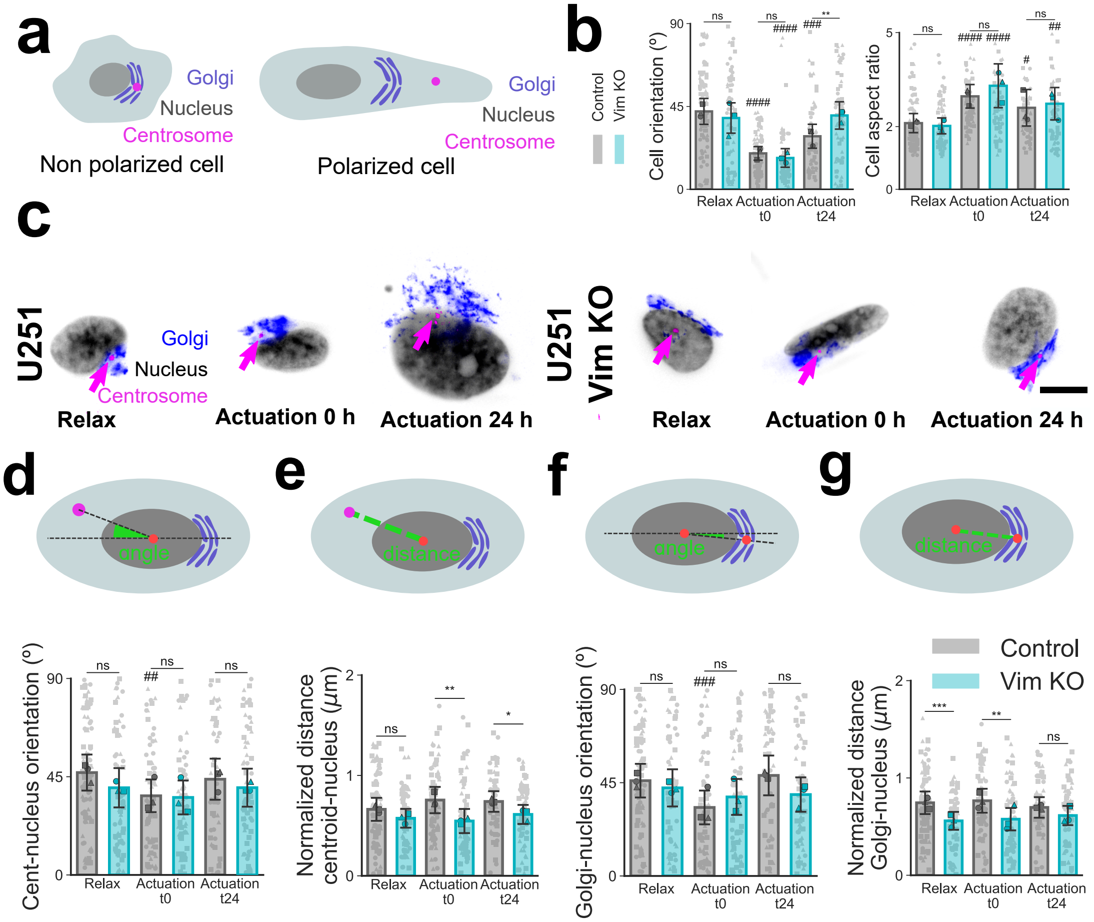


Figure S12. **Analysis of cell polarization under mechanical actuation.** **(a)** Schematics of non-polarized (rounded nucleus and cell morphology, centrosome and Golgi located close to the nucleus with a random angular position) compared to a polarized cell (cell with high aspect ratio, nucleus located towards the back of the cell with Golgi and centrosome in front). **(b)** Cell orientation and aspect ratio for control (grey) and vimentin KO (blue) cells before, right after, and 24 h after actuation. In the latter case, the stretching actuation is kept over 24 h. **(c)** Representative images of the angle between the nucleus-centrosome axis and the direction of the stretching actuation for control and vimentin KO cells before, right after, and 24 h after actuation. Scale bars: 10 $\mu m$. Analysis of polarization parameters for control (gray) and vimentin KO (blue) cells before, right after and 24 h after actuation: **(d)** angle between the nucleus-centrosome axis and the direction of actuation, **(e)** normalized distance between the centrosome and the nucleus with respect to the nucleus radius, **(f)** angle between the nucleus-Golgi axis and the direction of actuation, and **(g)** normalized distance between the Golgi and the nucleus with respect to the nucleus radius. n=50-100 analyzed cells per condition and time-point from N=3 independent experiments. In the explanatory cartoons, the red dots represent the center point for the nucleus, and Golgi is used to determine the distances and angles calculated. Data are presented as mean ± SEM. Group comparisons were performed by two-way ANOVA with multiple comparisons, Comparisons between control and vimentin KO cells are represented by asterisk symbols (*) and comparisons with the corresponding relaxed state for each condition are marked by hash symbols (#). For all symbols, * $p \leq 0.05$, ** $p \leq 0.01$, *** $p \leq 0.001$, **** $p \leq 0.0001$.

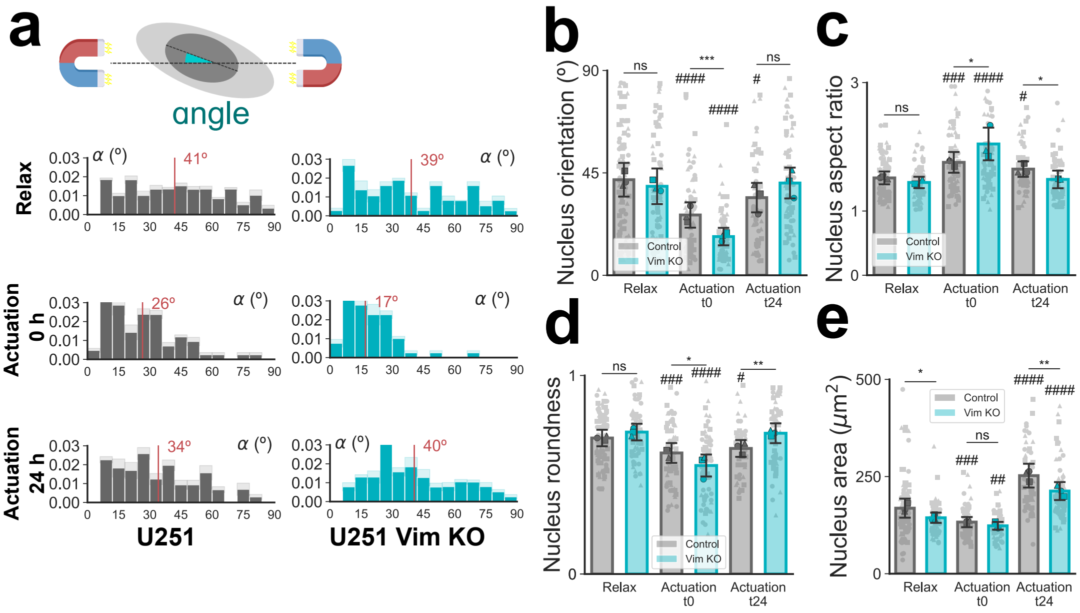


Figure S13. **Cell nucleus deforms and aligns in the direction of actuation in a vimentin-dependent manner.** **(a)** Histograms of the angle between the cell nucleus main axis orientation and the direction of stretching actuation for control (grey) and vimentin KO (blue) cells before, right after, and 24 h after actuation. Analysis of morphological parameters of the nucleus for control (grey) and vimentin KO (blue) cells before, right after, and 24 h after actuation: **(b)** Relative angle between the nucleus orientation and the actuation direction, **(c)** nucleus aspect ratio (ratio between the minimum and maximum Feret diameters), **(d)** nucleus area, and **(e)** nucleus roundness. n=50-100 analyzed cells per condition and time-point from N=3 independent experiments. Data are presented as mean ± SEM. Group comparisons were performed by two-way ANOVA with multiple comparisons, Comparisons between control and vimentin KO cells are represented by asterisk symbols (*) and comparisons with the corresponding relaxed state for each condition are marked by hash symbols (#). For all symbols, * $p \leq 0.05$, ** $p \leq 0.01$, *** $p \leq 0.001$, **** $p \leq 0.0001$.

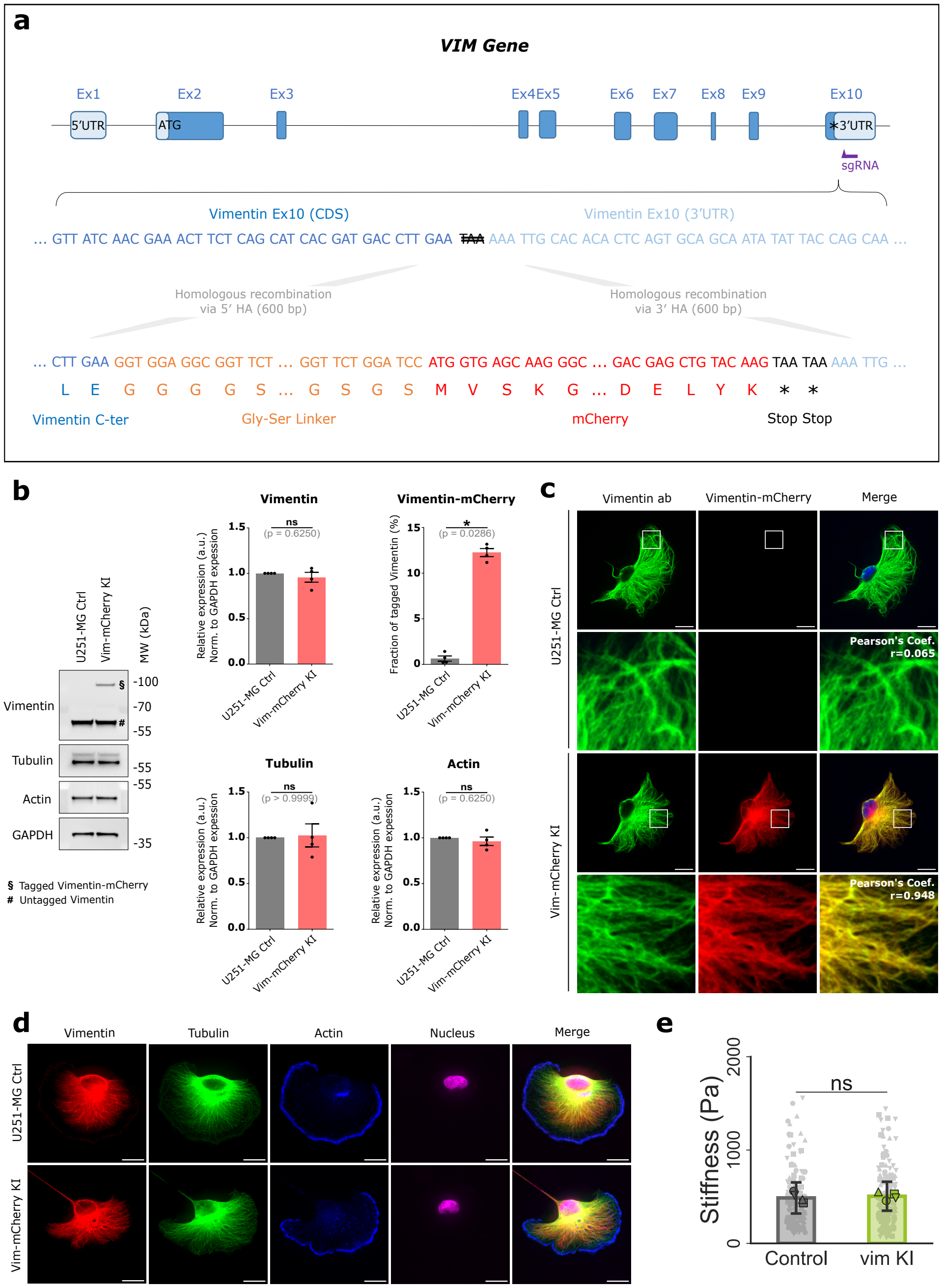


Figure S14. **Generation and characterization of the Vimentin–mCherry Knock-In cell line.** **(a)** CRISPR–Cas9 targeting strategy at the endogenous VIM locus. The sgRNA (purple) targets exon 10 near the stop codon. The donor template contains 5' and 3' homology arms corresponding to the exon 10 coding sequence (CDS, dark blue) and the 3' untranslated region (3' UTR, light blue), and an insert comprising a Gly–Ser linker (orange), the mCherry coding sequence (red), and two stop codons (black), resulting in an in-frame C-terminal fusion of Vimentin with mCherry expressed under endogenous regulatory elements. **(b)** Western blot of parental U-251 MG control (Ctrl) and Vimentin–mCherry Knock-In (KI) cell lysates probed for Vimentin, Tubulin, Actin, and GAPDH. Quantification of protein levels normalized to GAPDH and the fraction of tagged Vimentin relative to total Vimentin are shown (mean ± SEM; N = 4 independent experiments; Wilcoxon test for relative expression; Mann–Whitney test for tagged fraction). **(c)** Immunofluorescence of Ctrl and KI cells stained for Vimentin (green) and imaged for endogenous mCherry fluorescence (red). Colocalization in representative images was quantified using the Pearson correlation coefficient (N = 3 independent experiments, n = 10 cells per condition, scale bar: 20$\mu$m). **(d)** Immunofluorescence staining of Vimentin, Tubulin, F-actin (phalloidin), and nuclei (Hoechst) in Ctrl and KI cells, showing comparable cytoskeletal organization (N = 3 independent experiments, n = 10 cells per condition, scale bar: 20$\mu$m). **(e)** Nanoindentation data of control (U251 -MG Ctrl) and Vim KI (Vim-mCherry KI) under relaxed conditions. Data are presented as mean ± SEM with individual data points overlaid. Statistical comparisons were performed by unpaired t-test; ns p>0.05.

### 2. Supplementary Tables

Table S1: Summary of model variables and parameters with their biological interpretation. All experimental results in this study are expressed in terms of relative variations of cell stiffness and structural evolution quantified through fractional anisotropy (FA) and principal fiber orientation (**n**). Accordingly, the internal variables describing cytoskeletal organization and the reported mechanical variations are dimensionless quantities. To ensure consistency with this relative formulation, stress contributions in the constitutive model are non-dimensionalized. All stiffness-related parameters are scaled with respect to a reference baseline value of 100 associated with the isotropic background contribution (representing the effective cytosolic and non-explicit cellular components). Parameter magnitudes are selected based on experimental characterization and values reported in the literature, and then normalized relative to this baseline. Characteristic time scales governing remodeling kinetics retain their physical dimensions to allow meaningful comparison with experimentally observed temporal evolution. This normalization strategy does not affect model predictions in relative terms, but improves robustness and interpretability in view of intrinsic cellular variability.

| Symbol | Name | Mechanical view | Biological view | Effect | Value / range | Source |
|---|---|---|---|---|---|---|
| **Network descriptors (shared across networks)** | | | | | | |
| $\varphi$ | Fibrosity / density | Weight of a network's stress contribution | Fraction of actin/vimentin in load-bearing form (bundled or cortical) | $\uparrow\varphi \Rightarrow$ larger stress share from that network | *Evolving* (0–1) | State variable |
| FA | Fractional anisotropy | Orientation order parameter | Degree of filament alignment (0 isotropic, 1 highly aligned) | $\uparrow$FA $\Rightarrow$ stronger structural anisotropy | *Evolving* (0–1) | State variable |
| $\mathbf{n}$ | Preferred direction (in-plane) | Principal network direction used in $G$ and $I_4$ | Network's dominant stress/shape direction that fibers predominantly follow | Sets direction along which stiffness and remodeling are predominant | *Evolving* (0–90°) | State variable |
| $\kappa$, $G$ | Dispersion and structured tensor | $\kappa$ encodes spread around $\mathbf{n}$; $G = \kappa\mathbf{I} + (1-3\kappa)\,\mathbf{n}\otimes\mathbf{n}$ | Descriptors of the network organization based on FA and n | Lower spread $\Rightarrow$ larger $I_4$ and anisotropy | *Evolving* (0–1/3) | Determined by FA and n |
| **Isotropic background (cytoplasm + non-explicit components)** | | | | | | |
| $\mu_{\text{iso}}$ | Isotropic shear modulus | Baseline shear stiffness | Passive stiffness of cytoplasm + non-explicit filaments | $\uparrow\ \mu_{\text{iso}} \Rightarrow$ higher baseline modulus | 100 | Baseline for normalization |

*(continued on next page)*

(continued from previous page)

| Symbol | Name | Mechanical meaning | Biological interpretation (biologist-friendly) | Effect on behavior | Value / range | Source / where set |
|---|---|---|---|---|---|---|
| $\alpha_{\mathrm{iso}}$ | Isotropic non-linearity exponent | Controls strain-stiffening via $I_1^{\alpha-1}$ | Cytoplasm stiffening at large strains | $\uparrow\ \alpha_{\mathrm{iso}} \Rightarrow$ sharper stiffening curve | 1 | Assumed linear |
| **Vimentin network** | | | | | | |
| $\mu_{\mathrm{vim}}$ | Vimentin shear modulus | Baseline shear stiffness in vimentin fiber stress (tension-only via $I_4$ gate) | Stiffness of actin bundles along their axis | $\uparrow\ \mu_{\mathrm{vim}} \Rightarrow$ stiffer response in vim direction | 20 | [1] |
| $\beta_{\mathrm{vim}}$ | Vimentin non-linearity exponent | Exponential stiffening | Recruitment/strain-stiffening of IF under load | $\uparrow\ \beta_{\mathrm{vim}} \Rightarrow$ stronger strain stiffening | 3 | [1, 2] |
| $\tau_{\mathrm{vim}}$ | Vimentin characteristic time | Baseline vimentin remodeling time | How fast vimentin reorganizes | Larger $\tau \Rightarrow$ slower alignment/recovery | 1 000 -10 000 s | [3] |
| $\lambda_{\mathrm{crit}}$ | Stretch threshold (vimentin) | Gate for decrease in vimentin characteristic time | Minimum effective stretch to increase vimentin reorganization speed | Faster vimentin reorganization when $\lambda > \lambda_{\mathrm{crit}}$ | 12.5% | [4] |
| **Stress fibers (actin bundles)** | | | | | | |
| $\mu_{\mathrm{SF}}$ | SF shear modulus | Baseline shear modulus in SF (tension-only via $I_4$ gate) | Stiffness of actin bundles along their axis | $\uparrow\ \mu_{\mathrm{SF}} \Rightarrow$ higher axial stiffness | 80 | [5] |
| $\beta_{\mathrm{SF}}$ | SF nonlinearity | Exponential stiffening | Bundle recruitment/strain-stiffening | $\uparrow\ \beta_{\mathrm{SF}} \Rightarrow$ sharper high-strain rise | 1.5 | [6] |
| $\lambda_{\mathrm{thres}}$ | Stretch threshold (SF) | Gate for polymerization onset in kinetics | Minimum effective stretch to start SF build-up | Higher threshold $\Rightarrow$ delayed SF growth | 1.1 | [7] |
| $k^{\mathrm{SF}}_{\mathrm{pol},0}$ | SF polymerization constant | Baseline polimerization rate | Baseline rate at which new SF are added to the network | $\uparrow\ k^{\mathrm{SF}}_{\mathrm{pol},0} \Rightarrow$ faster bundle accumulation | $1/450\, s^{-1}$ | [3] |
| $k^{\mathrm{SF}}_{\mathrm{depol},0}$ | SF depolymerization constant | Baseline depolimerization rate | Baseline rate at which SF are depolimerized into G-actin | $\uparrow\ k^{\mathrm{SF}}_{\mathrm{depol},0} \Rightarrow$ faster bundle destruction | $10^{-4} s^{-1}$ | [8] |

(continued on next page)

*(continued from previous page)*

| Symbol | Name | Mechanical meaning | Biological interpretation (biologist-friendly) | Effect on behavior | Value / range | Source / where set |
|---|---|---|---|---|---|---|
| $\tau_{\mathrm{act}}$ | SF re-orientation time | Timescale for $\mathbf{n}_{\mathrm{SF}}$ rotation | Speed at which bundles align with load | $\downarrow\ \tau\ \Rightarrow$ quicker alignment | 60 s | [9] |
| $\tau_{\mathrm{act}}^{\mathrm{int}}$ | SF integration time | Transfer timescale for new aligned fibers into $G$ | Consolidation of new growth into aligned network | $\downarrow\ \tau\ \Rightarrow$ faster FA increase | 60 s | [9] |
| **Cortex (actin cortex sheet)** | | | | | | |
| $\mu_{\mathrm{c}}$ | Actin cortex shear modulus | Baseline actin cortex stiffness | Load-bearing of the actin cortex surrounding the cell | $\uparrow\ \mu_{\mathrm{c}}\ \Rightarrow$ higher background stiffness | 50 | [10, 11] |
| $\alpha_{\mathrm{c}}$ | Cortex nonlinearity exponent | Controls cortical strain-stiffening via $I_1^{\alpha-1}$ | How quickly the cortex stiffens as it stretches | $\uparrow\ \alpha_{\mathrm{c}}\ \Rightarrow$ sharper stiffening | 1 | [12] |
| $k_0^{\mathrm{c}}$ | Actin cortex turnover constant | Baseline turnover rate | Baseline rate at which new fibers are polymerized and depolymerized in the cortex | $\uparrow\ k_0^{\mathrm{c}} \Rightarrow$ faster actin cortex turnover | $0.01\ s^{-1}$ | [13] |
| $\lambda_{\mathrm{crit}}$ | Buckling onset stretch | Critical stretch along $\mathbf{n}_{\mathrm{cortex}}$ for instability onset | Compression threshold where cortex begins to lose load capacity | Higher $\lambda_{\mathrm{crit}} \Rightarrow$ easier buckling | 0.9 | [14] |
| $m_{\mathrm{buck}}$ | Buckling multiplier (0–1) | Scales cortical stress: 1 traction; $< 1$ compression | Effective loss of cortical load-bearing when compressed | $\downarrow$ to 0 with strong compression; 1 in traction | *Evolving* (0–1) | Computed (damage law) |

| Primer | Sequence |
|---|---|
| HA 5'_fwd | GCAGCTACATGAAACAGC |
| HA 5'_rev | cgcctccaccTTCAAG GTCATCGTGATG |
| Linker-mCherry_fwd | tgaccttgaaGGTGGAGGCGGTTCTGG |
| Linker-mCherry_rev | tgtgcaatttTTATTACTTGTACAGCTCGTCCATGCC |
| HA 3'_fwd | caagtaataaAAATTGCACACACTCAGTG |
| HA 3'_rev | TCTGTTTGTTATGATTAAAATCAC |

Table S2: Primer sequences used for the generation of a U251-MG Vimentin–mCherry Knock-In Cell Line (Vim-mCherry KI)

| Antibody | Dilution | Source | Reference |
|---|---|---|---|
| anti-vimentin | 1:1000 | Santa-Cruz | sc-6260 |
| anti-tubulin | 1:1000 | Sigma-Aldrich | MAB1864 |
| anti-GAPDH | 1:10000 | Sigma-Aldrich | MAB374 |
| anti-mouse HRP-conjugated | 1:10000 | Jackson ImmunoResearch | 715-035-151 |
| anti-rat HRP-conjugated | 1:10000 | Jackson ImmunoResearch | 112-035-003 |
| anti-actin hFAB rhodamine antibody | 1:10000 | Bio- Rad | 12004164 |
| Alexa Fluor 488 donkey anti-mouse | 1:1000 | Jackson ImmunoResearch | 715-295-150 |
| rhodamine (TRITC) goat anti-rat | 1:1000 | Jackson ImmunoResearch | 112-025-003 |
| Alexa Fluor 488 anti-mouse | 1:1000 | Jackson ImmunoResearch | 715-545-151 |
| Alexa Fluor 488 anti-rat | 1:1000 | Jackson ImmunoResearch | 712-545-153 |
| Hoechst 34580 | 1:10,000 | Invitrogen | |
| Alexa 647 phalloidin | 1:2000 | Abcam | lot GR3256003-6 |

Table S3: Antibodies used for Western blot and immunofluorescence imaging.

### 3. Glioblastoma constitutive model

#### *3.1. Model overview and rationale*

Model overview and rationale: We formulate a continuum-mechanics model of glioblastoma cells that is explicitly grounded on microstructural mechanisms associated with cytoskeleton dynamics. The model is motivated and supported by a consistent set of experimental observations, including (i) structural imaging of the cytoskeleton by fluorescence microscopy and (ii) mechanical testing by nanoindentation. These experimental assays have been conducted under different steady-state and transient mechanical conditions. In addition, perturbation experiments (i.e., modulating specific cytoskeleton components) provide direct evidence on how specific cytoskeletal components regulate the mechanical response. Across these assays, we consistently observe a strong dependence of the measured structural and mechanical response on the actin network and on vimentin intermediate filaments, whereas the contribution of microtubules appears comparatively less dominant under the loading conditions considered here. This motivates a constitutive description in which the overall response emerges from the interaction of distinct microstructural phases, with special emphasis on actin and vimentin. Accordingly, we describe the cell as a composite continuum whose total

Helmholtz free-energy density is additively decomposed into three main contributions,

$$\Psi = \Psi_{\rm iso} + \Psi_{\rm vim} + \Psi_{\rm act}. \tag{1}$$

The isotropic term represents the contribution of the cytosolic matrix and other constituents that are not explicitly modeled as filamentous networks (with microtubules embedded in this effective response), and is assumed to depend solely on the deformation gradient $\mathbf{F}$,

$$\Psi_{\rm iso} = \Psi_{\rm iso}(\mathbf{F}). \tag{2}$$

The remaining terms account for the filamentous cytoskeletal networks. In particular, $\Psi_{\rm vim}$ represents the contribution of the vimentin intermediate filament network, while $\Psi_{\rm act}$ represents the contribution of the actin network. Both are modeled as anisotropic energy contributions associated with oriented microstructural architectures and are therefore assumed to depend not only on the deformation gradient $\mathbf{F}$, but also on additional sets of internal variables,

$$\Psi_{\rm vim} = \Psi_{\rm vim}(\mathbf{F}, \mathcal{Z}_{\rm vim}), \qquad \Psi_{\rm act} = \Psi_{\rm act}(\mathbf{F}, \mathcal{Z}_{\rm act}). \tag{3}$$

Here, $\mathcal{Z}_{\rm vim}$ and $\mathcal{Z}_{\rm act}$ denote collections of microstructural variables that characterize the state of the vimentin and actin networks, respectively, and whose evolution in time captures cytoskeletal remodeling, reorganization, and changes in anisotropy induced by the loading history.

Within this framework, the first Piola–Kirchhoff stress tensor is obtained from the free-energy density through the standard constitutive relation

$$\mathbf{P} = \frac{\partial \Psi}{\partial \mathbf{F}} - p\,\mathbf{F}^{-T}, \tag{4}$$

where $p$ is a Lagrange multiplier enforcing the incompressibility constraint [15]. The total stress thus results from the additive contributions of the isotropic, actin, and vimentin terms, together with the volumetric constraint associated with incompressibility.

In the proposed formulation, the different cytoskeletal and cellular constituents are assumed to experience the same macroscopic deformation, leading to an isostrain assumption underlying the additive decomposition of the free-energy density. This choice reflects the fact that, at the scale of the continuum description adopted here, the actin network, the vimentin network, and the remaining cellular components are mechanically embedded within the same cellular volume and are therefore subjected to the same overall deformation gradient. While it is well recognized that strong biochemical and mechanical interactions exist between cytoskeletal networks [16], resolving local strain heterogeneities and explicit network–network interpenetration would require a fully resolved multiscale or discrete description, which is beyond the scope of the present study. Instead, the isostrain assumption provides a tractable framework that captures the dominant features of the cell's mechanical response, while allowing each network to contribute through its own constitutive law and internal remodeling mechanisms. Importantly, this assumption does not preclude indirect coupling between networks: interactions are effectively accounted for through their concurrent response to the same deformation and through the experimentally calibrated parameters governing each contribution. In this sense, the isostrain framework represents a first-order, physically motivated approximation that balances biological realism with model interpretability and computational efficiency, and is sufficient to reproduce the key experimental trends observed in imaging and mechanical assays.

#### 3.2. *Isotropic contribution*

##### 3.2.1. *Summary of assumptions for isotropic contribution*

The isotropic contribution is introduced to represent the effective mechanical response of cellular components that are not explicitly described as organized filament networks, including the cytosolic matrix, subcellular structures lacking a persistent preferential orientation, and cytoskeletal elements such as microtubules treated in an averaged sense:

- The isotropic contribution is assumed to exhibit no dominant directional dependence under the experimental conditions considered, justifying its description as an isotropic material response.
- This term captures the baseline mechanical stiffness of the cell and its nonlinear elastic behavior under deformation.
- The isotropic contribution is assumed to be incompressible at the time scales of interest, reflecting the high water content of the cytoplasm and the limited ability of the cell to undergo volumetric changes [17].
- Incompressibility is enforced through a pressure-like Lagrange multiplier, ensuring that the isotropic response captures shape changes without artificial volume variations.
- The isotropic contribution provides an effective mechanical environment in which anisotropic cytoskeletal networks are embedded, supplying baseline resistance to deformation while actin and vimentin introduce directionality, remodeling, and load-dependent reinforcement.

##### 3.2.2. *Energetic and stress formulation*

The term $\Psi_{\text{iso}}$ is introduced to represent the effective mechanical response of all constituents that are not explicitly modeled as filamentous networks, including the cytosolic matrix and additional subcellular components (with microtubules absorbed into this effective background response). Since these constituents do not exhibit a persistent preferential orientation at the scale of the continuum description adopted here, we assume $\Psi_{\text{iso}}$ to be isotropic, i.e. dependent only on the deformation gradient $\mathbf{F}$ through the first invariant of the right Cauchy–Green tensor $\mathbf{C} = \mathbf{F}^T\mathbf{F}$

$$I_1 = \text{tr}(\mathbf{C}), \qquad \Psi_{\text{iso}} = \Psi_{\text{iso}}(I_1). \tag{5}$$

We employ a generalized incompressible isotropic potential of the form [18]

$$\Psi_{\text{iso}}(I_1) = \frac{\mu_{\text{iso}}\, 3^{\,1-\alpha_{\text{iso}}}}{2\,\alpha_{\text{iso}}}\,(I_1^{\alpha_{\text{iso}}} - 3^{\alpha_{\text{iso}}}) \tag{6}$$

where $\mu_{\text{iso}}$ is the shear modulus and $\alpha_{\text{iso}}$ controls the degree of strain-stiffening. Note that this model reduces to Neo-Hookean when $\alpha_{\text{iso}} = 1$, as in this study. The associated first Piola–Kirchhoff stress contribution follows from differentiation with respect to $\mathbf{F}$,

$$\mathbf{P}_{\text{iso}} = \frac{\partial \Psi_{\text{iso}}}{\partial \mathbf{F}} = 3^{\,1-\alpha_{\text{iso}}}\,\mu_{\text{iso}}\,I_1^{\alpha_{\text{iso}}-1}\,\mathbf{F}. \tag{7}$$

#### 3.3. *Vimentin contribution*

##### 3.3.1. *Summary of assumptions for vimentin contribution*

The constitutive description of the vimentin network is based on a minimal set of experimentally motivated assumptions aimed at capturing its dual role as a structural scaffold and a dynamically reorganizing cytoskeletal component:

- Vimentin filaments are assumed to respond primarily to deformation, and in particular to stretching [2]. Network reorganization is therefore driven by kinematic measures, with alignment emerging along the direction of maximum stretch.

- Vimentin reorganization is assumed to depend strongly on the mode of deformation. Nearly isotropic or volumetric deformation states do not promote alignment, whereas biaxial and especially uniaxial deformation progressively activate network reorganization.

- The kinetics of vimentin remodeling are assumed to be strain-dependent, with reorientation and alignment becoming significantly faster beyond a critical level of stretch [2, 4].

- The model distinguishes between reorientation of the network and development of anisotropy. Reorientation governs the evolution of the mean filament direction, whereas anisotropy quantifies the degree of alignment within the network.

- Anisotropy is assumed to increase selectively under sustained, directionally coherent tensile loading and only when the existing filament orientation is favorably aligned with the imposed deformation. Weakly directional loading does not promote anisotropy, while compressive states may reduce it [4, 2].

- The mechanical contribution of vimentin is assumed to be tension-dominated. The network transmits mechanical load primarily when stretched, whereas its contribution is strongly reduced under compression due to filament buckling [14].

*3.3.2. Energetic and stress formulation*

The contribution of the vimentin intermediate filament network is introduced to capture its structural and mechanical role as an anisotropic, load-bearing cytoskeletal component that reorganizes in response to deformation. At the scale of the present continuum description, the vimentin network is modeled as an oriented fibrous architecture whose mechanical response depends on the macroscopic deformation as well as on internal variables describing its microstructural state. Accordingly, the Helmholtz free-energy density associated with vimentin is assumed to depend on the deformation gradient $\mathbf{F}$, the mean fiber orientation $\mathbf{n}_{vim}$, and the fractional anisotropy $FA_{vim}$,

$$\Psi_{vim} = \Psi_{vim}(\mathbf{F}, \mathbf{n}_{vim}, FA_{vim}). \tag{8}$$

The anisotropic character of the vimentin network is introduced through a structural tensor of dispersion type. Experimental evidence indicates that vimentin filaments primarily sustain tensile loads, while under compression they tend to buckle and do not transmit significant stress. To capture this behavior, although the isotropic contribution is always accounted for, the anisotropic part of the structure tensor $\mathbf{G}_{vim}$ is formulated in terms of the positive part of the invariant as [19]

$$\mathbf{G}_{vim} = \kappa_{vim}\mathbf{I} + H\left\{I_{4,vim} - 1\right\}(1 - 3\kappa_{vim})\,\mathbf{n}_{vim} \otimes \mathbf{n}_{vim}, \tag{9}$$

where $H\{\cdot\}$ denotes the Heaviside function and $\kappa_{vim} \in [0, 1/3]$ is a dispersion parameter controlling the degree of alignment of the fibers. In the present formulation, $\kappa_{vim}$ is expressed as a function of the fractional anisotropy $FA_{vim}$, which provides a direct link between the continuum description and experimental measurements obtained from fluorescence microscopy. Specifically, the dispersion parameter is defined as [20]

$$\kappa_{vim} = \frac{1}{2}\frac{-6 + 4FA_{vim}^2 + 2\sqrt{3FA_{vim}^2 - 2FA_{vim}^4}}{-9 + 6FA_{vim}^2} \tag{10}$$

which ensures a smooth and monotonic transition between isotropic and highly aligned states as $FA_{vim}$ increases. Using the right Cauchy–Green tensor $\mathbf{C} = \mathbf{F}^{\mathsf{T}}\mathbf{F}$, we define the vimentin-specific invariant

$$I_{4,vim} = tr\left(\mathbf{G}_{vim}\mathbf{C}\right), \tag{11}$$

which measures the effective stretch experienced by the vimentin network along its preferred orientation while accounting for angular dispersion.
The free-energy density is chosen as [19]

$$\Psi_{vim} = \frac{\mu_{vim}}{2\beta_{vim}}\left[\exp\left(\beta_{vim}\left(I_{4,vim} - 1\right)^2\right) - 1\right], \tag{12}$$

where $\mu_{vim}$ is an effective stiffness parameter and $\beta_{vim}$ controls the degree of strain stiffening. This choice ensures that the vimentin network contributes to the mechanical response only in tension, while remaining inactive under compressive states.
The first Piola–Kirchhoff stress contribution associated with the vimentin network follows from differentiation of the free-energy density with respect to the deformation gradient,

$$\mathbf{P}_{vim} = \frac{\partial \Psi_{vim}}{\partial \mathbf{F}} = 2\mu_{vim}\left(I_{4,vim} - 1\right)\exp\left(\beta_{vim}\left(I_{4,vim} - 1\right)^2\right)\ \mathbf{F}\mathbf{G}_{vim}. \tag{13}$$

#### *3.3.3. Evolution of microstructural variables*

The mechanical response of the vimentin network is not only governed by the instantaneous deformation, but also by the evolution of its internal microstructural state. In the present model, this state is characterized by two internal variables: (i) the preferred fiber orientation $\mathbf{n}_{vim}$, and (ii) the fractional anisotropy $FA_{vim}$, which quantifies the degree of alignment of the filament network. Both variables evolve in time in response to deformation-induced cues, consistently with experimental observations of cytoskeletal remodeling under sustained loading.

**Maximum principal stretch direction and deformation mode:** Microstructural reorganization of vimentin is assumed to be driven by stretch, in agreement with the fact that filament alignment is directly associated with deformation of the cytoskeletal network. Accordingly, the target direction for reorientation is defined as the maximum principal stretch direction. Let $\mathbf{C} = \mathbf{F}^T\mathbf{F}$ be the right Cauchy–Green tensor, with spectral decomposition

$$\mathbf{C} = \sum_{i=1}^{3} \lambda_i^2\, \mathbf{n}_i \otimes \mathbf{n}_i, \tag{14}$$

where $\lambda_i$ are the principal stretches and $\mathbf{n}_i$ the associated eigenvectors. We denote by $\lambda_{max}$ the largest principal stretch and by $\mathbf{n}_{max}$ its corresponding direction.
In addition to the magnitude of deformation, the mode of deformation plays a central role in cytoskeletal remodeling. To quantify this effect, a scalar deformation-mode indicator is introduced as

$$\chi = \frac{|\ln(\lambda_{max})|}{\sum_{i=1}^{3} |\ln(\lambda_i)| + \varepsilon}, \tag{15}$$

with $\varepsilon \ll 1$ a small numerical regularization. This indicator satisfies that $\chi = 1/3$ under nearly isotropic (triaxial) deformation but it adopts a null value under the assumption of incompressibility, intermediate values under biaxial deformation, and $\chi$ reaches its maximum value under uniaxial deformation. Such a measure allows the model to distinguish between deformation states that promote directional alignment and those that do not.

**Evolution of the vimentin orientation:** Similar to previous work on fibre reorientation [21], the preferred orientation of the vimentin network is assumed to evolve toward the maximum principal stretch direction, driven by a relaxation-type process. The evolution equation is written as

$$\dot{\mathbf{n}}_{vim} = \frac{1}{\tau_{vim}} D_n(\chi) \left(\mathbf{n}_{max} - (\mathbf{n}_{max} \cdot \mathbf{n}_{vim})\, \mathbf{n}_{vim}\right), \tag{16}$$

where the term in parentheses represents the projection of $\mathbf{n}_{max}$ onto the plane orthogonal to $\mathbf{n}_{vim}$, ensuring preservation of unit norm. Reorientation of the preferred vimentin direction toward the maximum principal stretch direction is continuously performed except during disorganization regimes. Disorganization regimes are mathematically defined as temporal steps where the network transits from a highly aligned network (i.e., $FA_{vim} > 0.4$) to lower anisotropic values. The factor $D_n(\chi)$ modulates the reorientation rate depending on the deformation mode and is defined as a smooth sigmoid function,

$$D_n(\chi) = \frac{1}{1 + \exp[-a_n\,(\chi - \chi_n)]}, \tag{17}$$

with parameters $a_n > 0$ and $\chi_n$ controlling the sharpness and activation threshold of the transition. This choice reflects the hypothesis that fiber reorientation is negligible under isotropic deformation and progressively activated as the deformation becomes increasingly uniaxial.

The characteristic reorientation time $\tau_{vim}$ is assumed to depend on the level of stretch,

$$\tau_{vim} = \tau_\infty + \frac{\tau_0 - \tau_\infty}{1 + \exp[\alpha_{vim}\,(\lambda_{max} - \lambda_{crit})]}, \tag{18}$$

where $\tau_0$ and $\tau_\infty$ are limiting reorientation times at low and high stretch, respectively, $\alpha_{vim}$ controls the sensitivity to deformation, and $\lambda_{crit}$ is a critical stretch beyond which reorientation accelerates. This formulation captures the experimentally observed increase in remodeling rates once a deformation threshold is exceeded.

**Evolution of fractional anisotropy:** The fractional anisotropy $FA_{vim}$ measures the degree of alignment of the vimentin network and is allowed to evolve toward a deformation-dependent target value. Its evolution is governed by

$$\dot{FA}_{vim} = \frac{1}{\tau_{FA}} \left(FA_{vim}^{target} - FA_{vim}\right). \tag{19}$$

The target anisotropy is defined as

$$FA_{vim}^{target} = FA_{vim} + D_{FA}(\chi)\ |FA_{max} - FA_{vim}|\ \cos(2\theta)\ \left(1 - e^{-c\,\lambda_{max}}\right), \tag{20}$$

where $FA_{max} \in \{0, 1\}$ represents the limiting anisotropy state, $\theta$ is the angle between the current vimentin orientation (convected by the deformation) and the maximum principal stretch direction, $c > 0$ controls the stretch sensitivity, and $D_{FA}(\chi)$ is a deformation-mode-dependent activation function analogous to $D_n$. The angular dependence through $\cos(2\theta)$ ensures that anisotropy increases only when the vimentin network is favorably aligned with the loading direction, while misaligned configurations promote misalignment. The stretch-dependent factor $(1 - e^{-c\,\lambda_{max}})$ determines the maximum degree of network alignment as a function of the effective stretch on the system. This formulation reflects the hypothesis that sustained, directionally coherent tensile loading promotes progressive alignment of intermediate filaments, while isotropic or weakly directional deformation does not induce significant structural ordering.

#### *3.4. Actin contribution*

##### *3.4.1. Summary of assumptions for actin contribution*

The contribution of the actin cytoskeleton is modeled based on a minimal set of assumptions aimed at capturing its dual structural role and its strong coupling to cytoskeletal remodeling and vimentin integrity:

- Actin is decomposed into stress fibers and an actin cortex, treated as mechanically and kinetically distinct components [6, 13].
- Stress fibers are assumed to sustain mechanical loads primarily under tension and to exhibit an anisotropic response governed by an evolving internal reference configuration, orientation, and degree of alignment [5, 6].
- Stress-fiber mechanics depend on continuous turnover: pre-existing fibers progressively relax their elastic strain, while newly polymerized fibers are incorporated in a stress-free state [9].
- Stress-fiber polymerization is promoted by sufficient elastic stretch, whereas depolymerization is weak under baseline conditions and strongly enhanced in the absence of vimentin [7].
- Newly formed stress fibers are assumed to align with the vimentin network, while pre-existing fibers gradually reorient towards the principal tensile direction depending on the deformation mode [22].
- The actin cortex is modeled as an incompressible, isotropic layer whose mechanical contribution is active mainly under tension [13].
- Cortical actin turnover depends on in-plane deformation [13] and is strongly modulated by the presence of vimentin, reflecting its stabilizing role on cortical integrity.
- Compressive in-plane states are assumed to induce cortical buckling, leading to a reduction in effective load transmission [23, 24].

##### *3.4.2. Mass balance and phase transformation*

The mass-balance formulation is motivated by extensive experimental evidence showing that actin continuously cycles between monomeric and polymerized states, and that mechanical and biochemical cues regulate how polymerized actin is partitioned between stress fibers and the cortical network [8, 6]. Stress fibers are primarily associated with force transmission and mechanosensing, whereas the actin cortex contributes to cell shape maintenance and surface tension. By explicitly accounting for these three actin phases and their exchange dynamics, the model provides a biologically grounded description of actin remodeling that can be naturally coupled to the mechanical constitutive response of each network. Following this motivation, actin is modeled as a conserved cytoskeletal constituent that can dynamically transition between different structural organizations within the cell. In the present framework, we distinguish three actin phases: (i) monomeric or globular actin (G-actin), (ii) polymerized actin organized into stress fibers, and (iii) polymerized actin forming the actin cortex. These three phases represent the dominant functional states of actin at the cellular scale and are sufficient to capture the experimentally observed redistribution of actin under mechanical stimulation. We assume that the total amount of actin within the cell is conserved over the time scales of interest. Accordingly, the actin content is described in terms of normalized fractions as

$$\phi_G + \phi_{SF} + \phi_c = 1, \tag{21}$$

where $\phi_G$ denotes the fraction of G-actin, $\phi_{SF}$ the fraction of actin incorporated into stress fibers, and $\phi_c$ the fraction of actin forming the cortical network.

This mass balance reflects the biological fact that actin continuously polymerizes and depolymerizes, while the total actin pool remains approximately constant. Polymerization processes transfer actin from the G-actin pool to either stress fibers or the cortex, whereas depolymerization returns actin from these polymerized structures back to the G-actin pool. In the model, G-actin therefore acts as a shared reservoir that feeds both polymerized actin networks.

**Kinetics of actin redistribution:** The temporal evolution of the actin fractions is governed by phenomenological rate equations that describe polymerization and depolymerization processes. These kinetics are formulated in a minimal form that is consistent with experimental observations. The evolution of the stress-fiber fraction is given by

$$\dot{\phi}_{SF} = k_{pol}^{SF}\,\phi_G - k_{depol}^{SF}\,\phi_{SF}, \tag{22}$$

where $k_{pol}^{SF}$ and $k_{depol}^{SF}$ are effective polymerization and depolymerization rates, respectively. This equation expresses that stress fibers are formed by recruitment of monomeric actin from the G-actin pool and disassemble back into G-actin over time [8, 9].

Similarly, the evolution of the cortical actin fraction is written as

$$\dot{\phi}_c = k_{pol}^{c}\,\phi_G - k_{depol}^{c}\,\phi_c, \tag{23}$$

with $k_{pol}^{c}$ and $k_{depol}^{c}$ denoting the corresponding rates for cortical actin.

The G-actin fraction is governed by

$$\dot{\phi}_G = k_{depol}^{SF}\,\phi_{SF} + k_{depol}^{c}\,\phi_c - \left(k_{pol}^{SF} + k_{pol}^{c}\right)\phi_G. \tag{24}$$

This formulation always ensures exact mass conservation of actin and reflects the role of G-actin as a dynamically available pool that buffers actin redistribution between different structural organizations. In addition, for physical consistency, the actin phase fractions are interpreted as physical volume fractions and are therefore implicitly restricted to the interval $\phi_i \in [0, 1]$.

#### *3.4.3. Energetic and stress formulation*

The actin cytoskeleton is represented by two mechanically relevant polymerized networks: (i) an oriented stress-fiber network, and (ii) an actin cortex forming a largely isotropic shell-like reinforcement at the cell periphery. Consistent with the mass balance introduced above, the mechanical contribution of actin is scaled by the corresponding phase fractions $\phi_{SF}$ and $\phi_c$, and is written as an additive free-energy density

$$\Psi_{act} = \phi_{SF}\,\Psi_{SF} + \phi_c\,\Psi_c + \phi_G\,\Psi_G. \tag{25}$$

$$\Psi_{SF} = \Psi_{SF}(\mathbf{F}, \mathbf{n}_{SF}, FA_{SF}, \mathbf{F}_{SF})\,, \tag{26}$$

$$\Psi_c = \Psi_c(\mathbf{F}, m_{buck}), \tag{27}$$

where $\mathbf{n}_{SF}$ and $FA_{SF}$ are internal variables describing the evolving orientation and alignment of stress fibers, and $\mathbf{F}_{SF}$ is an internal remodeling/relaxation variable (introduced below in the evolution section) that controls stress relaxation of the stress-fiber network. $m_{buck}$ is an internal variable associated with actin cortex buckling. We have included a contribution associated with G-actin ($\phi_G \Psi_G$), but this term will not introduce an effective mechanical contribution.

**Stress-fiber network:** The stress-fiber response is modeled as an anisotropic fibrous contribution, with fiber dispersion controlled by the fractional anisotropy $FA_{SF}$. We define an elastic-like measure for the stress-fiber network through the internal variable $\mathbf{F}_{SF}$ (its evolution is defined later) as

$$\mathbf{C}_e = \mathbf{F}_e^{-T}\,\mathbf{C}\,\mathbf{F}_e^{-1}, \qquad \text{with} \qquad \mathbf{F} = \mathbf{F}_e\,\mathbf{F}_{SF}, \tag{28}$$

with $\mathbf{F}_e$ being the elastic part of the deformation gradient associated with stress fibers, which represents the portion of deformation effectively stored in the network, and accounts for the continuous evolution of the stress fibers configuration during remodeling. The anisotropic character of stress fibers is introduced through the structural (dispersion) tensor

$$\mathbf{G}_{SF} = \kappa_{SF}\,\mathbf{I} + H(I_{4,e} - 1)\,(1 - 3\kappa_{SF})\,\mathbf{n}_{SF} \otimes \mathbf{n}_{SF}, \tag{29}$$

where $\kappa_{SF} = \kappa(FA_{SF})$ has the same form used for vimentin and $I_{4,e} = \mathrm{tr}(\mathbf{G}_{SF}\mathbf{C}_e)$. As for vimentin, stress fibers are assumed to transmit load predominantly in tension, while their contribution is strongly reduced under compression due to filament buckling.

The stress-fiber free-energy density is chosen in exponential (strain-stiffening) form [19]

$$\Psi_{SF} = \frac{\mu_{SF}}{2\beta_{SF}}\left[\exp\Big(\beta_{SF}\,(I_{4,e} - 1)^2\Big) - 1\right], \tag{30}$$

where $\mu_{SF} > 0$ controls the network stiffness and $\beta_{SF} > 0$ controls strain-stiffening. Differentiation with respect to $\mathbf{F}$ yields the first Piola–Kirchhoff stress contribution,

$$\mathbf{P}_{SF} = \frac{\partial \Psi_{SF}}{\partial \mathbf{F}} = \left[2\mu_{SF}\,(I_{4,e} - 1)\exp\Big(\beta_{SF}\,(I_{4,e} - 1)^2\Big)\,\mathbf{F}_e\,\mathbf{G}_{SF}\right] : \frac{\partial \mathbf{F}_e}{\partial \mathbf{F}}. \tag{31}$$

**Actin cortex:** The actin cortex is a thin, highly dynamic network of polymerized actin located immediately beneath the cell membrane. Structurally, it forms a peripheral shell that plays a central role in regulating cell shape, surface tension, and the mechanical response of the cell boundary. In indentation-based mechanical assays, such as those considered in this work, the cortex is therefore expected to contribute significantly to the measured response, as it is the first cytoskeletal structure engaged by the applied load. Although the actin cortex is spatially localized near the cell membrane, the present model adopts a homogenized continuum description of the cell. Within this framework, the mechanical effect of the cortex is introduced as an effective constitutive contribution that captures its dominant influence on the macroscopic stress response under deformation, without explicitly resolving its geometric thickness or spatial distribution. This approach is justified by the separation of scales between the cortical thickness and the characteristic dimensions of the cell, and by the focus on global force–deformation behavior rather than on local stress distributions within the cortex itself.

Accordingly, the cortex is modeled as an effective isotropic, strain-stiffening network whose contribution

is modulated by the amount of polymerized actin available in the cortical layer. The cortex is assumed to transmit mechanical loads predominantly in tension [13]. Under compressive deformation, thin filamentous shells such as the actin cortex are prone to wrinkling or buckling [23, 24], which strongly reduces their ability to sustain compressive stresses. To account for this asymmetry, the model introduces a scalar buckling reduction factor $m_{buck} \in [0, 1]$ that modulates the effective mechanical contribution of the cortex. This factor remains close to unity when the cortex is stretched, corresponding to efficient load transmission, and progressively decreases as the deformation state becomes compressive, reflecting the onset of buckling and loss of effective stiffness.
Within this homogenized description, the energetic contribution of the actin cortex is written as an incompressible isotropic strain-stiffening potential [18] scaled by $m_{buck}$,

$$\Psi_c = \frac{3^{\,1-\alpha_c}\,\mu_c}{2\alpha_c}\, m_{buck}\, \left(I_1^{\alpha_c} - 3^{\alpha_c}\right), \tag{32}$$

where $\mu_c > 0$ is a stiffness-like parameter and $\alpha_c$ controls the degree of strain stiffening. The corresponding first Piola–Kirchhoff stress contribution is obtained by differentiation as

$$\mathbf{P}_c = 3^{\,1-\alpha_c}\,\mu_c\, m_{buck}\, I_1^{\alpha_c-1}\,\mathbf{F}. \tag{33}$$

Moreover, to capture the fact that the actin cortex transmits mechanical load primarily in tension, we introduce a scalar buckling reduction factor $m_{buck}$, which modulates the effective cortical contribution to the stress response. In the absence of buckling, the cortex behaves as an efficient load-bearing network and $m_{buck} \approx 1$, whereas under sufficiently strong compression buckling or wrinkling strongly reduces its ability to transmit stress and $m_{buck}$ approaches zero.
This formulation should be interpreted as an effective representation of the mechanical role of the actin cortex at the cell scale. Rather than resolving the cortex as a separate geometric layer, the model captures its influence through a constitutive term that becomes particularly relevant under tensile deformation of the cell boundary, as occurs during indentation. In this way, the approach balances biological realism with computational tractability, while remaining consistent with experimental observations of cortical mechanics.

*3.4.4. Evolution of microstructural variables*

**Stress-fiber network:** The stress-fiber network is endowed with an internal reference configuration $\mathbf{F}_{SF}$ that evolves due to remodeling and turnover as

$$\mathbf{F}_{SF} = \frac{\phi_{SF}^n}{\phi_{SF}}\left[\mathbf{F}_{SF}^n + \frac{\Delta t}{\tau_{act}}\left(\mathbf{F} - \mathbf{F}_{SF}^n\right)\right] + \frac{\phi_{SF} - \phi_{SF}^n}{\phi_{SF}}\,\mathbf{F}. \tag{34}$$

The parameter $\tau_{act}$ modulates the progressive loss of elastic memory of pre-existing fibers through remodeling, while the turnover term reflects that newly polymerized actin is incorporated in a stress-free state. Together, these mechanisms ensure that stress-fiber mechanics are governed not only by the applied deformation but also by the history of cytoskeletal remodeling, allowing for stress relaxation and adaptation under sustained loading.
The polymerization and depolymerization rates entering the stress-fiber mass balance are assumed to be regulated by the elastic stretch effectively experienced by the actin network along its dominant directions (derive from $I_{4,e}$). Only tensile contributions are assumed to promote actin assembly [7], which is enforced

using Macaulay brackets in the definition of the polymerization rate is defined as

$$k_{pol}^{SF} = k_{pol,0}^{SF} \left\langle \frac{\sqrt{\langle I_{4,e} \rangle}}{\lambda_{thres}} - 1 \right\rangle , \tag{35}$$

where $\lambda_{thres} > 1$ denotes a stretch threshold beyond which stress fibers actively polymerize, and $k_{pol,0}^{SF}$ is a reference polymerization rate. This choice reflects the experimentally observed activation of stress-fiber assembly under sufficiently large tensile strains, while suppressing polymerization under compressive or weakly tensile states.

Motivated by the experimental results provided herein, depolymerization is taken to be weak under baseline conditions but strongly enhanced in the absence of vimentin. To account for this effect, we introduce a binary indicator variable $\delta_{vim}$, defined as

$$\delta_{vim} = \begin{cases} 1, & \text{if vimentin present,} \\ 0, & \text{if vimentin depleted.} \end{cases} \tag{36}$$

The depolymerization rate is then written as

$$k_{dep}^{SF} = k_{dep,0}^{SF} \left[ 1 + \frac{1 - \delta_{vim}}{k_{dep,0}^{SF}} \left( \frac{\langle \sqrt{\langle I_{4,e} \rangle} - \lambda_{thres} \rangle}{\lambda_{thres}} \right)^2 \right] , \tag{37}$$

with $k_{dep,0}^{SF}$ a reference depolymerization rate. This formulation captures the stabilizing role of the vimentin network on stress fibers, whereby loss of intermediate filaments leads to accelerated actin disassembly under deformation.

The evolution of the stress-fiber anisotropy parameter is defined as

$$\dot{FA}_{SF} = \frac{1}{\tau_{act}^{int}} \frac{\phi_{SF} - \phi_{SF}^{n}}{\phi_{SF}} \left( FA_{vim} - FA_{SF} \right) + \frac{1}{\tau_{act}} \frac{\phi_{SF}^{n}}{\phi_{SF}} D_{FA}(\chi) \left| FA_{max} - FA_{SF} \right| \cos(2\theta) \left( 1 - e^{-0.15\, \lambda_{max}} \right) , \tag{38}$$

where $\tau_{act}^{int}$ denotes a characteristic time, and $\theta = \arccos \left( \frac{\mathbf{n}_{e,max} \cdot \mathbf{n}_{SF}}{\|\mathbf{n}_{e,max}\| \, \|\mathbf{n}_{SF}\|} \right)$ is the angle between the current stress-fiber direction $\mathbf{n}_{SF}$ and the principal elastic stretch direction $\mathbf{n}_{e,max}$. The first term accounts for the contribution of newly polymerized stress fibers. The ratio $(\phi_{SF} - \phi_{SF}^{n})/\phi_{SF}$ represents the fraction of newly incorporated fibres within the current stress-fiber population. These fibres are assumed to be integrated following the pre-existing cytoskeletal network, here dominated by the organization of the vimentin network. This hypothesis is motivated by experimental evidence showing that vimentin provides a mechanical and structural template that guides actin assembly and stabilizes filament alignment. The second term describes the gradual reorientation of pre-existing stress fibers. The ratio $\phi_{SF}^{n}/\phi_{SF}$ represents the fraction of fibres that persist from the previous configuration and therefore undergo remodeling rather than de novo assembly. The factor $\cos(2\theta)$ ensures that reorientation is promoted when fibers are sufficiently aligned with the principal tensile direction and suppressed when they are transverse to it. The target anisotropy $FA_{max}$ acts as a binary indicator of whether the local mechanical environment favors the development of anisotropy or its reduction, thereby preventing unphysical growth of alignment under unfavorable loading configurations. The scalar function $D_{FA}(\chi)$ modulates the efficiency of stress-fibre reorientation depending on the deformation mode $\chi$, promoting alignment under predominantly uniaxial stretching and suppressing it under multiaxial or isotropic deformation states. Finally, the multiplicative factor $(1 - e^{-0.15\, \lambda_{max}})$

accounts for the requirement of a finite deformation magnitude to activate cytoskeletal remodeling: small elastic stretches produce negligible reorganization, whereas increasing stretch progressively enhances the rate of stress-fibre reorientation.

The evolution of the dominant stress-fiber direction is given by

$$\dot{\mathbf{n}}_{SF} = H_n \left[ \frac{1}{\tau_{act}^{int}} \frac{\phi_{SF} - \phi_{SF}^n}{\phi_{SF}} \mathbf{n}_{vim} + \frac{1}{\tau_{act}} \frac{\phi_{SF}^n}{\phi_{SF}} D_n(\chi) \left( \mathbf{n}_{e,max} - (\mathbf{n}_{e,max} \cdot \mathbf{n}_{SF}) \mathbf{n}_{SF} \right) \right], \qquad \|\mathbf{n}_{SF}\| = 1, \tag{39}$$

where $\mathbf{n}_{vim}$ denotes the dominant vimentin direction and $D_n(\chi)$ modulates the reorientation rate depending on the deformation mode $\chi$. The prefactor $H_n$ implements the remodeling activation criterion used in the numerical model, so that significant reorientation is restricted to configurations that either (i) actively promote anisotropy growth or (ii) correspond to weakly organized stress fibres ($FA_{SF} < 0.4$), in which case the network is assumed to remain sufficiently labile to reorient. The first term inside brackets accounts for the incorporation of newly polymerized fibers: the ratio $(\phi_{SF} - \phi_{SF}^n)/\phi_{SF}$ represents the fraction of new fibres within the current stress-fibre population, which are assumed to be seeded along the vimentin scaffold $\mathbf{n}_{vim}$. The second term describes the rotation of pre-existing fibres: the ratio $\phi_{SF}^n/\phi_{SF}$ weights the contribution of the old population, which reorients towards the principal elastic stretch direction. The projection of the direction ensures pure rotation without spurious changes in fiber magnitude; accordingly, the evolution is complemented by the normalization constraint $\|\mathbf{n}_{SF}\| = 1$, consistent with enforcing a unit director.

**Actin cortex:** For the actin cortex, polymerization and depolymerization rates are assumed to depend on in-plane deformation [13] and on the presence of vimentin. Polymerization is enhanced by in-plane areal stretch and is defined as

$$k_{pol}^c = k_0^c \left( 1 + \left\langle \sqrt{\det \mathbf{F}_{xy}} - 1 \right\rangle \right), \tag{40}$$

where $k_0^c$ is a reference rate, reflecting the tendency of the cortex to reinforce under membrane stretching. Depolymerization is modulated by both stretch anisotropy and vimentin integrity,

$$k_{dep}^c = k_0^c \left( 1 + \eta_{vim}(1 - \delta_{vim}) \right) \left[ 1 - \frac{1}{2} \langle \lambda_{max} - 1 \rangle + \langle 1 - \lambda_{min} \rangle \right], \tag{41}$$

where $\lambda_{\max}$ and $\lambda_{\min}$ are the principal stretches of the in-plane $XY$ sub-tensor of the deformation gradient, corresponding to the plane of the actin cortex. This formulation captures the suppression of cortical turnover under in-plane expansion: uniaxial stretch moderately reduces depolymerization, whereas biaxial expansion, which increases both principal stretches, produces a stronger inhibition and thus greater cortical stabilization. The parameter $\gamma_{\rm vim} > 0$ controls the magnitude of the increase in cortical actin depolymerization induced by vimentin depletion. This formulation reflects experimental observations indicating that loss of the vimentin network leads to destabilization of the actin cortex, whereas its presence confers mechanical robustness and resistance to turnover. By construction, this term acts solely as a modulatory factor and does not alter the baseline turnover kinetics when vimentin is present.

Regarding the evolution of $m_{buck}$, experimental evidence indicates that cortical remodeling and mechanical stability are strongly coupled with the presence and organization of the vimentin network. Fluorescence imaging and perturbation studies show that vimentin provides a structural scaffold that supports actin reorganization and stabilizes peripheral cytoskeletal structures. Motivated by this observation, the compressive state of the cortex is evaluated along the current in-plane vimentin direction, which serves as a robust proxy for the dominant structural axis of the cytoskeleton in the region probed by indentation as

$$E_{vim} = \frac{1}{2}\left(\lambda_{vim}^2 - 1\right), \qquad \lambda_{vim} = \sqrt{\mathbf{n}_{vim}^T \mathbf{C}\, \mathbf{n}_{vim}}, \tag{42}$$

where compression corresponds to $E_{vim} < 0$. We introduce a nonnegative compressive measure $s = \max(-E_{vim}, 0)$, and a critical compression threshold $s_{crit} = \frac{1}{2}\left(1 - \lambda_{crit}^2\right), \qquad \lambda_{crit} < 1$, beyond which buckling is assumed to be fully developed. Rather than prescribing the buckling state instantaneously, the model treats $m_{buck}$ as an internal variable that evolves in time toward a deformation-dependent target state $m_{buck}^*(s)$. This target state represents the buckling configuration favored by the current deformation and is defined as

$$m_{buck}^*(s) = \begin{cases} 1, & s = 0, \\ 1 - \dfrac{\exp(b\, s/s_{crit}) - 1}{\exp(b) - 1}, & 0 < s < s_{crit}, \\ 0, & s \geq s_{crit}. \end{cases} \tag{43}$$

where $b > 0$ controls the sharpness of the transition from a fully load-bearing to a buckled, load-inefficient state. The actual buckling reduction factor $m_{buck}$ relaxes toward this target state according to the flow rule

$$\dot{m}_{buck} = \frac{1}{\tau_{buck}}\left(m_{buck}^* - m_{buck}\right), \qquad m_{buck}(0) = 1, \tag{44}$$

where $\tau_{buck}$ is a characteristic time scale associated with cortical buckling/wrinkling and recovery. This formulation reflects the fact that buckling and unbuckling of the actin cortex are dynamic processes governed by cytoskeletal remodeling and membrane–cytoskeleton interactions, rather than instantaneous mechanical switches.


### Acknowledgements

The authors acknowledge support from the European Research Council (ERC) under the European Union's Horizon 2020 Research and Innovation Programme (Grant agreement No. 947723, project: 4D-BIOMAP, and Grant agreement No. 101247449, project: MAGMATED) and from the European Innovation Council (EIC) under the Horizon Europe Programme (Grant agreement No. 101284471, project: NEOMAG). The authors acknowledge support from from MICIU/AEI/10.13039/501100011033 under Grant number PID2023-149255NB-I00 and PID2023-152631OB-I00 and from FEDER, UE. CGC acknowledges support from the Ministerio de Universidades, Spain (FPU20/01459). This work was also supported by La Ligue contre le cancer (EL2023 - DN/IP/IQ – 17691), Worldwide Cancer Research (WCR 23-0156 ), INSERM PCSI (N° 22CP073-00) and Institut Pasteur (PTR-548-22).


### References


[1] P. A. Janmey, U. Euteneuer, P. Traub, M. Schliwa, Viscoelastic properties of vimentin compared with other filamentous biopolymer networks., The Journal of cell biology 113 (1) (1991) 155–160.

[2] A. V. Schepers, C. Lorenz, P. Nietmann, A. Janshoff, S. Klumpp, S. Köster, Multiscale mechanics and temporal evolution of vimentin intermediate filament networks, Proceedings of the National Academy of Sciences 118 (27) (2021) e2102026118.

[3] J. Stricker, P. Maddox, E. Salmon, H. P. Erickson, Rapid assembly dynamics of the escherichia coli ftsz-ring demonstrated by fluorescence recovery after photobleaching, Proceedings of the National Academy of Sciences 99 (5) (2002) 3171–3175.

[4] Y.-C. Lin, N. Y. Yao, C. P. Broedersz, H. Herrmann, F. C. MacKintosh, D. A. Weitz, Origins of elasticity in intermediate filament networks, Physical review letters 104 (5) (2010) 058101.

[5] N. Gavara, R. S. Chadwick, Relationship between cell stiffness and stress fiber amount, assessed by simultaneous atomic force microscopy and live-cell fluorescence imaging, Biomechanics and Modeling in Mechanobiology 15 (3) (2016) 511–523. doi:10.1007/s10237-015-0706-9.
URL http://link.springer.com/10.1007/s10237-015-0706-9

[6] A. F. Pegoraro, P. Janmey, D. A. Weitz, Mechanical Properties of the Cytoskeleton and Cells, Cold Spring Harbor Perspectives in Biology 9 (11) (2017) a022038. doi:10.1101/cshperspect.a022038.
URL http://cshperspectives.cshlp.org/lookup/doi/10.1101/cshperspect.a022038

[7] A. Roshanzadeh, T. T. Nguyen, K. D. Nguyen, D.-S. Kim, B.-K. Lee, D.-W. Lee, E.-S. Kim, Mechanoadaptive organization of stress fiber subtypes in epithelial cells under cyclic stretches and stretch release, Scientific reports 10 (1) (2020) 18684.

[8] T. D. Pollard, Rate constants for the reactions of atp-and adp-actin with the ends of actin filaments., The Journal of cell biology 103 (6) (1986) 2747–2754.

[9] M. A. Smith, E. Blankman, M. L. Gardel, L. Luettjohann, C. M. Waterman, M. C. Beckerle, A zyxin-mediated mechanism for actin stress fiber maintenance and repair, Developmental cell 19 (3) (2010) 365–376.

[10] Š. Z. Jokhadar, J. Iturri, J. L. Toca-Herrera, J. Derganc, Cell stiffness under small and large deformations measured by optical tweezers and atomic force microscopy: effects of actin disruptors ck-869 and jasplakinolide, Journal of Physics D: Applied Physics 54 (12) (2021) 124001.

[11] T. Wakatsuki, B. Schwab, N. C. Thompson, E. L. Elson, Effects of cytochalasin D and latrunculin B on mechanical properties of cells, Journal of Cell Science 114 (5) (2001) 1025–1036. doi:10.1242/jcs.114.5.1025.
URL https://journals.biologists.com/jcs/article/114/5/1025/979/Effects-of-cytochalasin-D-and-latrunculin-B-on

[12] M. L. Gardel, F. Nakamura, J. H. Hartwig, J. C. Crocker, T. P. Stossel, D. A. Weitz, Prestressed f-actin networks cross-linked by hinged filamins replicate mechanical properties of cells, Proceedings of the National Academy of Sciences 103 (6) (2006) 1762–1767.

[13] G. Salbreux, G. Charras, E. Paluch, Actin cortex mechanics and cellular morphogenesis, Trends in cell biology 22 (10) (2012) 536–545.

[14] Y. Ideses, V. Erukhimovitch, R. Brand, D. Jourdain, J. S. Hernandez, U. Gabinet, S. A. Safran, K. Kruse, A. Bernheim-Groswasser, Spontaneous buckling of contractile poroelastic actomyosin sheets, Nature communications 9 (1) (2018) 2461.

[15] G. A. Holzapfel, Nonlinear solid mechanics: A continuum approach for engineering science, Meccanica 37 (4) (2002) 489–490. doi:10.1023/A:1020843529530.
URL https://doi.org/10.1023/A:1020843529530

[16] F. Huber, A. Boire, M. P. López, G. H. Koenderink, Cytoskeletal crosstalk: when three different personalities team up, Current Opinion in Cell Biology 32 (2015) 39–47, cell architecture. doi:https://doi.org/10.1016/j.ceb.2014.10.005.

[17] E. Moeendarbary, L. Valon, M. Fritzsche, A. R. Harris, D. A. Moulding, A. J. Thrasher, E. Stride, L. Mahadevan, G. T. Charras, The cytoplasm of living cells behaves as a poroelastic material, Nature Materials 12 (3) (2013) 253–261. doi:10.1038/nmat3517.
URL https://doi.org/10.1038/nmat3517

[18] O. Lopez-Pamies, A new i1-based hyperelastic model for rubber elastic materials, Comptes Rendus Mécanique 338 (1) (2010) 3–11. doi:https://doi.org/10.1016/j.crme.2009.12.007.
URL https://www.sciencedirect.com/science/article/pii/S1631072109002113

[19] T. C. Gasser, R. W. Ogden, G. A. Holzapfel, Hyperelastic modelling of arterial layers with distributed collagen fibre orientations, Journal of The Royal Society Interface 3 (6) (2006) 15–35. arXiv:https://royalsocietypublishing.org/rsif/article-pdf/3/6/15/480363/rsif.2005.0073.pdf, doi:10.1098/rsif.2005.0073.
URL https://doi.org/10.1098/rsif.2005.0073

[20] D. Garcia-Gonzalez, N. S. Race, N. L. Voets, D. R. Jenkins, S. N. Sotiropoulos, G. Acosta, M. Cruz-Haces, J. Tang, R. Shi, A. Jérusalem, Cognition based btbi mechanistic criteria; a tool for preventive and therapeutic innovations, Scientific Reports 8 (1) (2018) 10273. doi:10.1038/s41598-018-28271-7.

[21] A. V. Melnik, A. Goriely, Dynamic fiber reorientation in a fiber-reinforced hyperelastic material, Mathematics and Mechanics of Solids 18 (6) (2013) 634–648. arXiv:https://doi.org/10.1177/1081286513485773, doi:10.1177/1081286513485773.

[22] S. Seetharaman, S. Etienne-Manneville, Cytoskeletal crosstalk in cell migration, Trends in cell biology 30 (9) (2020) 720–735.

[23] M. P. Murrell, M. L. Gardel, F-actin buckling coordinates contractility and severing in a biomimetic actomyosin cortex, Proceedings of the National Academy of Sciences 109 (51) (2012) 20820–20825.

[24] R. Kusters, C. Simon, R. Lopes Dos Santos, V. Caorsi, S. Wu, J.-F. Joanny, P. Sens, C. Sykes, Actin shells control buckling and wrinkling of biomembranes, Soft Matter 15 (2019) 9647–9653.
